# Redistribution of sidechain-sidechain interactions govern ligand-specific binding affinity changes in missense Shank1 PDZ variants

**DOI:** 10.1101/2025.07.03.662971

**Authors:** Anna Sánta, Zsófia E. Kálmán, Eszter Nagy-Kanta, Borbála Jakab, Zoltán Gáspári, Bálint Péterfia

**Affiliations:** Faculty of Information Technology and Bionics, Pázmány Péter Catholic University, Budapest, Hungary

**Keywords:** : Postsynaptic density, PDZ domain, peptide binding, alternate interactions, missense mutation

## Abstract

Shank proteins represent a family of abundant scaffolds in the postsynaptic density. Their dysfunctions had been identified as possible causes behind autism spectrum disorders and various types of cancer. The remarkably promiscuous PDZ domain of the Shank family is highly conserved through isoforms, and contains a unique dynamic segment, the β2-β3 loop, located close to the binding site. We used the Shank1 PDZ as a model system to analyze the perturbing effects of five disease-associated missense mutations on the binding of different partner peptides. Using experimental methods and molecular dynamics simulations, we characterized the interactions in detail, focusing on their dynamic aspect. While the investigated variants in general weaken most interactions, the R736Q variant, unique in having increased thermal stability, also binds the GKAP peptide with higher affinity than the wild type. Overall, our results show that the perturbing effect of mutations is highly partner-specific and depends on the dynamic rearrangements of both uniformly occurring and ligand-specific residue-residue interactions.

## Introduction

Shank1, a member of the Shank family consisting of three proteins, is one of the major scaffolding proteins at the postsynaptic density (PSD) of excitatory synapses. All three members of the protein family are composed of ankyrin-repeat-, SH3-, PDZ- and SAM domains as well as a long, proline-rich disordered region, with high sequence identity in their conserved domains. (Sheng & Kim, 2000) **(*Fig. 1A*)** Due to these protein interacting domains and regions, Shank1 is able to form supramolecular complexes with a great number of other proteins. Shank1 has been associated with neurological disorders such as autism spectrum disorder, schizophrenia, mental disability and recently, its role in tumor progression has also been revealed, which are detailed below. The PDZ domain of Shank1 is a remarkably promiscuous Class I PDZ domain, with a unique β2-β3 loop region that is only found in the Shank family. Otherwise, its secondary structure composition is highly conserved, as well as its regions involved in partner binding. Structural features of its bound state are well documented via experimentally determined structures, with several binding partners, especially GKAP. (Ali et al., 2021; Im et al., 2003; Im et al., 2010; Lee et al., 2011; Ponna et al., 2018; Zeng et al., 2016) **(*Fig. 1B-D*)**

**Figure 1.**
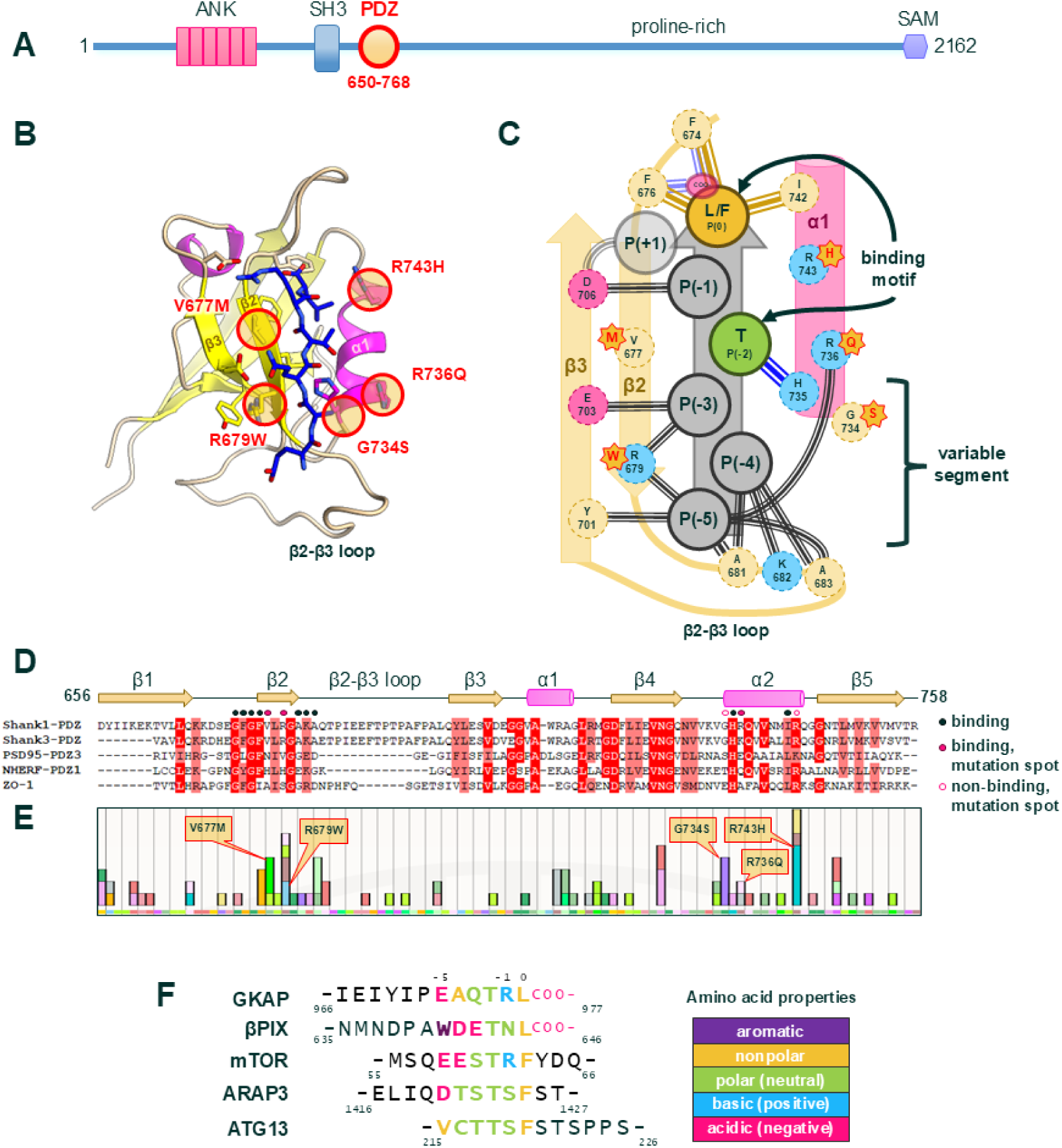
The Shank1-PDZ and its binding modes. **(A)** Domain composition of the Shank1 protein, with the PDZ domain and the span of our construct highlighted in red (the canonical PDZ spans from 663-757). For a list of constructs, refer to the Supplementary Tables. **(B)** Structure model of the Shank1-PDZ, bound to the C-terminal of GKAP (blue), based on PDB ID 1Q3P. Secondary structure elements are colored yellow for beta-strands and magenta for alpha-helices. Locations of point mutations are circled red and labelled. **(C)** Schematic representation of the Shank1-PDZ ligand binding pocket, with variable sidechain bonds depicted with black triple lines. Note that not all of the variable bonds are present in all peptide binding interactions. The nonpolar interactions of P(0) and the hydrogen bond of P(-2) are highly conserved. The carboxylate interaction is only relevant in terminal motifs, while P(+1) is only found in internal motifs, therefore they are depicted in lower opacity to reflect that. **(D)** Secondary structure composition of the Shank1-PDZ, with multiple sequence alignment comparing it to other PDZ domains. Identities and similarities within 80% are highlighted with red and light red respectively. Note that the β2-β3 loop segment is only found in the Shank PDZ domains. Positions that participate in binding according to currently known experimental structures of the Shank1-PDZ and/or are affected by mutations are marked, with legend included. **(E)** Histogram of mutations from the COSMIC database, overlaid onto the Shank1-PDZ sequence. The mutations analyzed in this study are labelled. **(F)** Partner peptides analyzed in this study, with binding motif amino acids colored according to their properties.

Regarding neural disorders, unlike Shank2 or -3, Shank1 is known to be expressed predominantly in the brain, in a distinctly different pattern with highest abundance in the cortex, cerebellum and hippocampus. Shank1 knock-out mice exhibit a complex behavioral phenotype and decreased expression of GKAP and Homer proteins, which are well-known direct interaction partners of Shank1. (Sungur et al., 2014) The C-terminus of GKAP is bound to the PDZ domain of Shank1, while Homer interacts with the proline-rich segment. (Tong et al., 2014; Tu et al., 1998) So far, Shank1 have been regarded as a low risk ASD gene probably due to the obscure, mild phenotype of KO mice - unlike Shank3, which is associated with severe autism-like symptoms in KO mouse models, and is the cause of Phelan-McDermid syndrome in humans. (Sungur et al. 2014; Alexandrov et al., 2017) Lack of Shank3 results in downregulation of the Akt/mTORC1 pathway, resulting in decreased protein synthesis and neuronal dysfunction. (Burbach, 2016) However, recently, knock-in mice bearing the R882H missense mutation (localized downstream to the PDZ domain) have been reported to display core ASD symptoms with social disability and repetitive behaviors due to downregulation of mGluR1-Homer-IP3R1-calcium signaling. (Qin et al., 2022) This finding underlines that missense mutations are promising tools to explore the role that Shank proteins and their domains play in neural disorders, and suggests that loss-of-function PDZ domain missense mutations could be used to elucidate the role of the GKAP-Shank1 interaction (and other PDZ interactions) in such disorders. Only one missense mutation of the Shank1 PDZ domain (R736Q) is described in relation to ASD, but its pathogenicity is uncertain. (Leblond et al., 2014)

Beyond their synaptic functions, Shank proteins have been recognized to play a role in several malignancies with altered expression and survival correlation. (Chang et al., 2022; Lilja et al., 2017; Xu et al., 2021) Shank expression or DNA methylation are independent prognostic markers in colon cancer and leukemia. (Loi et al., 2019; Wang et al., 2020) In addition, the expression of Shank proteins regulates the fate and development of stem cells through different signaling pathways. (Liu et al., 2022) The tumor promoting effects of Shank1 were found to be mediated by the AKT/mTOR signaling pathway in colon cancer cell lines, and by MDM2 in non-small lung cancer cells. (Chen et al., 2022; Wang et al., 2020) Nuclear translocation of MDM2 is known to be triggered by the AKT/mTOR pathway, which suggests a common pathway of Shank1 in these cell lines. There is no direct evidence as to which domain of Shank1 may play a role in activating the Akt/mTOR pathway.

Deletion of the PDZ domain of Shank2 (nearly identical in sequence to the Shank1-PDZ) causes the protein to lose its oncogenicity via Hippo signaling, but whether Shank1 or Shank3 also participate in this pathway is unknown. The aforementioned Shank2 pathway is driven by the interaction with ARHGEF7 (βPIX), also a known interaction partner of the Shank1-PDZ. (Xu et al., 2021)

As of now, no mouse model is known that was created with the aim of modeling oncogenic behavior of Shank1 specifically. Xenograft experiments with overexpressed Shank1 and Shank2 showed tumor promoting effects. (Chen et al., 2022; Xu et al., 2021) No specific point mutations of Shank1 are known to be directly associated with cancer. However, several point mutations found in tumor samples are documented in the COSMIC database. (Sondka et al., 2024) **(*Fig. 1E*)**

To avoid investing significant effort in generating transgenic mice with neutral benign point mutations, it is crucial to experimentally characterize potential variants in vitro. This study examines the effects of one ASD-associated and four somatic missense variants of the Shank1-PDZ domain on partner binding capacity and structure in vitro, starting with GKAP as the reference model, and expanding to four other partners with possible clinical significance. From these additional partners, βPIX bears a C-terminal binding motif, whereas ARAP3, ATG13 and mTOR exhibit internal binding sequences. ***(Fig. 1F, Table S1A)*** We assess the thermal stability and binding properties of the Shank1-PDZ variants and use molecular dynamics simulations to aid the interpretation of our findings.

## Results

### The most frequent somatic and ASD-associated mutations affect positions around the ligand-binding pocket

Somatic mutations for this study were picked from the COSMIC database, originally with the intent of preferring binding-site mutations for experimental investigation. Crystal structures 1Q3P, 3QJN (and 3L4F), 7A9B, 6YWZ and 8S1R (which display Shank1-PDZ in complex with GKAP, βPIX, ARAP3, a modified synthetic construct based on GKAP, and a synthetic peptide with sequence similarity to mTOR, respectively) were used as references to analyze the context of each mutation. (Ali et al., 2021; Hegedüs et al., 2021; Im et al., 2003; Im et al., 2003; Lee et al., 2011; Li et al., 2024) First, it was noted that the binding site is indeed a hot spot for mutations, and the most common missense mutation spots were located on either the β2 strand or the α2 helix forming its main parts. Four mutations fitting this criteria were picked. The one mutation picked from a study on ASD is also located in this area and has two entries in COSMIC. **(*Fig. 1E*)** (Leblond et al., 2014; Sondka et al., 2024)

The arginine sidechain affected by the most common mutation (R743H) according to the COSMIC database, is described to interact with ligands only in a subset of available experimental structures. In its close proximity, the C-terminal residue P(0) of the ligand is buried within a hydrophobic pocket formed by F674, F676 and I742, which is partially covered by R743, which simultaneously also might form a salt bridge with the peptide backbone at P(-1), however this was only observed in some models. (Ali et al., 2021; Lee et al., 2011) We predict that a mutation to histidine, a sidechain with less charge and volume, could unfavourably affect these interactions. Another possibility is a transient interaction which was seen only in a study on a complex with a synthetic GKAP-based ligand (6YWZ), so far. This interaction involves an acylhydrazone fragment component at the same position as P(-5), for which, R743 competes with R736 and R679. (Hegedüs et al., 2021)

Residues V677 and R679 are both on the β2 strand, and both bear side chains that were either documented to interact with ligands, or are in close proximity to a known interaction. V677 in the WT is surrounded by P(-1) and P(-3) of the ligand, both of which are possibly part of the binding interface in most of the complexes. Ligand positions P(-3) and P(-5) form various interactions with R679 in all known structures, including the aforementioned dynamic, transient interaction with the synthetic, GKAP-based ligand. (Ali et al., 2021; Hegedüs et al., 2021; Im et al., 2003; Lee et al., 2011; Li et al., 2024; Ponna et al., 2018) These two mutations are the closest to the β2-β3 loop in the sequence, a highly mobile segment unique to and conserved in Shank PDZ domains. (Lee et al., 2011)

The G734S mutation is located right before the α2 helix. It is in similarly close spatial proximity to the β2-β3 loop as position 679, and is next to residue H735 which H-bonds to the conserved Thr P(-2) of the binding motif. (Im et al., 2003)

R736Q is unique among the mutations in this study as the only one that was known to be associated with ASD, although entries are also documented in the COSMIC database. (Leblond et al., 2014; Sondka et al., 2024) The sidechain is on the surface and there is no data supporting that it contributes to any binding interactions with known partner proteins, although it was observed to interact with the aforementioned synthetic ligand based on the GKAP peptide, along with R743 and R679. (Hegedüs et al., 2021)

### Missense mutations affect the stability but not the structure of Shank1-PDZ

ECD spectra recorded at 30 °C suggested a well folded, globular structure with beta sheet predominance that is in accordance with the available X-ray structure of the domain (PDB ID 1Q3O (Im et al., 2003)) and PDZ domains in general. **(*Fig. 2A-C*)** No significant changes in the ECD spectrum at 30 °C were observed in any of the mutated structures. **(*Fig. 2A*)** However, remarkable differences were detected in their thermal stability both by thermal shift assays (TSA) and CD scan measurements. Melting points observed by TSA were found to be 59 °C for the WT, and 56.9, 52.0, 58.2, 61.6 and 58.8 °C for the V677M, R679W, G734S, R736G and R743H variants, respectively. These are in good accordance with the CD scan measurements. ***(Fig. 2B, 2D)*** Both the decrease at 200 nm and the increase at 210 nm in the CD signals upon heating suggests the increase of the proportions of random coil and β-strand structural elements in the domain. This is reinforced by analysis with BeStSel (Micsonai et al., 2025), that resulted in an estimation of decrease in α-helical structures and an increase in β-strands, turn and other elements, suggesting unfolding and aggregation.

**Figure 2.**
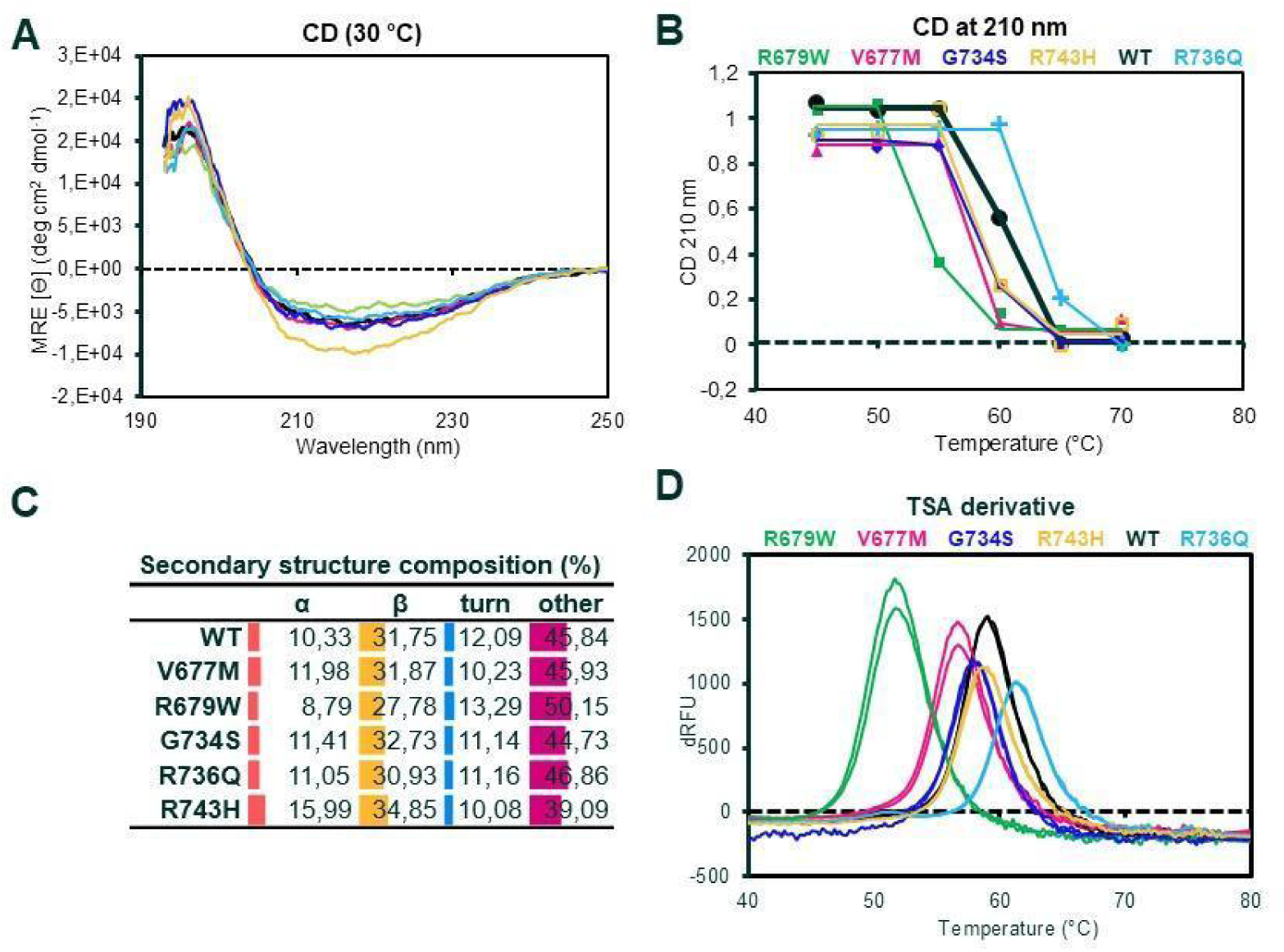
Structural stability of the Shank1-PDZ missense variants. **(A)** CD values at 30°C. Note that the curves are nearly identical. **(B)** Normalized CD curves of the PDZ isoforms at 210 nm. Note that the inflection points of the curves align with the peaks in the TSA data below at D, indicating that the results from the two methods are consistent. **(C)** Secondary structure compositions calculated from CD spectra at 30 °C with BeStSel. (Micsonai et al., 2025) „Helix1” and „Helix2” categories were added up into „α”, „Anti1”, „Anti2”, „Anti3”, and Para into „β”. **(D)** TSA results of the different Shank1-PDZ isoforms. Peaks indicate melting points.

### Effect of the mutations on domain stability and GKAP binding move in parallel

In order to examine the functional effect of the mutations, interaction of the wild type and missense variant PDZ domains with the GKAP C terminal GH1 construct was tested by a simple pull-down assay. ***(Fig. 3A, Fig. S1)*** The R679W variant was pulled down in significantly less amount by GH1 bait than the other variants, for which these experiments showed little difference relative to the wild type. This result was corroborated by fluorescence polarization spectroscopy measurements. Using a 12 amino acid long, FITC labeled GKAP C-terminus peptide ligand (CT12), the dissociation constants (K_d_) of WT, V677M, R679W, G734S, R736Q and R743H variants was calculated to be 4.59, 16.9, 325.27, 10.6, 2.55 and 7.62 µM respectively, indicating an almost hundredfold difference between the affinity of the WT and the R679W variant. ***(Fig. 3B, Table S1B)*** Similar ratios were obtained by biolayer interferometry, although with lower measured K_d_ values, 1.5 and 14.5 µM for WT and R679W respectively, a tenfold difference in this case. ***(Fig. 3C, Table S1B)*** Taking the results of all methods into account, it can be stated that the variant R679W nearly eliminates the interaction, while V677M and G734S attenuate it to a notable extent. ***(Table 1)***

**Figure 3.**
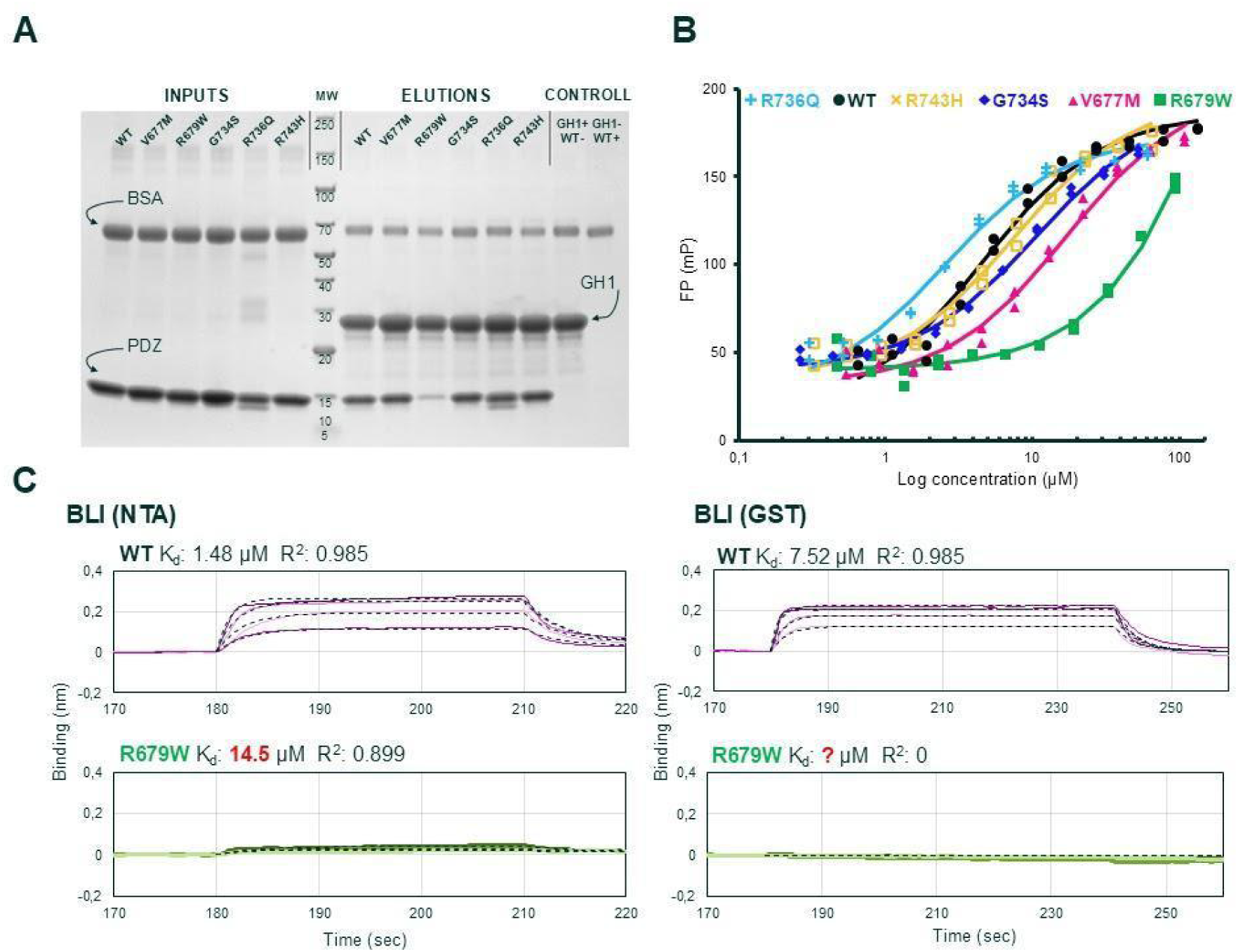
Interaction results with the GKAP partner protein. **(A)** SDS-PAGE of the pull-down assay which allowed for quantitative measurement of relative binding affinities. **(B)** FP results with fitted curves for all variants. **(C)** Examples of BLI datasets (colored lines) with fitted curves (black dashed lines) and resulting K_d_ values and R^2^ values describing fitting quality. Note the similarities in curve kinetics and biolayer response values in the respective experiments.

**Table 1.** Summary of all binding affinity data obtained with Shank1-PDZ variants and GKAP. Relative binding affinities are listed next to each data column in bold italic.

| | ΔAUC (gel) | | $K_d$ (FP) | | $K_d$ (BLI-NTA) | | $K_d$ (BLI-GST) | |
| --- | --- | --- | --- | --- | --- | --- | --- | --- |
| | | (%) | | (μM) | | (μM) | | (μM) |
| **WT** | 21 | **1** | 4.6 | **1** | 1.5 | **1** | 7.5 | **1** |
| **V677M** | 37 | **1.8** | 16.9 | **3.7** | 6.7 | **4.5** | 62.6 | **8.3** |
| **R679W** | 84 | **4** | 325.3 | **70.9** | 14.5 | **9.7** | NA | **NA** |
| **G734S** | 47 | **2.2** | 10.6 | **2.3** | 1.7 | **1.1** | 52.4 | **7.0** |
| **R736Q** | 16 | **0.8** | 2.6 | **0.6** | 2 | **1.3** | 1.8 | **0.2** |
| **R743H** | 25 | **1.2** | 7.6 | **1.7** | 1.4 | **0.9** | 27.3 | **3.6** |

The ligand binding ability of the Shank1-PDZ domain moves in parallel with the destabilizing effect of the mutations: the V677M and R679W variants both exhibit a weaker interaction and a lower T_m_ than the WT domain, whereas the R736Q variant forms a slightly stronger interaction and has a higher T_m_. **(*Fig. 4A*)**

**Figure 4.**
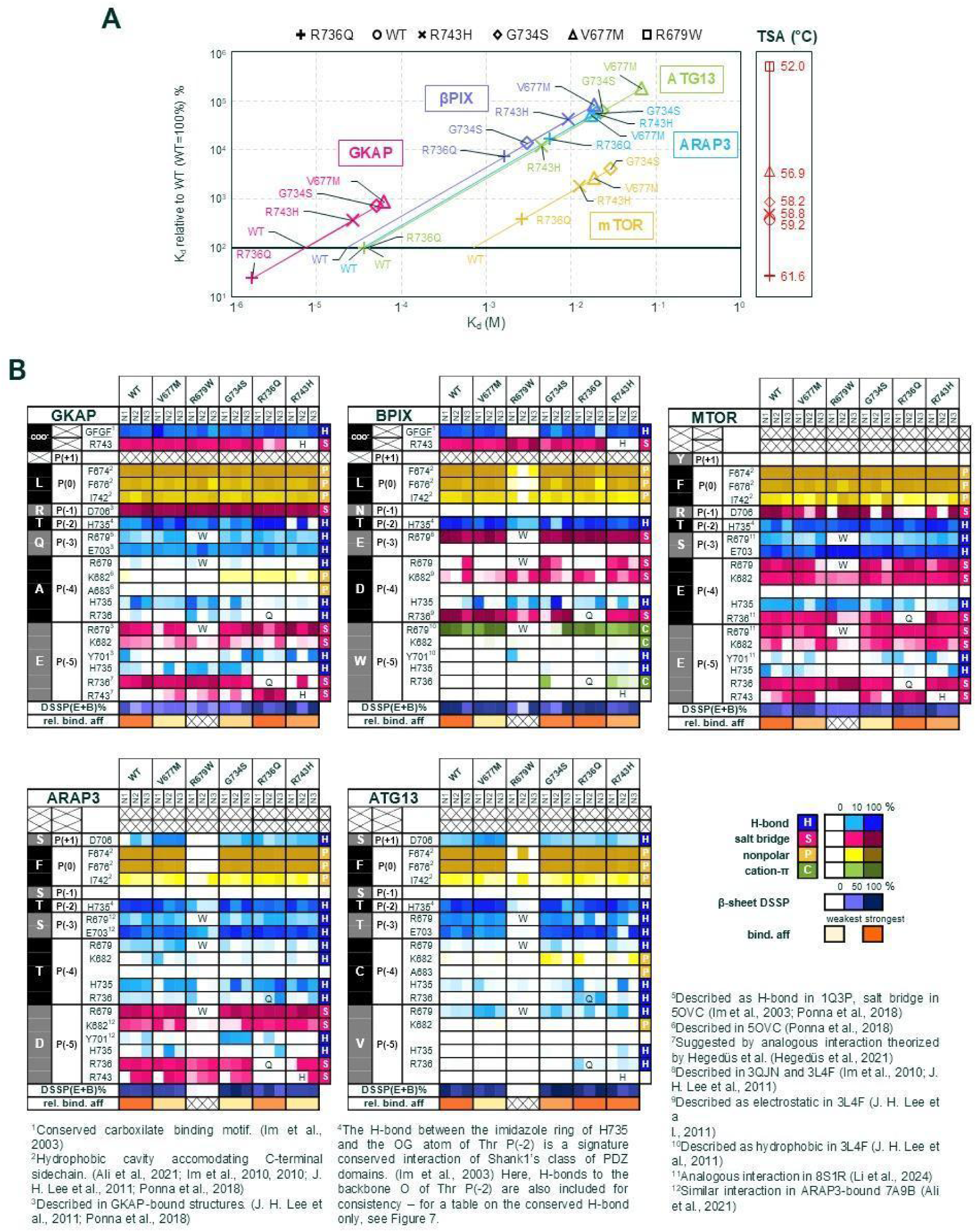
Interactions with GKAP, βPIX, mTOR, ARAP3 and ATG13. **(A)** Relative and absolute K_d_ values of all variants and peptides. For a table of full series of values, refer to the Supplementary Tables. ***(Table S1B)* (B)** Tables representing all PDZ sidechain to peptide sidechain/backbone interactions in our models. The first column shows the peptide sequence, the second the peptide position labels, the third interacting PDZ domain sidechains, and the rest of the columns list the results (abundance of each bond as %) of the three MD runs for each variant. Single letter abbreviations denote the mutated residues in the respective positions. The last two rows show the summarized β-sheet propensity of the peptide, and experimental relative binding affinities calculated from BLI-GST results. Specific interactions were given footnotes listing related literature. For a full collection of exact values refer to the Supplementary Tables. ***(Table S2B)***

**Figure 5.**
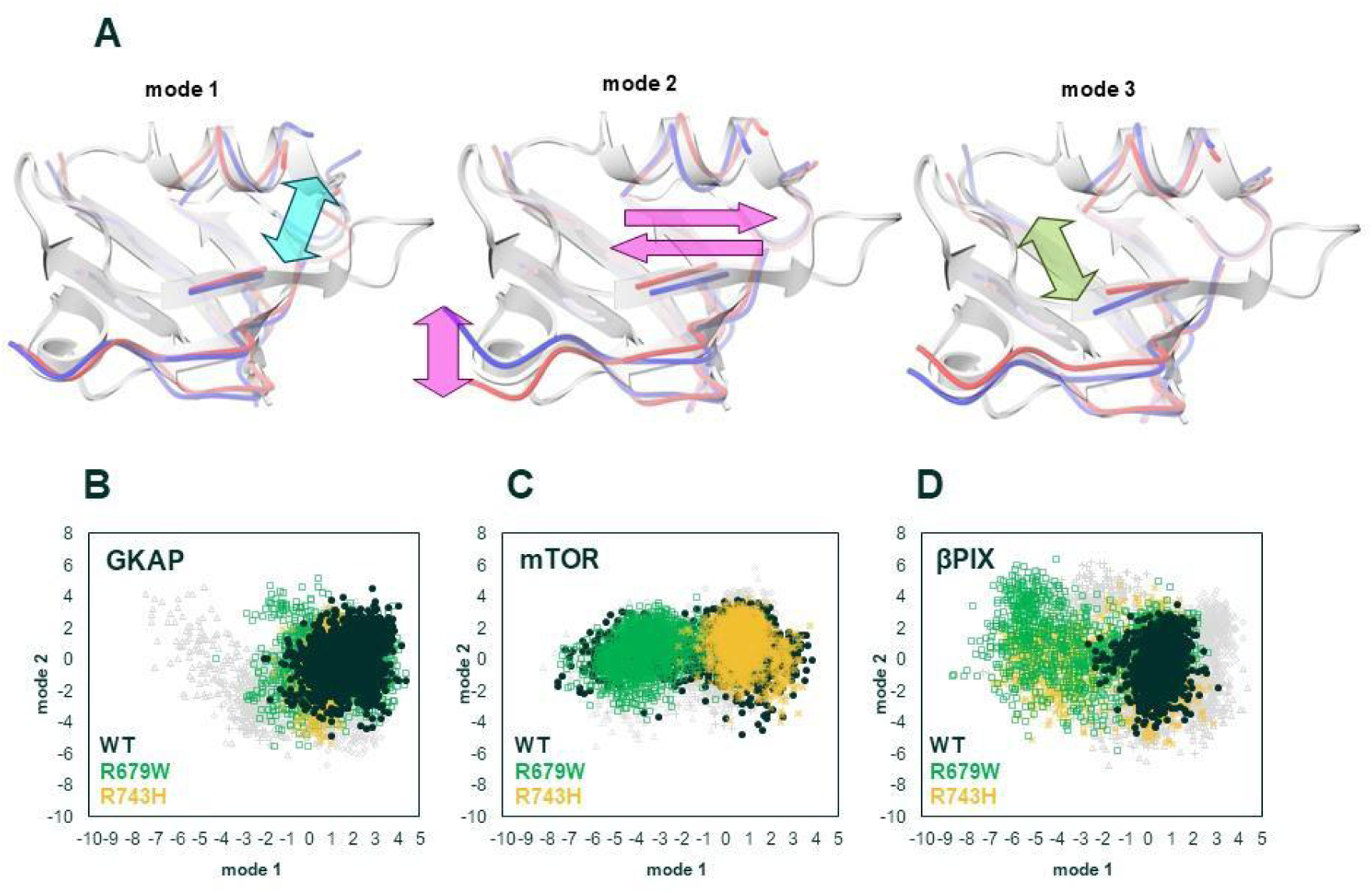
Dynamic properties of the PCA core. **(A)** Models representing the internal motions described by the first three PCA modes. The transparent cartoon representation indicates the PSD95 PDZ3 structure 1BE9 to aid visualizing the location of the core segments within the general PDZ framework. **(B)** Distributions of PCA modes 1 and 2 in the GKAP complexes, with WT, R679W and and R743H highlighted. Note that all three of these variants occupy a more restricted conformational space along mode 1 and the other variants also only scarcely populate the rest of the space. **(C)** Distributions of PCA modes 1 and 2 in the mTOR complexes, with WT, R679W and and R743H highlighted. Note that in this case, two subclusters are clearly identifiable, with relatively equal distribution along mode 1, and the three variants occupying both, the „left” and the „right” clusters respectively, while the distribution along mode 2 is less wide, creating two more symmetric and better defined clusters than seen for other peptides. (D) Distributions of PCA modes 1 and 2 in the GKAP complexes, with WT, R679W and and R743H highlighted. Note that in this case, out of the three highlighted variants, only WT is limited in its motion along mode 1, while the other two variants extend out, although with much lower occupancy than observed for some mTOR complexes. For a full series of figures, see the Supplementary material ***(Fig. S4, Table S2C-D)***.

### Binding affinity changes of the weaker binders exhibit varying correlations with domain stability

In order to investigate the partner binding of Shank1-PDZ variants with other partners, we established a BLI-based protocol using GST-tagged peptides. First, we confirmed that the GST-GKAP-CT12 construct produced similar K_d_ values to those obtained with FP and NTA-BLI as described above. ***(Table 1, Table S1B)*** We observed the very same trends but the absolute K_d_ values appear to be higher than the NTA-BLI results, leading to amplified differences between the variants. As a result, the most unstable variant, R679W did not produce measurable binding with any of the peptides, therefore no K_d_ values were obtained for it. For all peptides, the relative K_d_ values with respect to the WT domain agree well with those reported by Ali et al., with a correlation coefficient of 0.99, but the absolute values are tenfold higher. ***(Table 2)*** We conclude that regardless of these discrepancies stemming from the different methodologies, the relative values within our model system were retained and therefore relevant.

**Table 2.** Summary of K_d_ values with various peptides and WT Shank1-PDZ.

|  | **K<sub>d</sub> (ITC)**<br>**(Zeng et al.,**<br>**2016)**<br>**(μM)** | **K<sub>d</sub> (ITC)**<br>**(Im et al.,**<br>**2010)**<br>**(μM)** | **K<sub>d</sub> (MST. FP)**<br>**(Ali et al.,**<br>**2021)**<br>**(μM)** | **K<sub>d</sub>**<br>**(BLI-GST)**<br>**(μM)** |
| --- | --- | --- | --- | --- |
| **GKAP** | 5.81 |  |  | 7.52 |
| **βPIX** |  | 2.6 | 0.79 | 23.23 |
| **ARAP3** |  |  | *3.65 | 35.76 |
| **ATG13** |  |  | 3.8 | 39.15 |
| **mTOR** |  |  | 84 | *812.5 |

\*value averaged from two different results obtained by same group each

All the peptides turned out to be weaker binders to the wild type PDZ than GKAP, in the following order: βPIX (23.23 uM), ARAP3 (35.76 uM), ATG13 (39.15 uM) and mTOR (720 uM), with the last being significantly, over one order of magnitude weaker than the others. The strongest binder after GKAP is βPIX, the only other C-terminal peptide. Generally, the destabilizing mutations exerted a destabilizing effect on the interactions, with the variants V677M, G734S and R743H consistently producing higher K_d_ values than WT in all experiments. However, the different mutations affected the binding of the peptides in a highly variable manner. For example, R743H, which binds GKAP only slightly weaker than the WT variant, was shown to be the second weakest binder to βPIX after V677M, and produced similar results to V677M and G734S for the ARAP3 interaction. Even more intriguing are the results with R736Q, the variant with the highest thermal stability and GKAP-binding affinity: for most peptides the R736Q variant is a weaker binder than the WT, and even with ATG13 it was merely on par with WT. In summary, the correlation observed between thermal stability and GKAP-binding did not apply to any other of the investigated peptides. Overall, the trends indicate that higher destabilization is accompanied by weaker binding, but there is considerable variability between the ligands. **(*Fig. 4A*)**

### Docking calculations accurately reproduce available experimental structures with wild-type Shank1-PDZ

To rationalize the peptide-specific changes in the interactions with the Shank1-PDZ variants, we performed computational modeling of the PDZ:peptide complexes. Docking simulations were performed and it was determined that those models which have corresponding experimental structures available in the PDB reproduced them adequately, with low RMSD scores between them. ***(Table S2A)*** These models were then used as starting structures for our MD simulations.

### MD calculations suggest variable domain dynamics even for stable complexes

To analyze the PDZ:peptide interactions in more detail, MD simulations were performed on all variant-peptide combinations, using the models created with docking as the starting point. All complexes remained stable during the simulations, with no significant changes in the overall fold of the PDZ domain, with partial peptide dissociation only occurring for some R679W complexes, which is in agreement with the experimental results suggesting it to be the weakest binder. Secondary structure elements as assessed by DSSP remained highly similar to known Shank1-PDZ structures. The peptides exhibited a varying degree of β-strand propensity and low mobility at their conserved P(0)-P(-2) segment and increased dynamics towards the N-terminals. ***(Fig. S2)***

### Specific residue-residue interactions exhibit a high amount of alternative behaviours

Next, to analyze the PDZ:ligand bond network, we extracted the interactions of the PDZ sidechains to peptide sidechains and backbone, and omitted references to backbone-backbone interactions (except in unique cases), as the presence of a backbone-backbone H-bond network in the extended β-sheet binding mode is trivial, and is also redundantly represented by β-strand DSSP annotations. ***(Fig. 4B, Table S2B, Fig. S3)*** The list of specific interactions were chosen based on known interactions identified in experimental structures. Our simulations showed that static complexes suggested by X-ray structures were rarely the case - instead, several interacting peptide sidechains had alternative partners in the PDZ. The C-terminal carboxylate of the peptides of GKAP and βPIX was previously described as forming a conserved H-bond network with the GFGF motif in the PDZ, but an alternative salt bridge with the R743 position has emerged in the simulations. P(-4) and P(-5) of the peptides were particularly prone to alternating between PDZ sidechains. Moreover, some sidechains of the PDZ, such as R679 or H735, were also alternating between peptide sidechain partners. Notable observations regarding these sidechain-sidechain interactions will be detailed in the following paragraphs.

### Peptide specific alterations of internal dynamics are observable throughout the PDZ domain

While this description of binding shows alternative, variable interactions to an extent, in order to describe structural dynamics during interaction, we also performed principal component analysis (PCA) on two subsets of the sequence: a PDZ core, described in more detail by Dudola et al. (Dudola et al., 2020), and β2-β3 loop residues. The rationale behind this dissection is that analysis of the “core”, identified as a set of matching residue positions in 168 different PDZ structures, is expected to eliminate possible largely stochastic differences originating from unique flexible regions during the simulations. On the other hand, the dynamics of the long β2-β3 loop can not be ignored even if its functional significance remains to be established, thus, we analyzed it separately (see below).

PCA of the structural core identifies rearrangements around the ligand binding site, affecting the N-terminal region of the α2 helix (PC1), the C-terminal end of the β3 strand and also the relative position of the α2 helix and the β2 strand (PC2), as well as a kind of narrowing-widening motion of the binding groove (PC3).

One of the notable observations is that several mTOR complexes seem to adopt conformations both along PC modes 1 and 2 that are not or only rarely sampled by other complexes ***(Fig. S4)***. Interestingly, some of the other complexes, like those formed with ARAP3 and ATG13, exhibit opposite motions along the same motional mode. In addition, all mTOR complexes sample a region along the second mode that is only rarely visited by complexes with the other peptides. Although the regions sampled by mTOR-bound domains are not observed entirely consistently for all variants, the emerging pattern is unique enough to raise the possibility that mTOR binding might induce a conformational response different from that in the other complexes. This hypothesis is consistent with the presence of two large Glu sidechains at the N-terminus the mTOR peptide that make contact with the part of the α2 helix exhibiting large displacements along PC1.

### The β2-β3 loop explores different conformations in the complexes

The long, flexible β2-β3 loop is unique to the Shank PDZ domains, and it explored a large conformational space during our simulations, with most of the motion concentrating to its N-terminal half, positions 680-690 ***(Fig. S5A, Table S2E-F)*** NMR chemical shifts indicated the presence of a short helical segment in the free domain (Sánta et al., 2022), and our DSSP analysis of the MD trajectories reveals that this region shows variable helical/turn character, usually below 50%. No discernible pattern was noted in the helical propensity of different complexes and variants. ***(Fig. S5B)***

The loop is relatively far from the mutated sites and remains generally flexible in all complexes. However, principal component analysis reveals some interesting differences between the PDZ variants. Here we focus on those that can be regarded as the most robust ones when taking into account all simulations performed.

The first two principal components are dominated by a shorter region of the loop (approximately 681-690), the first mode corresponding to a motion within the plane of the β-strand formed by the ligand but perpendicular to its long axis. The second mode is largely perpendicular to the first, with the loop alternating between being below and above the plane of the ligand strand. The third PC can be best described as a “compaction-extension” motion of the whole loop. **(*Fig. 6A*)** The three modes cover approximately 31%, 15% and 13% of the variability of the observed loop conformations, respectively. ***(Fig. S5C, Table S2G-H)***

**Figure 6.**
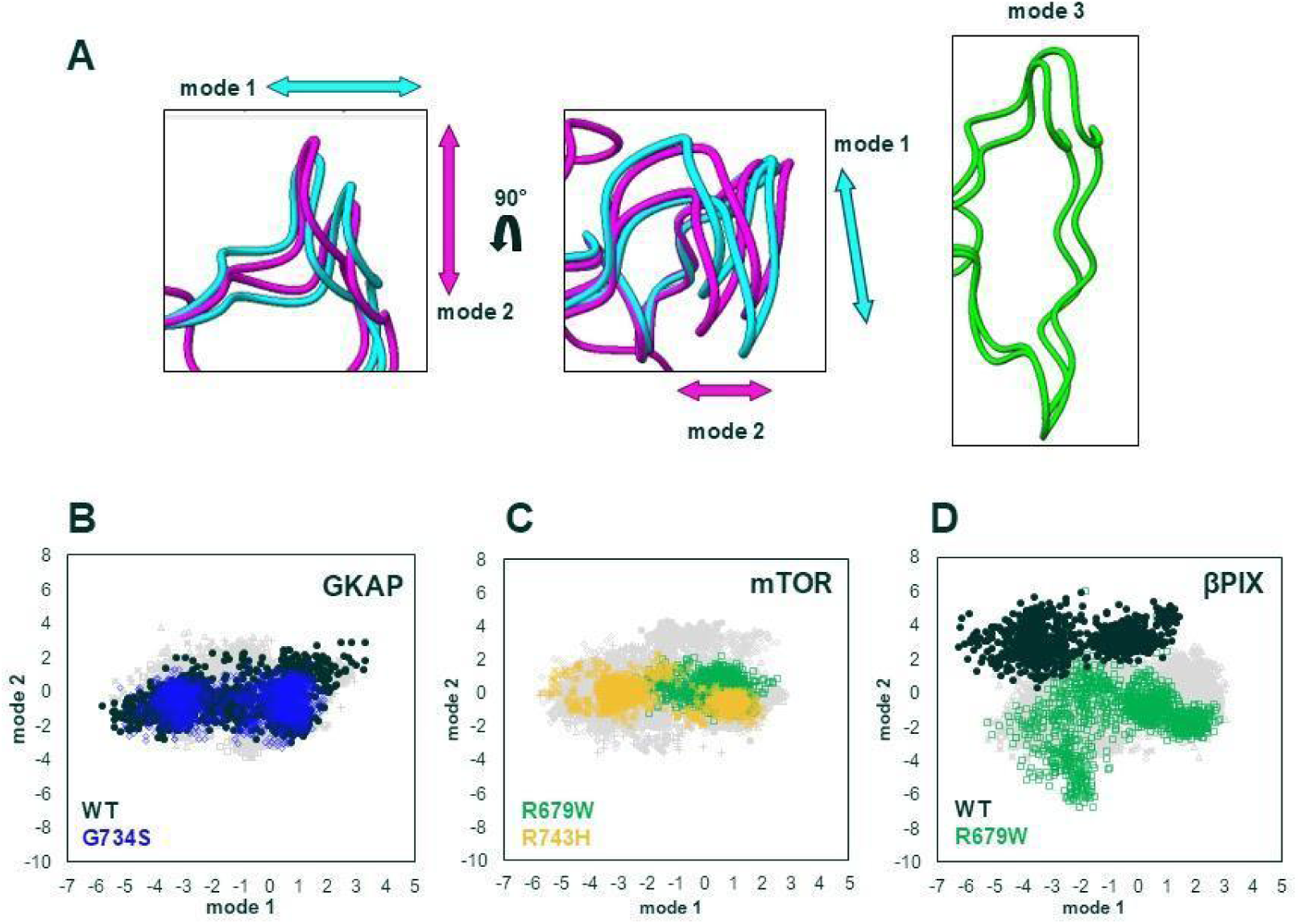
Dynamic properties of the βB2-β3 loop. **(A)** Models representing the loop movements described by the first three PCA modes. **(B)** Distributions of PCA modes 1 and 2 in the GKAP complexes, with WT and G734S highlighted. Note the reduced conformation space explored by the G734S variant, as well as the bimodal distribution of mode 1. **(C)** Distributions of PCA modes 1 and 2 in the mTOR complexes, with R679W and R743H highlighted. Note the differences in the distribution of the two variants along mode 1. **(D)** Distributions of PCA modes 1 and 2 in the βPIX complexes, with WT and R679W highlighted. Note the wider range of motion in mode 2 and the almost complete lack of overlap between the two highlighted variants. For a full series of figures showing PCA mode 1, 2 and 3 distributions, and original PCA data, refer to the Supplementary materials ***(Fig. S5, Table S2G-H)***.

Interestingly, mode 1 exhibits a largely bimodal distribution, with two clusters and a sparsely sampled transition region between them. When considering all our simulations, most peptides and variants populate both clusters, although individual simulations might get stuck in one of them. This bimodality is especially pronounced in the GKAP and mTOR complexes, but present in all to an extent. As for mode 2, the full range of this mode is only explored by the βPIX complexes of different PDZ variants, with the wild type and the R679W exploring the two extremes. For the complexes with this peptide, the conformational space covered by the WT domain is well separated from that characteristic for the variants. ***(Fig. 6B-D, Fig. S54D-G)*** There is also a separation along mode 3: most variants cover a limited region, but R679W and G734S have a tendency to extend further into a second, smaller cluster, with the exception of mTOR complexes only, where no variant explores this area. ***(Fig. S5H-K)*** As it is not expected that our simulations exhaustively cover all relevant conformational states, and we are aware that there might be artefacts that could shift the interpretation of data, we refrain from more specific statements here but note that our simulations suggest that the behavior of the β2-β3 loop is not entirely independent of neither the mutation nor the bound peptide.

### A novel role of R743 in carboxylate binding of the C-terminal motifs in GKAP and βPIX

The main-chain carboxyl group of the C-terminal peptides GKAP and βPIX was expected to form a firm hydrogen bond network with the conserved carboxylate binding 673-GFGF-676 motif of the PDZ domain. However, an alternative conformation emerged in our models, where the terminal carboxyl group flips and forms a salt bridge with R743. This observation suggests a novel, important role of the R743 sidechain besides being part of the “arginine triad” capable of binding P(-5), and hydrophobic stacking. The alternative carboxylate-stabilizing interaction of the R743 sidechain was not considered as an important factor in previous known models of the Shank1-PDZ, which made the abundance of the R743H mutation in tumor samples rather elusive. The carboxylate-binding role offers an explanation to this, as well as the weaker binding affinity, since the observed frequency of interactions of the carboxylate was the lowest in the R743H variant, as H743 does not form this interaction. A notable difference between the two C-terminal peptides analyzed was that while in the GKAP complex, each variant exhibited both C-terminal carboxylate interactions, the βPIX peptide in complex with the R679W variant consistently showed an almost exclusive preference for the R743 salt bridge. **(*Fig. 4B*)**

### The Shank1:GKAP complex: alternate interactions of H735 and P(-5) Glu in the WT and altered dynamics in the variants

Molecular dynamics simulations on the WT domain with the GKAP peptide reproduce most of the previously described residue:residue interactions. PDZ residues F674-F676-L678-I742 form a hydrophobic pocket accommodating the sidechain of P(0) throughout the simulation. Residue D706 forms a characteristic end-on bidentate salt bridge with the P(-1) Arg in ∼80% of the simulation frames. However, the expected P(-3) Gln or P(-5) Glu interactions with R679, especially the former, appear relatively weak, so does the P(-3):E703 interaction, and the P(-5):Y701 bond barely even form in two out of three simulation runs. Remarkably, P(-5) Glu forms alternate interactions with R736, which results in a bent conformation of the peptide that orients the backbones of P(-4) and P(-5) to face H735, which in turn forms transient H-bonds to these positions, while also dynamically changing its side chain orientation to bind to P(-2) Thr. The latter should be its expected conformation characteristic of Class I PDZ domains, however in our simulations its frequency is remarkably low (an average ∼30%) due to the competition with the alternative P(-4)-P(-5) bonds. **(*Fig. 4B*)**

In the missense variants, the hydrophobic P(0) sidechain interaction, as well as the D706:P(-1) Arg salt bridge remains largely unchanged. The Y701:P(-5) interaction appears more often in some of the variants, most prominently in R743H, but still only to a low extent. In the V677M variant, this Y701:P(-5) bond does not form, but some transient P(-5):R743 salt bridges emerge. In the R679W variant, W679 not only deforms the binding site but also exhibits a tendency to form a strong H-bond with the carbonyl O of P(-5) Glu, fixing the peptide in a distorted conformation that deviates from the expected β-strand structure. In the G734S variant, the H735:P(-2) H-bond occurs with lower frequency, while the alternative interaction formed by H735 to P(-5) has a higher tendency to form.

The R736Q variant forms the highest net number of specific interactions with GKAP. **(*Fig. 4B*)** No salt bridge can form between P(-5) Glu and Q736, which results in P(-5) forming end-on salt bridges with another arginine of the “triad”, R743, while simultaneously establishing an H-bond to Q736, which, we predict, together might compensate for the loss of this salt bridge interaction with the mutated R736. **(*Fig. 7*)** This also explains the increased prevalence of H-bonds between H735 and P(-2), as P(-5) can not establish contacts with this histidine in this bent conformation of the peptide. The most prominent changes in the R743H variant are the lack of the aforementioned carboxylate binding interaction of position 743, diminished H735 interactions and an increased prevalence in the P(-5):Y701 H-bond. **(*Fig. 4B*)** For schematic representations, see ***Fig. S3A***.

**Figure 7.**
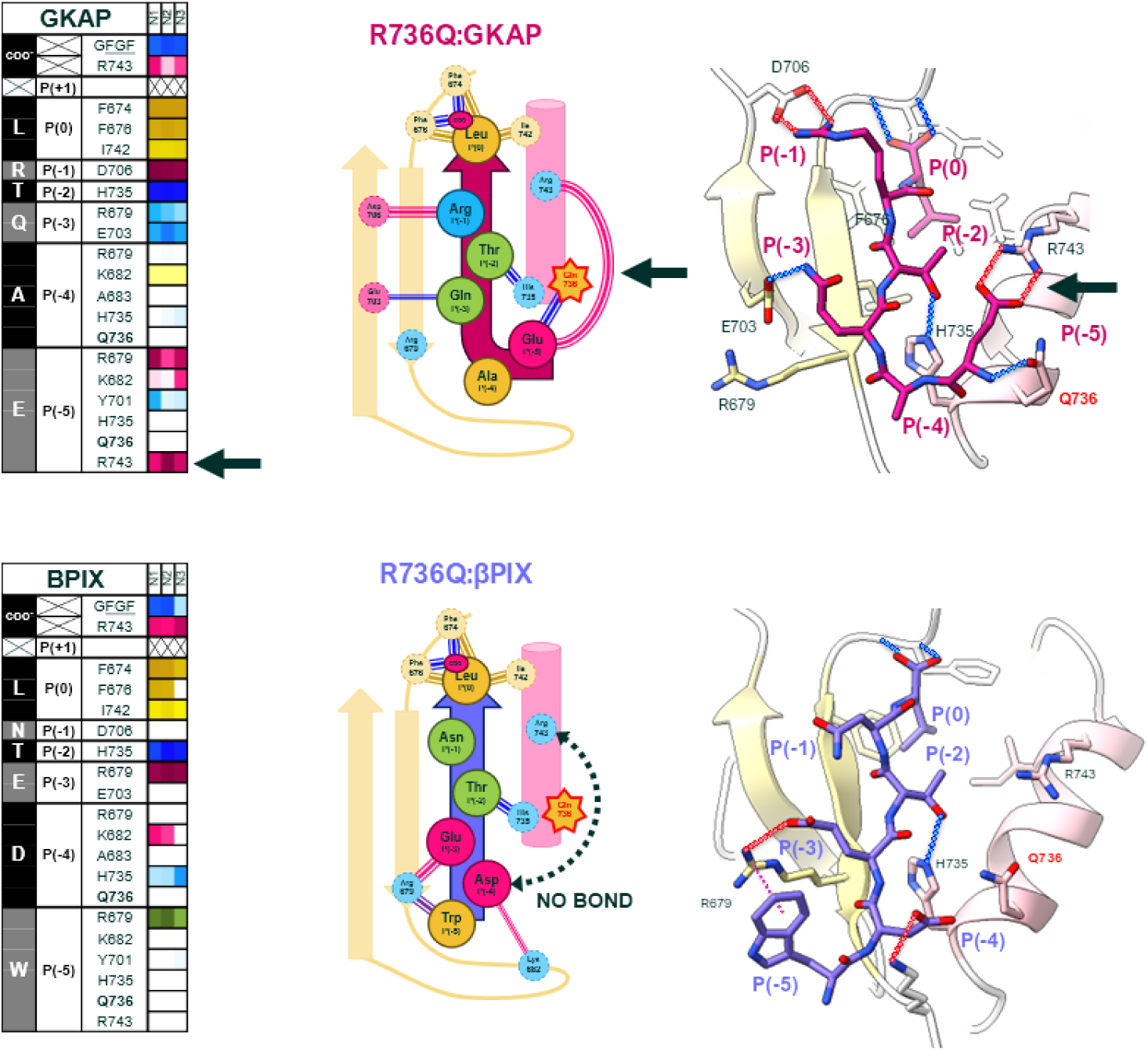
Schematic and 3D models of the R736Q:GKAP and R736Q:βPIX complexes. In the WT:GKAP interaction, P(-5) forms a salt bridge with R736, which in the variant is replaced by a transient salt bridge with R743 which results in a strong hydrogen bond with Q736. The analogous interaction to this in the βPIX peptide is the salt bridge between P(-4) and R736 – in the R736Q variant, this salt bridge is entirely lost and P(-4) does not form alternative bonds. Full a full series of schematic diagrams and figure legends, refer to the Supplementary Material ***(Fig. S3)***.

### Previously unseen cation-π stack in the βPIX interaction varies highly per variant

The peptide of βPIX exhibits a very stable β-strand conformation in complex with Shank1-PDZ WT but a reduced number of sidechain interactions compared to GKAP. The P(0) Leu sidechain remains inserted into the hydrophobic pocket. The H735:P(-2) sidechain H-bond is also more stable, appearing in ∼90% of the snapshots, as are the salt bridges formed by Asp P(-4) and Glu P(-3) with R736 and R679, respectively, in accordance with the experimental structure 3QJN and 3L4F. The only major deviation from these X-ray structures is the behavior of P(-5) Trp, which, instead of being inserted into the β2-β3 loop, stacks onto R679 in a cation–π interaction.

In the V677M variant complex, positions P(0)-P(-2) remain unaffected as expected. The P(-3):R679 salt bridge is slightly weakened, as R679 has a higher tendency to form an intramolecular interaction with E703 (characteristic of the apo WT structure).

The R679W mutation largely deforms the extended conformation of the bound peptide and the complex is characterized by prominent widespread loss of interactions and partial dissociation. As mentioned earlier, the conserved carboxylate binding H-bond network disappears in favour of the alternative salt bridge to R743.

The simulations of the G734S variant suggest a disorganization of the binding network, with several alternative conformations emerging, such as salt bridges and H-bonds of P(-4) to R679, K682 and H735, and alternative cation–π stacking by P(-5) to R736. The repeated runs are highly inconsistent, suggesting destabilization.

In the R736Q variant, the salt bridge between P(-4) Asp and R736 can be considered similar to the one formed by P(-5) Glu in GKAP. However, while Glu P(-5) of GKAP can compensate for the lack of the R679 salt bridge in this variant, by interacting with R743, P(-4) Asp of βPIX, being shorter, can not participate in an analogous salt bridge. **(*Fig. 4B*, *Fig. 7*)**

In the R743H variant, the alternative R736 cation-π stack is also regularly formed during the simulations, resulting in a periodically flipping tryptophane at P(-5).

For schematic representations, see ***Fig. S3B*.**

### A delicately balanced sidechain competition in the mTOR peptide is disrupted in the variant complexes

Although there is no available experimental structure for the mTOR:Shank1-PDZ complex, the PDB entry 8S1R, featuring a similar internal peptide motif (Ac-_[P(-5)]_EESTSFQGP_[P(+3)]_-CONH_2_), can be used as a proxy reference. Residue P(0) Phe of the mTOR peptide remains inserted into the hydrophobic pocket, and P(-1) Arg forms a salt bridge similarly to GKAP, which is generally stable, however a competing intrapeptide cation-π interaction between P(-1):P(+1) emerges which is less prominent in the WT but varies across variants. While the frequency of the H735 H-bond with P(-2) Thr is much lower than for GKAP, P(-3) Ser interacts with E703 more frequently than GKAP’s P(-3) Gln. Residues P(-5) and P(-4), both glutamates, form competitive interactions with R736 and R679, taking multiple turns over the course of the simulation. In the 8S1R X-ray structure, the sidechains are paired as P(-4):R736 and P(-5):R679. Remarkably, H735 prefers forming H-bonds to either of these glutamates rather than to P(-2) Thr. K682 can also form salt bridges with either of the two glutamates, but not with both simultaneously. Overall, P(-5) and P(-4) dynamically interact with R679, K682, H735 and R736 to a nearly equal extent.

Similarly to that observed for βPIX, the effect of the V677M replacement is seen towards P(-3) instead of P(-1). The P(-1) salt bridge forms generally less frequently, and a salt bridge between P(-5):R743 emerges.

In the R679W variant, many interactions are missing. The W679 sidechain is unable to form any salt bridges, and the alternative salt bridges to R736 and K682 fail to compensate for it, as they don’t appear in significantly higher frequency than in the other variants. Interestingly, R743 does not participate in any salt bridges to the peptide, which would be another opportunity to compensate for the lost interactions.

The G734S and R743H variants are characterized by highly variable interaction patterns throughout the simulations, especially in the P(-1) salt bridge. In R743H, interactions with H743 do not occur, and in one case, the increase in the P(-5):Y701 H-bond emerged, similarly to the GKAP simulation.

In the R736Q variant, significant loss of the P(-1) Arg salt bridge was observed, in favour of the intrapeptide cation-π stack, to an extent not observed in any of the other variants. The loss of the ionic nature of the interaction between P(-5) Glu and the mutated R736 can only be partially compensated by R743 because of the competition with R679, with which P(-4) Glu can also interact. H735 remains H-bonded to P(-2) Thr, similarly to the R736Q:GKAP interaction. **(*Fig. 4B*)**

For schematic representations, see ***Fig. S3C*.**

### H735 exhibits unique alternative H-bond in ARAP3

For this complex, comparisons can be made with the experimental structure 7A9B. While most interactions described for this X-ray structure are observed in our simulations, the H-bond between Ser P(+1) and D706 does not seem to be stably formed neither in the WT domain nor most of the variants. In the wild type complex, P(-3) Ser forms a stable H-bond with E703. Besides the Class I characteristic sidechain-sidechain H-bond to P(-2), the side chain of H735 also forms an alternative interaction with P(-4) Thr, creating two main conformational clusters. The P(-5) Asp forms a salt bridge with R679, and, with lower frequency, also with K682, R736 and in some cases, even R743.

The V677M variant exhibits a similar pattern with the notable exceptions of an increase in P(+1):D706 H-bonds. A disruption in the ratio of the two alternative H735 interactions (in WT, the conserved interaction is consistently, strongly preferred) can also be observed in V677M, as well as the rest of the variants. The R679W variant shows a tendency to partially dissociate from the partner peptide. G734S exhibits an increase in the P(-5):Y701 H-bond. R736Q, similarly to the case of βPIX, is characterized by the loss of salt bridges at the mutated position, as well as no R743 interaction compensating for it. **(*Fig. 4B*)**

Overall, for ARAP3, the observed differences between the variants are mostly subtle, with the exception of the dissociating R679W variant.

For schematic representations, see ***Fig. S3D*.**

### ATG13 binding is dominated by H-bonds and nonpolar interactions

ATG13 is the only analyzed peptide with no similar experimental structure available. The interaction pattern of WT:ATG13 appears to be dominated by H-bonds and hydrophobic interactions throughout the length of the peptide. The C-terminal P(+1) Ser forms a H-bond with D706 in less than 10% of the frames, and P(-1) Ser does not participate in any sidechain interactions. The only other prominent sidechain H-bonds in the whole peptide are formed between P(-3) Thr and R679 as well as E703, with a frequency of about 20-50% for both. Contrary to our expectations, the hydrophobic residues in the 680-690 segment of the β2-β3 loop do not contribute to the accommodation of P(-4) Cys and P(-5) Val, instead, the loop remains extended and the hydrophobic sidechain segments face the PDZ backbone.

The interaction pattern varies only little for most of the variant PDZ domains. Similarly to the case of ARAP3, V677M exhibits a slight increase in the P(+1):D706 H-bond, and the variants differ in their loop dynamics, but no other notable differences can be highlighted. The only exception is the complex with R679W which completely dissociates. **(*Fig. 4B*)** For schematic representations, see ***Fig. S3E*.**

## Discussion

### Overview of the variants

All of the most frequent Shank1-PDZ missense variations listed in the COSMIC database affect binding site residues. Considering that besides the binding site, the β2-β3 loop, unique to Shank PDZ domains, is also highly conserved in the family, this distribution of the mutations is not necessarily trivial.

Moreover, all the affected residues are located on the surface of the domain, with their sidechains pointing towards the solvent. Therefore, the large changes observed in thermal stability were rather surprising as changing these residues is not expected to disrupt internal residue networks. Perhaps a simple but partial explanation can be provided for the R679W variant where the R679:E703 salt bridge is disrupted. However, the small destabilizing effect of a Gly to Ser change in G734S and the increased stability in R736Q can not be easily explained on the basis of individual residue properties or clearly identifiable interactions. It should be noted that several X-ray structures of Shank1-PDZ indicate the presence of PDZ dimers (Ali et al., 2021; Im et al., 2003; Lee et al., 2011; Zeng et al., 2016) that might, in theory, increase the number of residue-residue interactions that should be considered when interpreting mutation effects. Our previous NMR study (Sánta et al., 2022) indicated that the Shank1 PDZ is monomeric in solution, and we have not observed any deviations from the expected molecular mass during protein purification. Thus, although the possibility of dimerization, especially upon ligand binding, cannot be completely dismissed for all variants, we chose to interpret the effects on the monomeric structure as the simplest plausible model.

MD simulations indicated only subtle differences between the global structure of the variant PDZ domains in the complexes, with the largest changes observed for the β2-β3 loop. ***(Fig. S2A-B)*** As could be expected, this region showed high flexibility and explored a wide range of conformations, with its helical propensity ranging from 1-48% (G, H, I and T states as assigned by DSSP). ***(Fig. S5B)*** The whole loop, spanning positions 680 to 700, is about one fifth of the length of the canonical domain, while it provides a relatively even larger solvent-accessible surface area, about one fourth of the total of the full PDZ. The sidechain replacements that caused the highest instability were V677M, R679W and G734S - the first two being the closest to the loop in the sequence, while 734 being in close spatial proximity in the structure.

### Dynamics of alternating interactions might be a key contributor to ligand selectivity and binding affinity

Molecular dynamics simulations can capture important aspects of the dynamic behavior of biomolecules, but can not necessarily be expected to accurately reflect all aspects of the actual conformational fluctuations and the delicate balance between all interactions, especially for the diverse systems investigated in this work. Thus, we interpret our analysis as a semiquantitative description that can shed light on several important dynamic aspects and possible interactions, but the actual dynamic balance between the observed conformational states might not be precisely captured.

Generally, the PDZ domains exhibited low fluctuations in backbone RMSD throughout the simulations, however, the peptides, especially their N-terminal segments, turned out to be remarkably flexible. ***(Fig. S2)*** This is also reflected in the behaviour of the corresponding PDZ sidechains of the binding site, detailed below.

The most remarkable general observation in our molecular dynamics calculations was the presence of a number of competing interactions. Most of these are alternative salt bridges, like the one formed by the C-terminal carboxylate and R743, which, in turn, also can pair with Asp/Glu residues at position P(-5). **(*Fig. 8A-B*)** This is somewhat surprising given the expected distance between residues P(0) and P(-5) in the extended conformation of the peptide, but during our simulations, the overall flexibility of the complex allowed the formation of both interactions by R743. Consequently, the Asp/Glu residues at P(-5) and in some cases P(-4) also exhibited alternating motions, either binding R743, or the other nearby arginine residues, R679 and R736, which were previously referred to as the “arginine triad”. **(*Fig. 8C*)**

**Figure 8.**
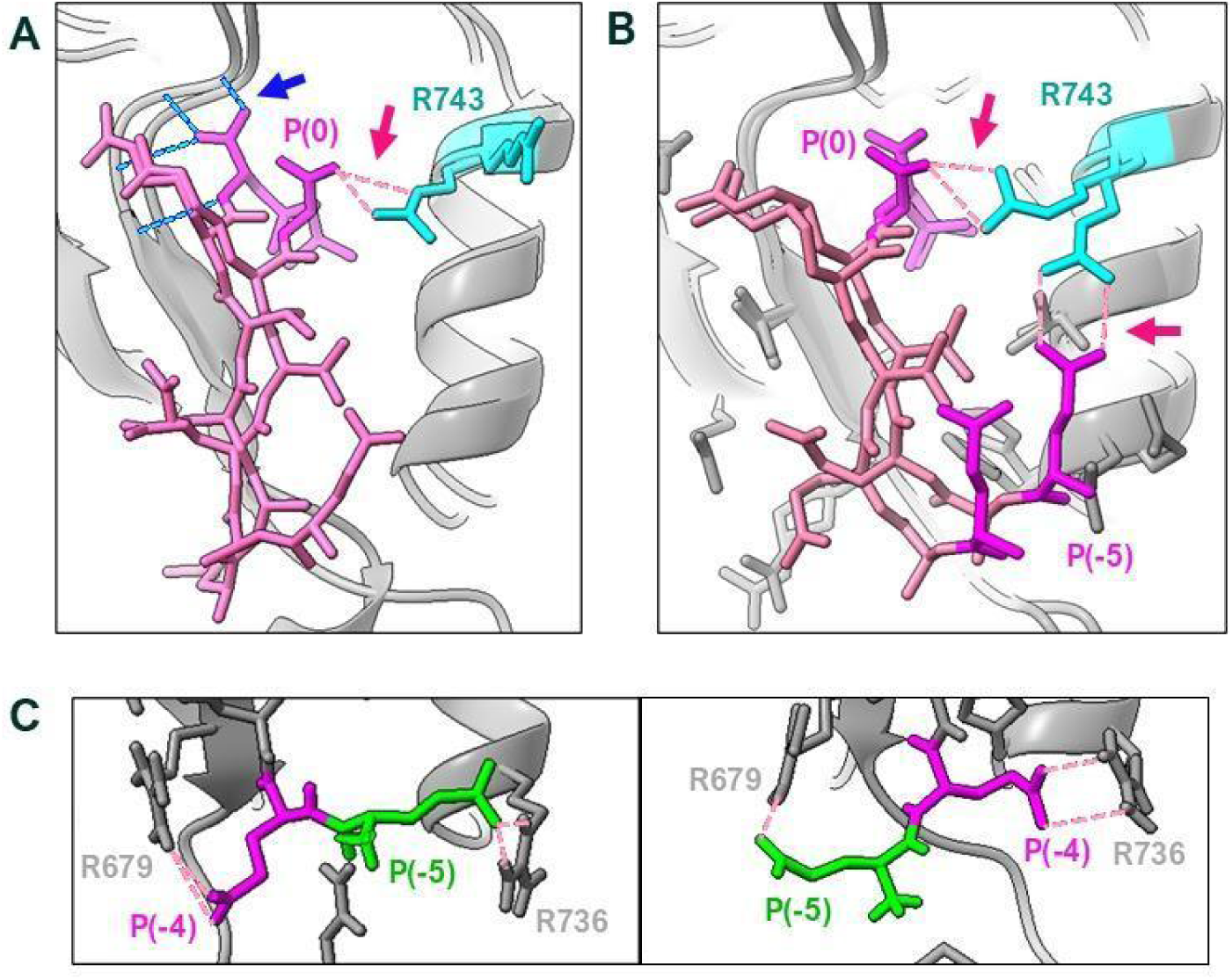
Examples of dynamic, alternating interactions and conformations observed in the MD simulations. Blue and red dashed lines represent H-bonds and salt bridges respectively. **(A)** Alternative conformations of the C-terminal carboxylate in the WT:GKAP complex: in one conformation, it forms a conserved H-bond network with the GFGF motif, in the other, a salt bridge with R743. Note that conformations where both are simultaneously present were also observed, and that the conformation change of the carboxylate did not significantly affect the P(0) Leu sidechain which remains inserted into the hydrophobic groove. **(B)** Alternative conformations of the R743 sidechain in the WT:GKAP complex: in one conformation R743 forms a salt bridge with the C-terminal carboxylate, in the other, with P(-5). **(C)** The „flip-flopping” behaviour of P(-4) and P(-5) of the mTOR peptide, where the two sidechains alternate between salt bridges formed with R679 or R736.

Another important dynamic interaction is formed by the conserved H735 imidazole side chain with the residues in the peptides in positions P(-2), P(-4), and P(-5). **(*Fig. 4B*, *Fig. 9A-B*)** This interaction varies largely for the different peptides and PDZ variants, ranging from a highly dynamic scenario in the case of GKAP and WT PDZ, where intramolecular H-bond formation is also observed, to the stable H-bond with residue P(-2) in the complexes with ATG13 and βPIX. ***(Fig. 9C, Fig. S6, Table S2I-K)*** The difference in the behavior of H735 with different partner peptides suggests a possible role of H735 in partner selectivity. This is rather unexpected as H735:P(-2) was believed to be a static interaction with the only role of stabilizing the peptide. It should be noted that our molecular dynamics simulations can not account for the changes of the protonation state of the imidazole ring, which was set once at the beginning of the simulations, was constant throughout, and was identical for each repeated run. Variations of protonation state might also contribute to the variability of the interactions.

**Figure 9.**
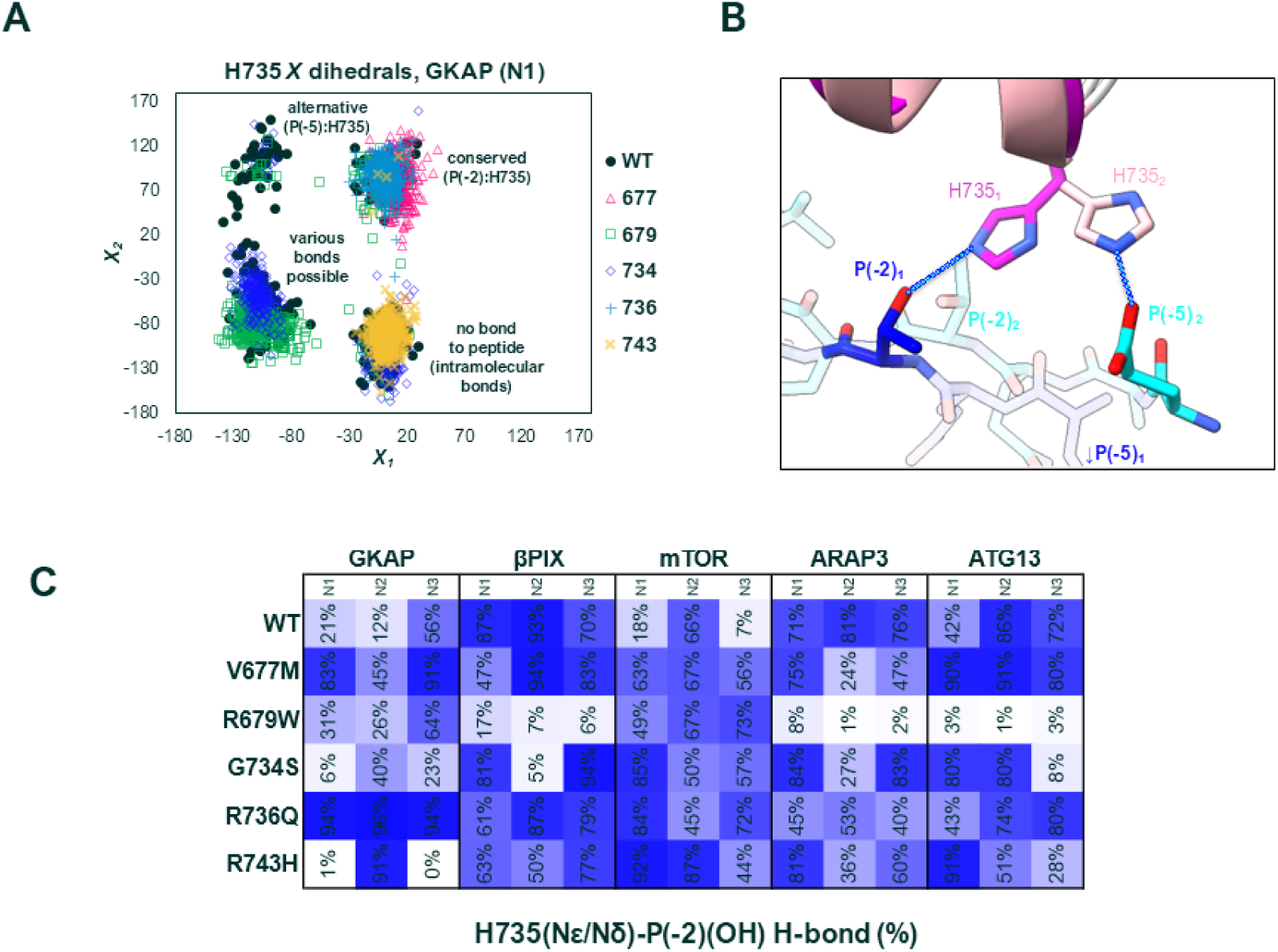
Dynamics of the H735 sidechain. **(A)** *Χ*_1_-*Χ*_2_ dihedrals explored by the H735 sidechain during WT:GKAP binding. Note that the *Χ*_1_-axis was shifted by 180° for better visualization. Each cluster is labelled with the sidechain bond that is probable to occur in said conformation. **(B)** Structures representing two possible extreme conformations of H735 and its H-bonds in the WT:GKAP complex. Darker colors denote the conserved conformation, which is labelled „1”. Possible H-bonds in each conformation are represented with blue, dashed lines. **(C)** Summary table showing the presence of the characteristic H735 H-bond throughout MD simulations as percentage, with proportional color intensity. Note that this data is nearly identical to what is seen in Fig. 4B, however the latter also includes bonds where the backbone of the peptide is involved, while this table only shows sidechain-sidechain bonds.

### Functional equivalence of sidechains in different peptides

We have also observed interactions from the peptide side that can more or less replace each other, suggesting the presence of some degree of functional equivalence between sequential positions in the partner peptides. For example, amino acid residues in peptide positions P(-1) and P(+1), typically positively charged ones, are often interacting with the PDZ residue D706. On the other hand, E703 tends to form an H-bond with P(-3), most commonly a serine in both internal and C-terminal ligands. P(+1), P(-1) and P(-3) tend to always remain pointing towards and interacting with the β-strand side of the binding pocket, a common orientation seen in similar PDZ domains. (Lee & Zheng, 2010)

Positions P(-4) and P(-5) show considerable variability, but with a preference towards glutamate or aspartate. Side chains in both positions can form salt bridges with the PDZ residues R679 and R736. Hydrophobic sidechains are also common in these two positions, and some models suggest that if present, the β2-β3 loop can stack over them to accommodate them (Im et al. 2010; Ponna et al. 2019), although in our models, this type of interaction was not prevalent. Our simulations suggest that the interactions formed by P(-4) and P(-5) can drastically influence the backbone conformation of the peptide, with a R736:P(-5) or R743:P(-5) interaction consistent with the emergence of a kinked structure, while hydrophobic interactions with the P(-4)-P(-5) positions result in altered β2-β3 loop dynamics in the PDZ.

For some of the interactions, the sum of intermolecular sidechain bonds correlates well with binding affinity, highlighting the example of the R736Q variant: compensatory interactions are formed which restore the stability of the complex in the GKAP interaction, while this phenomenon is not observed during, for example, βPIX binding, resulting in the loss of a significant salt bridge. **(*Fig. 7*)** However, not all of the simulations can be interpreted in such a straightforward manner. While in general, due to the extended β-sheet binding mode, the β-strand propensity of the peptide should be a good approximator of complex stability, those exhibiting a kinked peptide conformation were an exception from this rule and therefore a stable complex could exist even if the extended β-sheet structure does not fully form. Moreover, considering the relatively weak nature of the interactions (micromolar range, even using the measurement techniques that produced the lowest K_d_ values), it is a possibility that energetic contribution of overall PDZ stability versus the peptide binding network are on par, creating the complex response to the sidechain replacements observed in the experimentally determined binding affinities.

### Interplay between residue-residue interactions and internal rearrangements

Understanding the relationship between binding affinities and mutation induced structural perturbations is further complicated by the apparent dynamic nature of the peptide binding. Several transient, competing interactions were suggested by the MD simulations, including that of the C-terminal carboxylate and H735, previously considered to be stable interactions due to their conserved nature. This, in addition to the dynamics of P(-5) in relation to R679-R736-R743 that was also proposed by previous studies (Hegedüs et al., 2021) and the remarkable mobility of the β2-β3 loop paints a picture of a highly flexible complex.

The observed residue-residue interaction pattern offers some general clues on how these interactions contribute to the overall strength of the binding. The C-terminal ligands GKAP and βPIX show the strongest binding, indicating that a key interaction is formed in this region. The largest difference in the interaction pattern between these two peptides is observed for position P(-3), with GKAP forming hydrogen bonds and βPIX dominated by charged interactions. Besides, GKAP forms strong charged interactions with its P(-1) residue Arg.

The internal ligands ATG13 and ARAP3 show weaker binding that can be rationalized by the lack of the C-terminal interaction. This is consistent with the observations of Ali et al. regarding altered ARAP3 peptides and their binding affinities. (Ali et al., 2021) Both show a strong tendency to form hydrogen bonds with their P(-2) and P(-3) residues. These considerations emphasize the role of both the type and position of the interactions with a potential major contribution to the binding strength.

These very simple observations, however, do not explain why the internal ligand mTOR is the weakest binder, despite also forming hydrogen bonds with its P(-1) and P(-2) residues in addition to establishing a number of charged interactions at P(-4) and (P-5).

Principal component analysis of the molecular dynamics trajectories focusing on a set of core residues defined in a previous study suggests that the dynamic properties of mTOR complexes might be different from most other systems studied here. Although the differences observed are subtle, and not observed for all mTOR complexes with high occupancy, both the first and second PCs suggest that the conformational rearrangements occurring upon ligand binding might be different for mTOR than for the majority of the complexes, especially those formed with GKAP. The first PC shows a bimodal distribution with a state not substantially occupied by the complexes with any other peptide studied here. The second PC mode shows a region populated by all mTOR complexes but only by a few others. The conformational rearrangements described by these modes relate to the relative orientation and distance of the β2 strand the α2 helix, forming the binding site for residues P(-3)-P(-5). These observations suggest that not only the residue-residue interactions but also the conformational changes accompanying their formation might play a role in determining the strength of the PDZ:peptide complex.

*Possible consequences of the Shank PDZ mutations in associated diseases* All of the partner peptides analyzed in this study have potential or proven connections to disease pathways. Most notably, Shank proteins all have an essential role in regulating postnatal brain development via the Akt/mTOR pathway, however, little is known about the contribution of its individual domains. (Burbach, 2016) The GKAP:Shank interaction via the PDZ domain is crucial as it tethers various other PSD proteins to the NMDA receptors, consequently, GKAP is often described as the main interaction partner of the Shank PDZ domains. However, the contribution of this complex to the aforementioned growth factor pathway is elusive. (Naisbitt et al., 1999) Involvement of Shank2 in oncogenesis can be explained, at least in part, by the requirement of the interaction between the Shank2-PDZ and βPIX to regulate Hippo signaling. (Xu et al., 2021) For the remaining two peptides investigated in this study, we found no direct evidence in the literature on how they influence Shank1 function, however, ATG13 is also a member of the TOR pathway and is involved in autophagy, while ARAP3, similarly to βPIX, is involved in GTPase mediated signaling and regulates cytoskeletal remodeling. (Jung et al., 2010; Myers & Casanova, 2008)

Loss of Shank function can be one of the mechanisms behind ASD, as observed in Phelan-McDermid syndrome and experiments with KO and KI mice. (Hagmeyer et al., 2018; Qin et al., 2022; Sungur et al., 2014) Conversely, overexpression of Shank proteins is oncogenic. (Chen et al., 2022; Wang et al., 2020; Xu et al., 2021) Interestingly, all of the cancer associated variants analyzed in our study turned out to have destabilizing, loss-of-function effects, while the one found in also ASD was the variant with increased stability. However, as the above example with βPIX and the Shank2-PDZ shows, loss or gain of function in a single domain does not necessarily translate to models of under- and overexpression respectively. Instead, in an interaction network as abundant and complex as the one Shank1 is centered at, the perturbing effects of a mutation might require an explanation within a larger context. Our results show that each disease-associated variant has its own unique pattern in affecting these five interactions, to which overall domain stability was only a partial contributor. This suggests a highly complex perturbing effect on the interaction network of the domain, with binding preferences shifting from one partner to another in some cases. Systems biology simulations had recently demonstrated that small changes in the initial conditions, such as a single K_d_ value, can result in a large nontrivial perturbation on the PSD interaction network. (Miski et al., 2023) Experimental data like ours can be used as input to refine such modeling approaches to address similar systems more realistically.

Our results highlight the intricate interplay of residue-residue interactions in partner binding and show that the effects of mutations can be highly partner-specific in the case of a promiscuous partner binding domain. Notably, our Kd values span 4 orders of magnitudes (2 for the WT domain and 3 for each variant), covering almost the full range of SLiM:domain interactions observed in a recent large-scale study (Subbanna et al. 2025). We believe that the complex analysis of multiple affected interactions like our study can contribute to a more comprehensive description of the rewiring of protein:protein interaction networks upon specific mutations, leading to a deeper understanding of the interplay between molecular-level changes and phenotypes.

## Materials and Methods

### Selection of variants and interaction partners for the study

The only ASD associated missense mutation on the PDZ domain of Shank1, causing the R736Q variant, had been described by Sato et al. (Leblond et al., 2014; Sato et al., 2012) Other cancer associated somatic variants were collected from the COSMIC database. (Sondka et al., 2024) In this case, the four most frequently mutated positions (at the time of data collection) of the region were chosen for the study. Where multiple mutations were documented at the same position, the most frequent was picked. The C-terminal peptide of GKAP was chosen as the first model system as it is the most studied, and the additional peptides were chosen with the following considerations: diversity in binding motifs; diversity in associated GO terms; preference for ligands with existing experimental structures in complex with the Shank1-PDZ; preference for ligands with K_d_ values known from literature. Peptides for experiments were designed based on peptide constructs with known K_d_ values found in papers associated with PDB structures. (Ali et al., 2021; Lee et al., 2011; Zeng et al., 2016)

### Expression of constructs

The wild type Shank1-PDZ domain and its missense variants as well as the 186 amino acid long C-terminal segment of GKAP (GKAP-GH1) were expressed in *E. coli* strains. The Shank1-PDZ construct spans from G654 to K768 on the human Shank1 reference sequence (NP_057232.2, Q9Y566-1). A rat Shank-1 ORF plasmid (kindly provided by Enora Moutin from the University of Geneva) was used to amplify the insert for cloning. Note that in this region, Rat and Human ORFs code for the same protein sequences. The insert was ligated into NdeI and BamHI sites of a modified pET-15b vector (Novagen) that contains a tobacco etch virus (TEV) protease cleavage site instead of the thrombin sequence. The expressed construct is equipped with an N-terminal 6xHis tag. Missense mutations were introduced into the Shank1-PDZ plasmid by site directed mutagenesis using PCR primer pairs bearing the desired mutation for the amplification of the whole plasmid. The construct coding the C-terminal segment of GKAP (GKAP-GH1) corresponds to the rat GKAP1a D481-L666 region (Uniprot P97836-5) and its insert was picked up from a rat GKAP1a ORF template kindly provided by Enora Moutin. This 186 amino acid long segment differs from the human GKAP sequence of 792-977 only at K501 (Uniprot P97836-5), the human protein (Q9D415-1) has a glutamine amino acid at this position. This insert was ligated into the same modified pET-15b plasmid as used for Shank1-PDZ. The GKAP construct labelled CT43 is 100% identical to the human sequence covering residues 935-977 and was created using the same methods. Constructs coding 12 amino acid long peptides incorporating the binding motifs of GKAP, βPIX, ARAP3, mTOR and ATG13 were designed based on the human protein sequences (Uniprot entries Q9D415-1, Q14155-1, Q8WWN8-1, P42345 and O75143-1 respectively), using peptides listed by Ali et al. as a template. (Ali et al., 2021). In vitro mutagenesis was used to introduce the sequences into a pGex-4T1 vector, resulting in a GST-tagged construct containing a thrombin site. ***(Table S1A)***

### Protein purification

Protein expression from all constructs were induced in BL21 (DE3) cells (Novagen) with 1 mM IPTG at 4 MFU cell density and the recombinant proteins were expressed at 20 °C for 16 h in LB medium. Cell pellets were lysed by ultrasonic homogenization in 10% cell suspension using a lysis buffer (50 mM NaPi, 300 mM NaCl, pH 7.4). After homogenization, cell supernatants were purified by IMAC using 5 ml Nuvia™ Ni-affinity column (Bio-Rad), followed by His-tag removal from Shank1-PDZ proteins with TEV protease. Proteins were further purified by ion exchange chromatography, using 5 ml High Q column for Shank1-PDZ constructs (recombinant Shank protein was collected in the flowthrough fraction) and High S columns for the GKAP-GH1 construct (recombinant GH1 was eluted by NaPi buffer with 1 M NaCl). Protein samples were concentrated by ultrafiltration using Amicon® Ultra Centrifugal Filter with 3 kDa molecular weight cut off value, and the buffer of recombinant proteins was changed to low salt NaPi Buffer (50 mM NaPi; 20 mM NaCl, 0.02% NaN_3_; pH 7.4). The concentration of the proteins was measured by their absorbance at 280 nm using a NanoDrop2000 photometer, while their purity and exact molecular weight was analyzed by SDS-PAGE and LC–MS. Before pull-down and BLI assays, Shank1-PDZ constructs were purified additionally by reverse IMAC (the protease-treated sample is loaded onto an IMAC column - the cleaved protein remains in the flowthrough fraction) in order to get rid of trace amounts of His-tagged protein and peptide impurities. In the course of this step 20 mM of imidazole and 0.02% Tween 20 as well as 30 µl of equilibrated Nuvia™ IMAC resin was added to every mg of IEC purified PDZ samples followed by a 2 min RT incubation with gentle but thorough mixing. The beads were sedimented by centrifugation, and the flowthrough was retained. The production of the GST-tagged 12 amino acid peptide constructs was identical up to the purification steps. Cell lysates were then loaded onto a Bio-Rad Poly-Prep® Chromatography Column filled with Cytiva Glutathione Sepharose™ 4B beads. The aforementioned low salt NaPi Buffer was used for the binding step, and the same buffer with additional 20 mM glutathione was used to elute the proteins. No additional steps were used for these constructs because sample purity was sufficient and the glutathione content in the buffer was negligible at the dilutions used for later experiments.

### Pull-down experiments

For pull-down assays, the 6xHis-tagged GKAP-GH1 construct was used as the bait and the Shank1-PDZ wild type or its variants lacking the His-tag was the prey molecule. The proteins were purified for the assay as stated above. All steps of the protocol were performed in phosphate buffer saline (PBS) (50 mM NaPi; 20 mM NaCl; 0.1% Tween 20; 0.1% BSA; 20 mM imidazole; pH 7.4). For each reaction, a pellet of 20 µl 50% suspension of Nuvia IMAC beads equilibrated in PBS was mixed with 70 µM GH1 protein diluted in 40 µl of PBS buffer followed by a 2 min. RT incubation with mild shaking. In the GH1- WT+ control sample, 40 µl of PBS was added without GH1 protein. After the incubation, beads were washed 3 times with 150 µl of PBS and 75 µM Shank1-PDZ diluted in 30 µl PBS was mixed with the pellet of beads. In the case of GH1+ Shank1-PDZ – control sample PBS was added without Shank1-PDZ protein, while wild type Shank1-PDZ was added to the GH1- WT+ control sample. Addition of the prey was followed by a 2 min. RT incubation and three wash steps, then 20 µl of elution buffer (0,5 M imidazole in PBS) was added to the pellet of beads. After a 2 min. RT incubation step with mild shaking, 15 µl of 3x Laemmli buffer was added to the samples followed by an additional 2 min. shaking incubation. SDS-PAGE images ***(Fig. S1)*** were analyzed with ImageJ. (Schneider et al., 2012)

### Biolayer Interferometry

Prior to experiments, all Ni-NTA biosensors (Fortebio, USA) were hydrated in kinetic buffer (50 mM NaPi pH 7.4, 20 mM NaCl, 20 mM imidazole, 0.1% BSA, 0.02% Tween 20, 0.02% N_3_Na) at 25°C for 10 min. The ligand, the 6xHis-tagged GH1 or CT43 protein was immobilized on the surface of the biosensor at a 5 µg/ml (0.23 and 1 µM respectively) concentration in kinetic buffer for 120 sec. The initial baselines were then recorded in kinetic buffer for 30 s using a BLItz system (ForteBio, USA). The association and dissociation sensorgrams of WT and variant PDZ domains at concentrations ranging from 0.5 μM to 4 μM were recorded for 30 sec each in kinetic buffer, too. The equilibrium dissociation constant (K_d_) was determined from the BLI data at various concentrations of the PDZ domains using the global fitting method provided in the BLItz data analysis software. The BLI experiments with the anti-GST biosensors were conducted nearly identically, with the following exceptions. Imidazole was omitted from the kinetic buffer. The GST-tagged proteins were diluted to 20 µg/ml (0.7 µM), and the PDZ domains were measured at concentrations ranging from 1.25 μM up to 60 μM. Association and dissociation steps were extended to 60 s.

### Fluorescence polarization spectroscopy

Interaction of FITC-labeled CT12 GKAP peptide (FITC-IEIYIPEAQTRL, BioBasic) with the wild type and variant Shank1-PDZ samples were measured in kinetic buffer (50 mM NaPi pH 7.4, 20 mM NaCl, 20 mM imidazole, 0.1% BSA, 0.02% Tween 20, 0.02% N_3_Na) at 25°C. Series of 1.7 fold dilutions of the PDZ domains were prepared (70 µl of sample into 100 µl of kinetic buffer) with 50-150 µM starting concentrations. The concentration of the FITC-C12 peptide was 10 nM constant in all samples except the reference blank sample, where no peptide was added. Fluorescence polarization of the samples was measured in a black, flat 96 well plate (Greiner bio-one) in duplicates (100 µl/rxn) using a Spark 20M multimode microplate reader (TECAN) in filter mode. No Shank was added in the Reference samples, while in the Reference blank there was neither Shank, nor peptide. G-factor was estimated to be 1.067. The equilibrium dissociation constants (K_d_) were calculated from the inflection point of the fitted titration curves using ProFit. (available at https://github.com/GoglG/ProFit)

### Thermal shift assay

Shank1-PDZ protein samples were analyzed in 0.2 µg/µl (15.2 nM) final concentration in low salt NaPi buffer supplemented with Sypro Orange dye in 5x final concentration. Duplicates of 25 µl samples were analyzed in a PikoReal real-time PCR instrument. The temperature was elevated from 30°C to 80 °C with a 0.2 °C / 30 sec increment followed by 30 sec incubation and data collection protocol. Fluorescent data was collected in the SybrGreen channel. The negative derivative of the RFU was plotted as a function of temperature in order to get melting temperature values as the peak of the melting curves.

### ECD spectroscopy

Wild type and variant Shank1-PDZ proteins were measured by ECD spectroscopy using a JASCO J-1500 spectrometer (JASCO Corporation, Tokyo, Japan). All samples were diluted to 7 µM in low salt NaPi buffer. ECD spectra were recorded at 30 °C using 0.1 cm path length J/21 quartz cuvette (Jasco Corporation) and 300 µl sample with the following settings: 195–260 nm spectral range, 50 nm/min scanning speed, 1 nm bandwidth, 0.2 nm step size, 0.5 s response time and 3 scans of accumulation and baseline correction. For melting experiments, temperature scan measurements were carried out with the same settings above. Full ECD spectra were recorded at increasingly higher temperatures starting with 30 °C up to 70 °C in 5 °C increments. Thermal denaturation of the protein samples was demonstrated by the decrease of CD values at 200 nm and the increase at 210 nm, since these changes reflect the increase in the proportion of random coils. Secondary structure compositions were estimated with BeStSel. (Micsonai et al., 2025)

### Peptide docking

The WT and variant PDZ domains in complex with a C-terminal peptide “EAQTRL” of GKAP and apo PDZ domains were modelled in UCSF Chimera 1.18 (Pettersen et al., 2004) using MODELLER (Webb & Sali, 2016) and refined with FoldX (Schymkowitz et al., 2005), using structure 1Q3P from the PDB as a starting point. These structures were then used as templates for docking simulations. Peptides were docked using AutoDock Crankpep (ADCP). (Zhang & Sanner, 2019) First, a structural model for the WT domain in its apo form was created as described above, then the peptides “EAQTRL”, “WDETNL” and “DTSTSFS”, corresponding to GKAP, βPIX and ARAP3, respectively, were docked into this model, and then compared to the experimental structures available under the PDB IDs 1Q3P, 3QJM and 7A9B, respectively. For more details on PDB structures and constructs, see ***Table S1A, S2A***. RMSD scores between corresponding structures were calculated in UCSF ChimeraX 1.8 (Meng et al., 2023), with sidechains compared only up to the Cβ atoms. This is due to the ADCP output producing incomplete ligand sidechains, where each sidechain is represented by only a Cβ-Cγ pair with varying length of the covalent bond to represent volume. These RMSD scores remained below <1 Å for all three complexes, therefore we concluded that the method reproduces the experimentally determined bound conformations sufficiently well. Then, the remaining two peptides were docked into the same WT model (“EESTRFY” of mTOR and “VCTTSFS” of ATG13) and all docking simulations were repeated for each of the PDZ variants.

### Molecular dynamics simulations and analysis of structural ensembles

Explicit solvent molecular dynamics simulations of WT and all variants paired with each ligand peptide were performed with GROMACS (Páll et al., 2020) on the Komondor supercomputer. The AMBER99SB-IDLN force field was used, with a box setting of 1.0 n minimum distance. An 1000 ns time interval was simulated (for each complex), from which, 500 snapshots were taken. Each simulation was repeated an additional two times, using identical initial parameters, meaning altogether 3 simulations, for each of the peptide-variant pairs. Initial structures and conditions for the 3 repeats were identical. Protonation state was adjusted automatically by the pdb2gmx tool which resulted in a neutral H735 for all complexes. With 6 variants and 5 peptides (30 complexes), repeated 3 times, this resulted in 90 simulations from which 90 protein ensembles were created, each containing 500 snapshots, which we provide in the Supplementary Repository in PDB format. The snapshot ensembles were visually inspected in Chimera 1.8. (Pettersen et al., 2004) Structure superposition of the ensembles and RMSD calculations were done with LSQMAN and MOLMOL. (Kleywegt, 1996; Koradi et al., 1996) Secondary structure composition was calculated with DSSPCont. (Carter et al., 2003) HBPlus (McDonald & Thornton, 1994) was used to extract lists of possible H-bonds, defined as 4 Å. To calculate specific atom-atom distances, salt bridges (according to the distance-based criterion described by Barlow and Thornton (Barlow & Thornton, 1983), which includes not only charge-reinforced hydrogen bonds but also other specific charged interactions), cation-π interactions and *Χ*_1_*, Χ*_2_ torsion angle distributions, in-house Perl and Python scripts by Z. Gáspári and Z. Kálmán were used. Interactions between hydrophobic sidechains and segments were identified based on a distance of 4.0 Å between their CB atoms, or in case of arginine sidechains, averaged from distance to its CB, CG atoms. For the C-terminal hydrophobic pocket, this constraint was raised to 6.0 Å, due to the larger sidechains involved, and based on the distances observed on X-ray structures. The C-terminal conserved hydrogen bond with the GFGF motif was described by a single number obtained by averaging all such interactions of the F674, G675 and F676 sidechains, including those involving the PDZ backbone. Additionally, FoldX (Schymkowitz et al., 2005), PDBePISA (Krissinel & Henrick, 2007) and CaPTURE (Gallivan & Dougherty, 1999) were used on some individual snapshots for further verification of findings. RMSF for the loop region was calculated with the MDAnalysis library. (Gowers et al., 2016) Principal component analysis was conducted using the ProDy package. (Zhang et al., 2021) For extracting the PDZ core described in our previous work (Table S4 in Dudola et al. 2020), a structure alignment of the Shank1 PDZ domain and the PDB structure 1BE9, corresponding to PDZ3 of PSD95, was generated using MAMMOTH-Mult (Lupyan et al. 2005) and then the Cα atoms of residues matching with the core were extracted from all Shank1 PDZ structures in the MD snapshots. For the β2-β3 loop PCA, all full Shank1 PDZ structures were superimposed over all residues except the loop, then the Cα coordinates of all loop residues were extracted from the snapshots and the PCA was performed without another superposition step.

### Evaluation and visualization of data

DNA constructs were designed in SerialCloner 2.6.1. (RRID:SCR_014513). Sequence alignment was performed in BioEdit. (RRID:SCR_007361) (Hall, 1999) PCA modes were inspected in VMD. (Humphrey, Dalke, & Schulten, 1996) Other data were evaluated and visualized using Microsoft Excel (RRID:SCR_016137) or Gnuplot (RRID:SCR_008619). An Excel Solver add-in was used for sigmoid curve fitting of FP and CD measurements for figures. (Kemmer & Keller, 2010) Figures were edited in Microsoft Powerpoint.

## Supplementary Material

Besides figures and tables cited in the paper, we also provide the following supplementary files: *SupplementaryFigures.PDF* (Fig. S1-7), *SupplementaryTables_S1_experimental.xlsx* (Tables S1A-B), *SupplementaryTables_S2_computational.xlsx* (Tables S2A-K), as well a Zenodo repository containing raw experimental data and calculations from them (*Supplementary_CD_data.xlsx, Supplementary_FP_data.xlsx, Supplementary_TSA_data.xlsx*) and 90 MD simulation trajectories in the format of PDB ensembles, under the link https://zenodo.org/uploads/18662578.

## Author Contributions

**Anna Sánta:** Conceptualization, Investigation, Formal analysis, Writing - Original Draft, Writing - Review & Editing, Visualization. **Zsófia E. Kálmán:** Investigation, Formal analysis, Writing - Review & Editing. **Eszter Nagy-Kanta:** Investigation, Resources, Writing - Review & Editing. **Borbála Jakab:** Investigation, Writing - Review & Editing. **Zoltán Gáspári:** Writing - Review & Editing, Supervision, Funding acquisition. **Bálint Péterfia:** Conceptualization, Investigation, Resources, Writing - Original Draft, Writing - Review & Editing, Supervision.

## Acknowledgments

The authors acknowledge the support of the National Research, Development and Innovation Office - NKFIH - through grant OTKA K 137947 (to Z.G.), and the Digital Government Development and Project Management Ltd. for awarding us access to the Komondor HPC facility based in Hungary. The authors express gratitude for the contributions of: Enora Moutin for providing cDNA constructs, Viktor Farkas for the assistance with CD measurements, and Zsuzsanna Stránerné Szabó, Anna Oláh, József Hegedüs and Melinda Keresztes for their contribution to the experiments over the years.

## Abbreviations

ASD: autism spectrum disorder
BLI: biolayer interferometry
CD: circular dichroism
FP: fluorescence polarization
FT: flowthrough
GST: glutathione S-transferase
IEC: ion exchange chromatography
IMAC: immobilized metal ion chromatography
KI: knock-in
KO: knock-out
MD: molecular dynamics
PCA: principal component analysis
PBS: phosphate buffer saline
PDZ: PSD-95 (Postsynaptic Density protein 95), DLG1 (Drosophila Discs Large tumor suppressor protein), ZO-1 (Zona Occludens 1 protein) domain
PSD: postsynaptic density
RT: room temperature
SEC: size exclusion chromatography
SH3: SRC Homology 3 domain
SAM: Sterile Alpha Motif
TM: melting temperature
TSA: thermal shift assay

