## Supplemental Figures for "Redistribution of sidechain-sidechain interactions govern ligand-specific binding affinity changes in missense Shank1 PDZ variants"

A. Backbone RMSD values (Å) for the PDZ domain in the molecular dynamics runs for the various complexes (WT, V677M, R679W) 4

B. Backbone RMSD values (Å) for the PDZ domain in the molecular dynamics runs for the various complexes (G734S, R736Q, R743H) 5

C. Backbone RMSD values (Å) for the partner peptide in the molecular dynamics runs for the various complexes (WT, V677M, R679W) 6

D. Backbone RMSD values (Å) for the partner peptide in the molecular dynamics runs for the various complexes (G734S, R736Q, R743H) 7

Figure S2. 7

S3. Schematic representations of ligand binding ........ 8

A. GKAP 8

B. βPIX 9

C. mTOR 10

D. ARAP3 11

E. ATG13 12

F. 13

Figure S3. 13

S4. Analysis of the PDZ core ........ 14

A. PCA figures, mode 1-2, separated by peptide, colored by variant 14

B. Histograms of PCA mode 1 and mode 2, separated by peptide, colored by variant 15

C. PCA figures, mode 1-2, separated by variant, colored by peptide 16

D. Histograms of PCA mode 1 and mode 2, separated by variant, colored by peptide 17

E. PCA figures, mode 3-2, separated by peptide, colored by variant 18

F. Histograms of PCA mode 2 and mode 3, separated by peptide, colored by variant 19

G. PCA figures, mode 3-2, separated by variant, colored by peptide 20

H. Histograms of PCA mode 2 and mode 3, separated by variant, colored by peptide 21

Figure S4. 21

S5. Analysis of the β2-β3 loop........................................................................ 22

A. RMSF along the 680-700 segment 22

B. Helical propensity of the 680-690 segment 23

C. Cumulative explained variance of PCA modes 23

D. PCA figures, mode 1-2, separated by peptide, colored by variant 24

E. Histograms of PCA mode 1 and mode 2, separated by peptide, colored by variant 25

F. PCA figures, mode 1-2, separated by variant, colored by peptide 26

G. Histograms of PCA mode 1 and mode 2, separated by variant, colored by peptide 27

H. PCA figures, mode 3-2, separated by peptide, colored by variant 28

I. Histograms of PCA mode 3 and mode 2, separated by peptide, colored by variant 29

J. PCA figures, mode 3-2, separated by variant, colored by peptide 30

K. Histograms of PCA mode 3 and mode 2, separated by variant, colored by peptide 31

Figure S5. 32

S6. H735 *X* angle distributions........................................................................ 33

A. GKAP 33

B. mTOR 34

C. ARAP3 35

Figure S6. 35

### S1. Unedited SDS-PAGE image

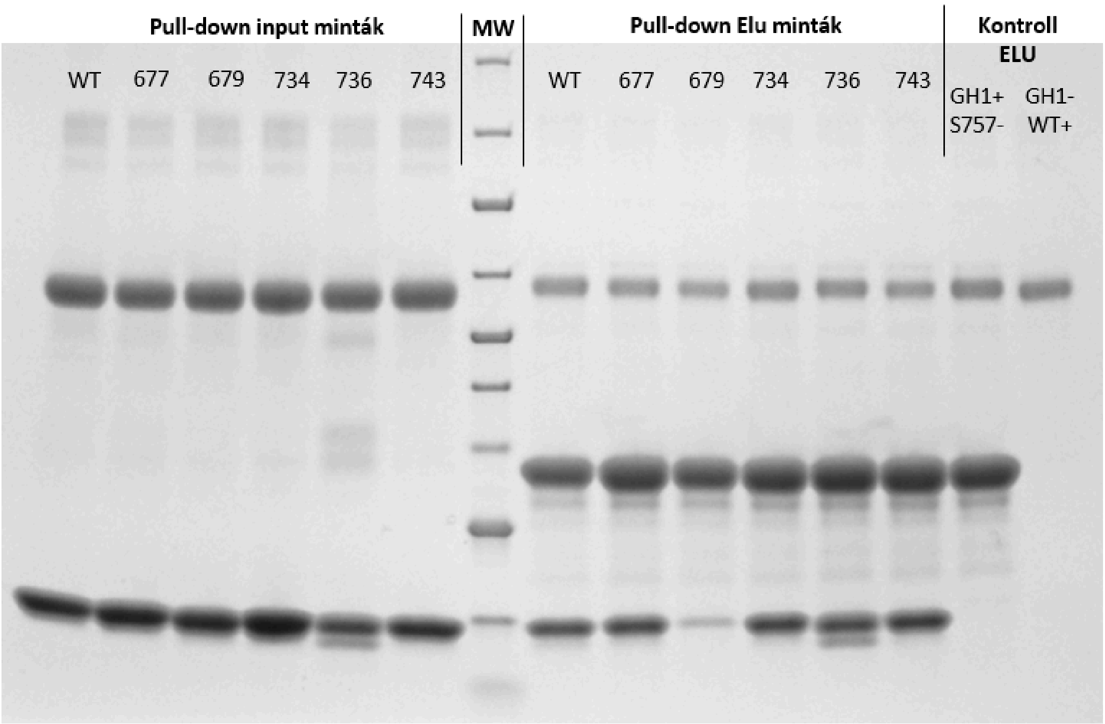

**Figure S1.**

The SDS-PAGE image from Fig. 3A with its original text. The incomplete mutant name labels and Hungarian-language text were saved onto the original image and therefore needed to be removed via editing to be covered with the final text used in the figure. We provide this image for transparency, to assure that no photo editing tools were used on parts of the image where samples are visible.

### S2. Fixed domain-peptide RMSD plots

A. Backbone RMSD values (Å) for the PDZ domain in the molecular dynamics runs for the various complexes (WT, V677M, R679W)

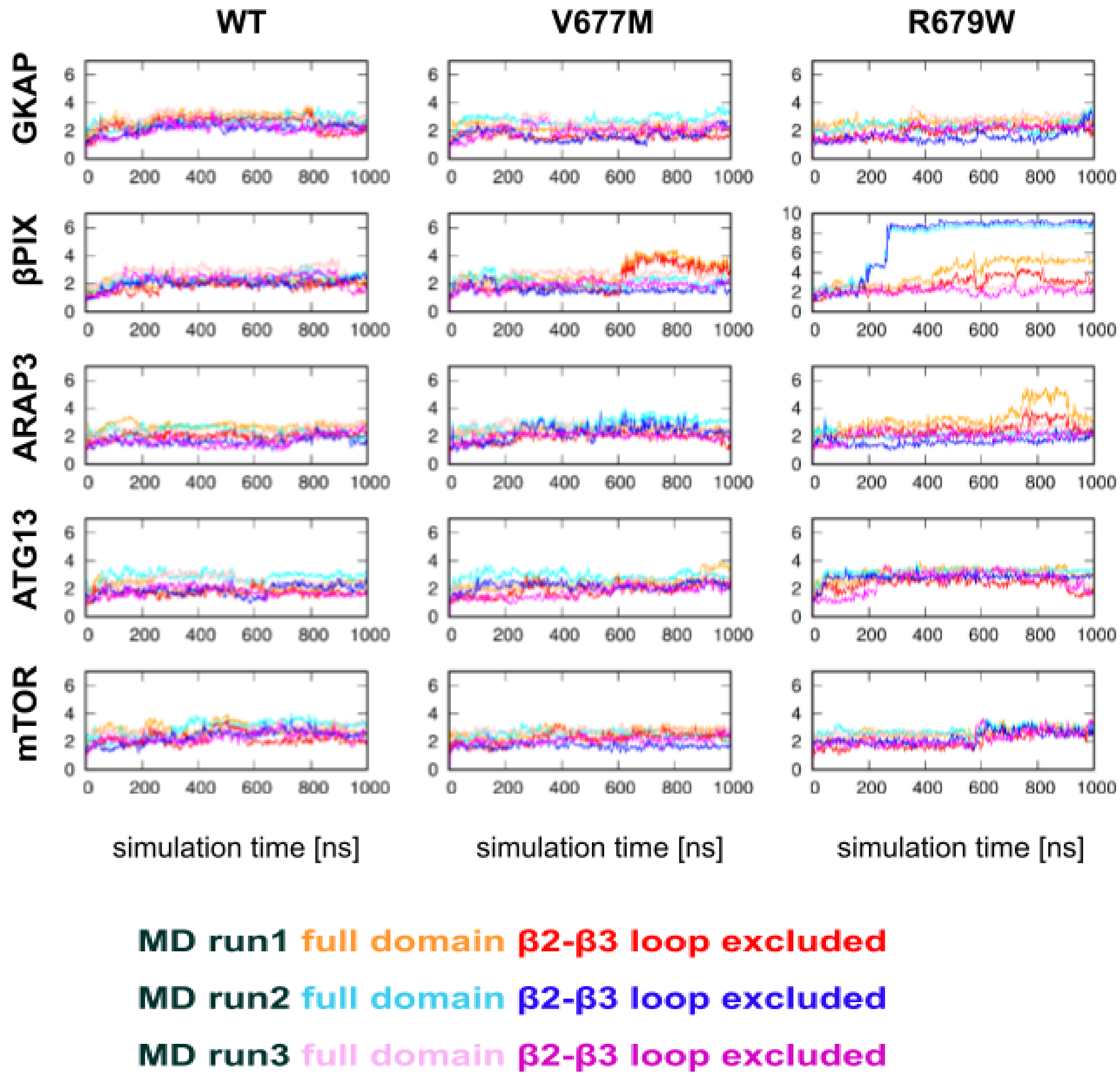

#### B. Backbone RMSD values (Å) for the PDZ domain in the molecular dynamics runs for the various complexes (G734S, R736Q, R743H)

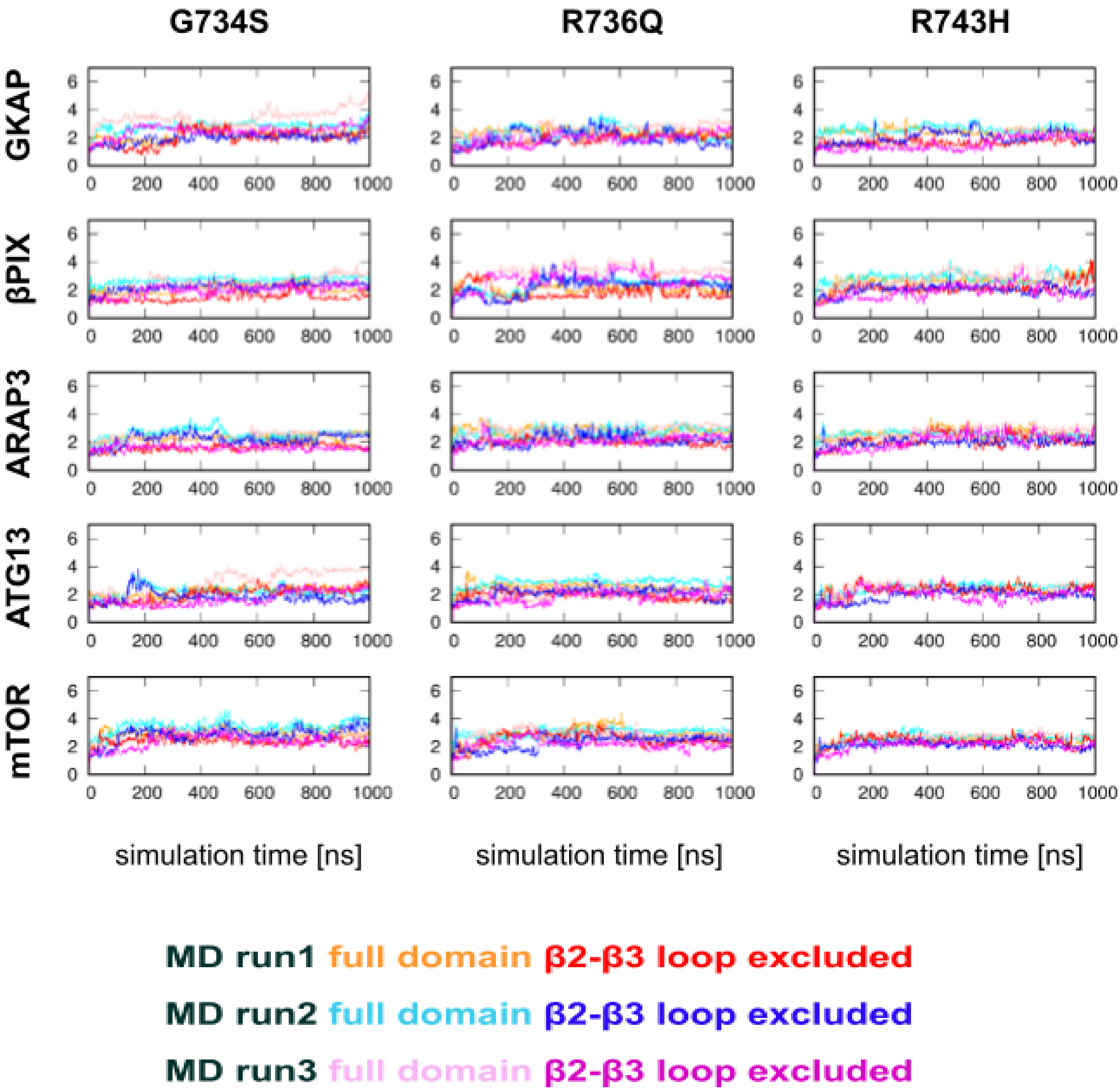

C. Backbone RMSD values (Å) for the partner peptide in the molecular dynamics runs for the various complexes (WT, V677M, R679W)

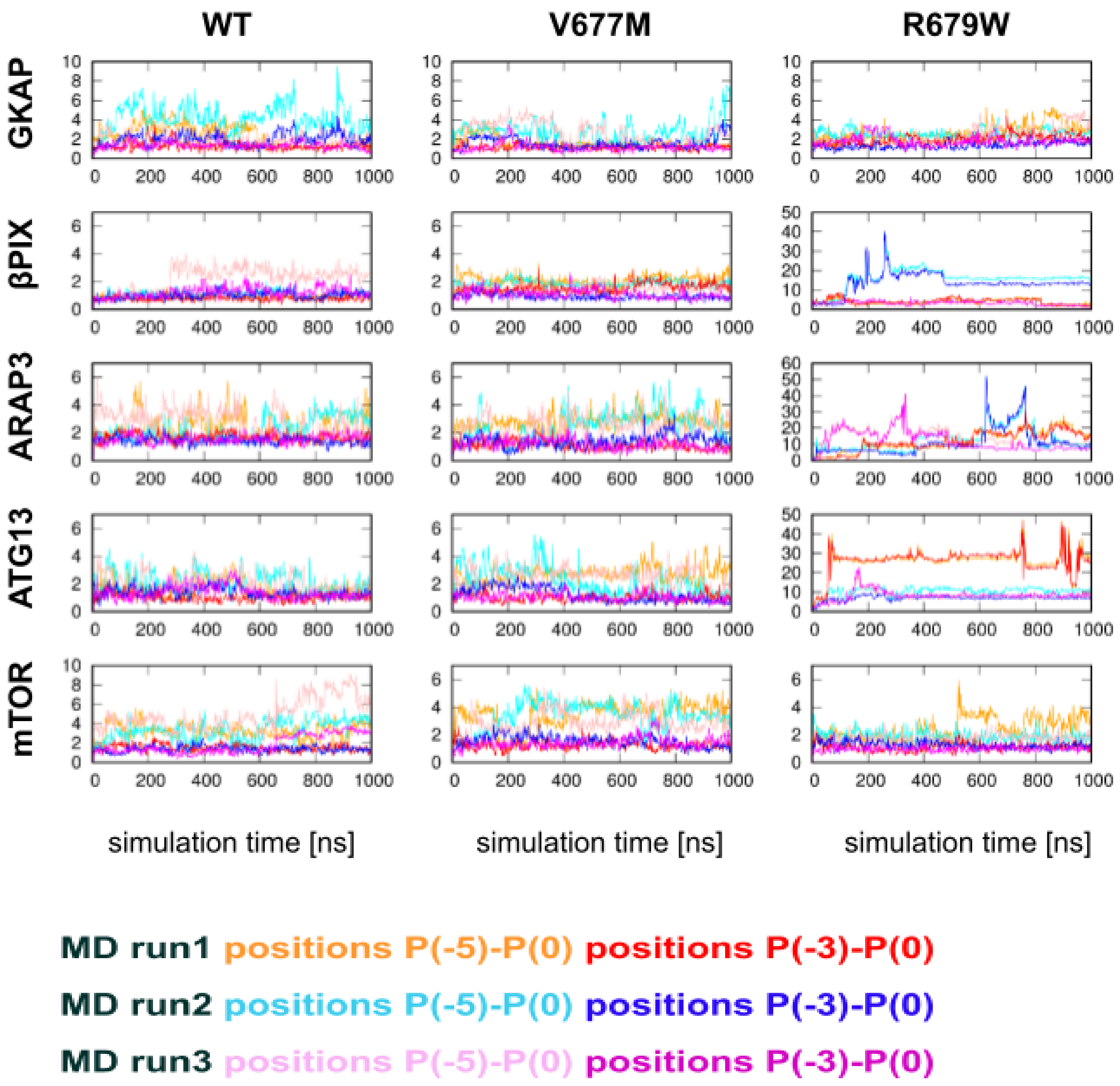

#### D. Backbone RMSD values (Å) for the partner peptide in the molecular dynamics runs for the various complexes (G734S, R736Q, R743H)

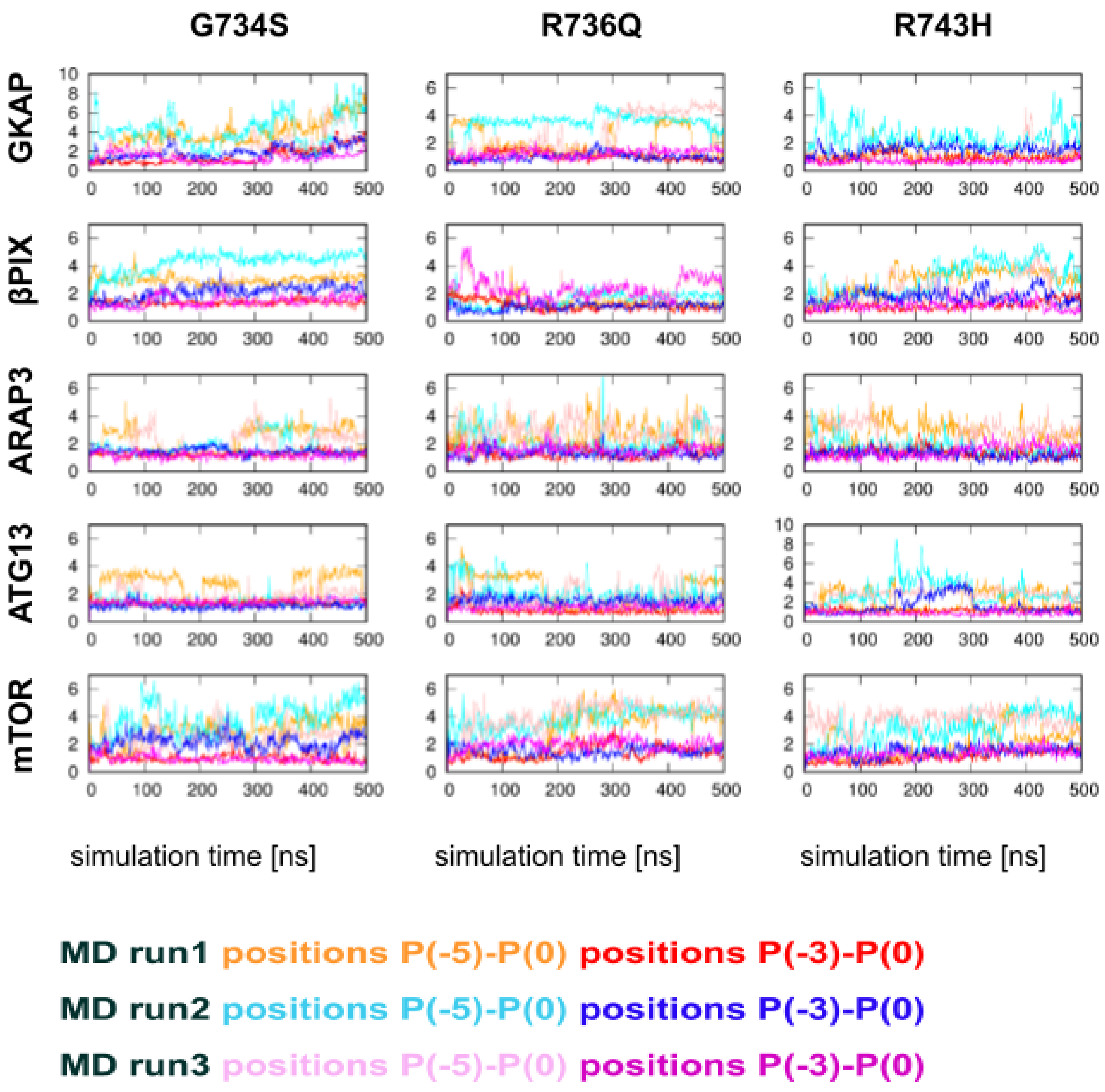

**Figure S2.**

**(A-D)** Backbone RMSD values (Å) through the molecular dynamics simulations. All reported RMSD values are calculated from structures superimposed to the first frame, using all residues in the PDZ domain only. RMSD calculations were performed for different backbone atom sets for the PDZ domain (full domain and β2-β3 loop excluded) and the peptide (all positions and C-terminal residues) as indicated by the color code. The backbone atoms considered are: Cα, C, N. Note that for some complexes, the y axis scale is different to account for the higher variability of domain conformations / peptide poses, especially for the highly destabilized R679W variant, for which the dissociation of the peptide has also been observed in some cases.

### S3. Schematic representations of ligand binding

#### A. GKAP

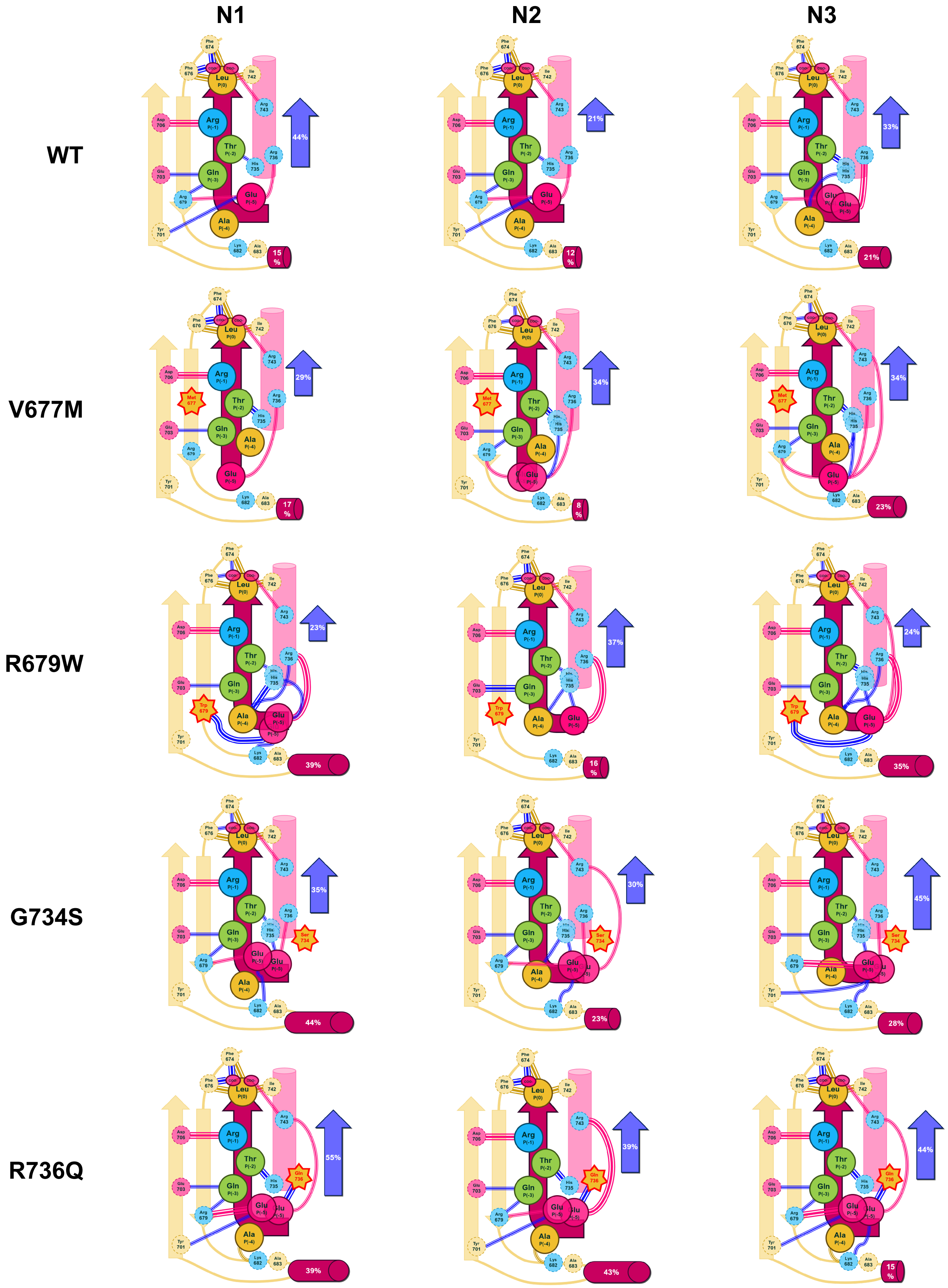

**R743H**

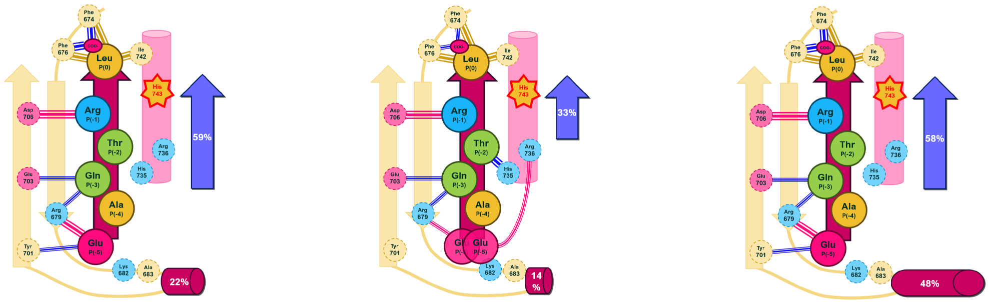

#### B. βPIX

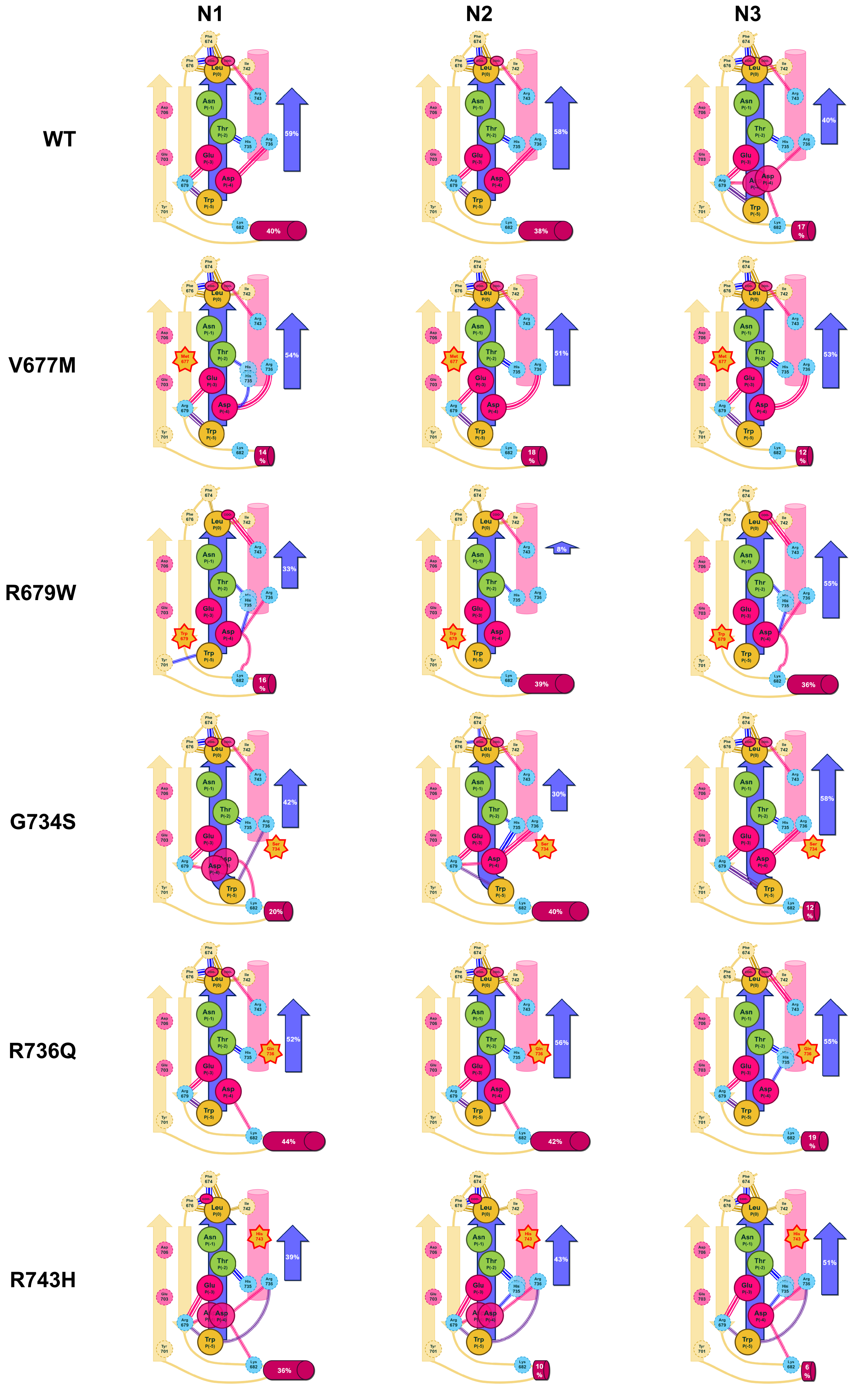

#### C. mTOR

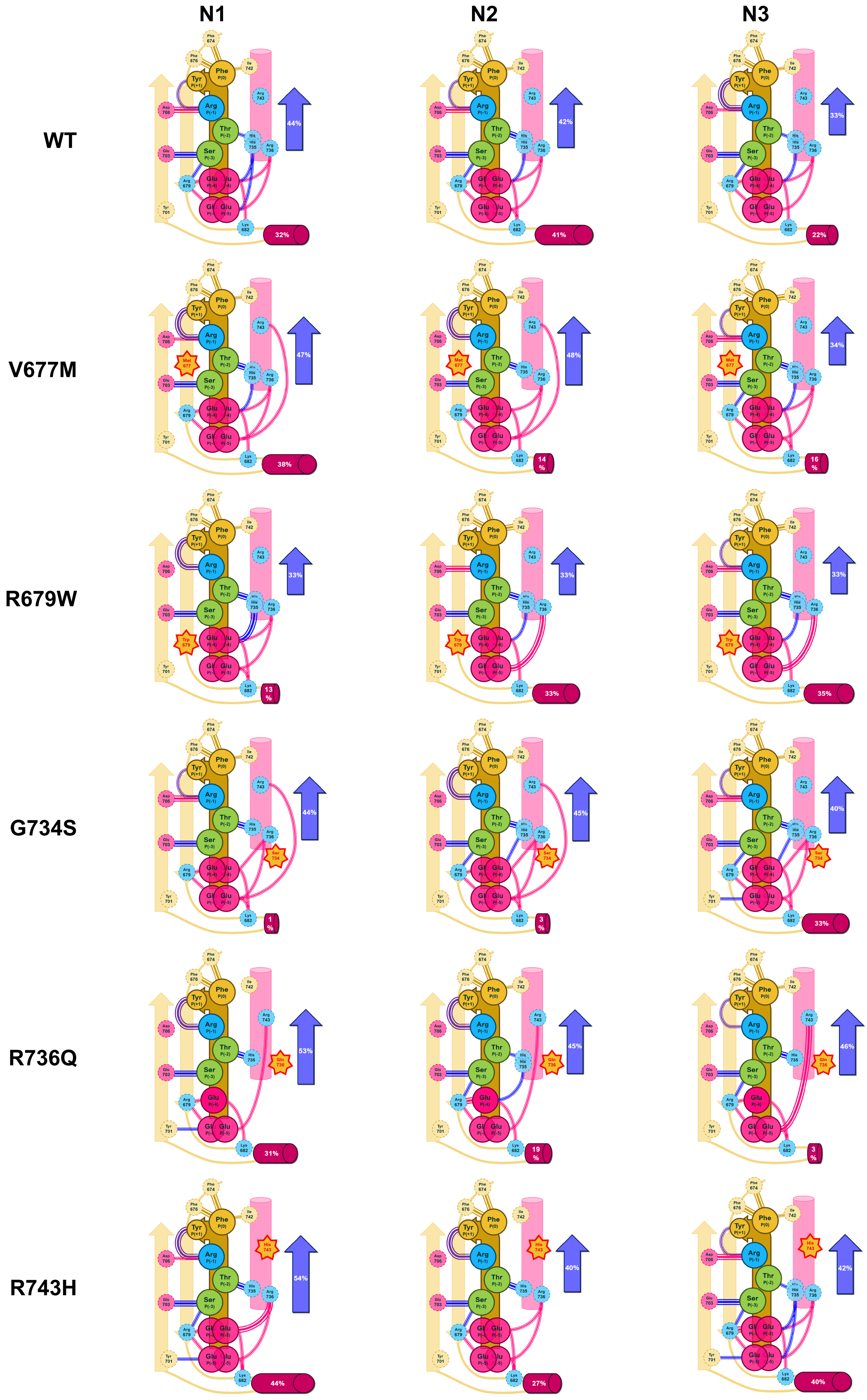
N1
N2
N3
WT
V677M
R679W
G734S
R736Q
R743H
44%
32%
42%
41%
33%
22%
47%
38%
48%
14 %
34%
16 %
33%
13 %
33%
33%
33%
35%
44%
1 %
45%
3 %
40%
33%
53%
31%
45%
19 %
46%
3 %
54%
44%
40%
27%
42%
40%

#### D. ARAP3

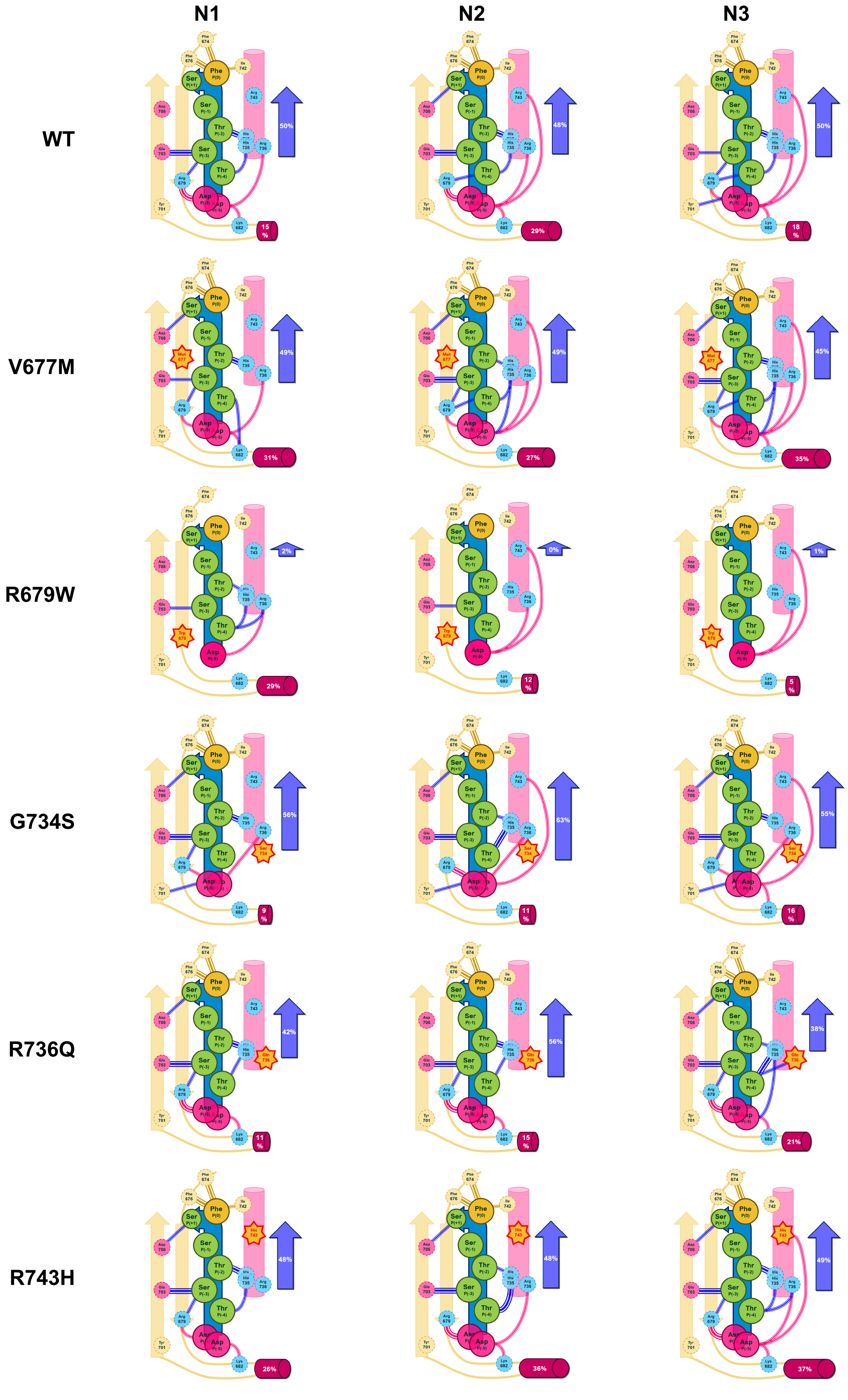
N1
N2
N3
WT
50%
15%
48%
29%
50%
18%
V677M
49%
31%
49%
27%
45%
35%
R679W
2%
29%
0%
12%
1%
5%
G734S
56%
9%
63%
11%
55%
16%
R736Q
42%
11%
56%
15%
38%
21%
R743H
48%
26%
48%
36%
49%
37%

#### E. ATG13

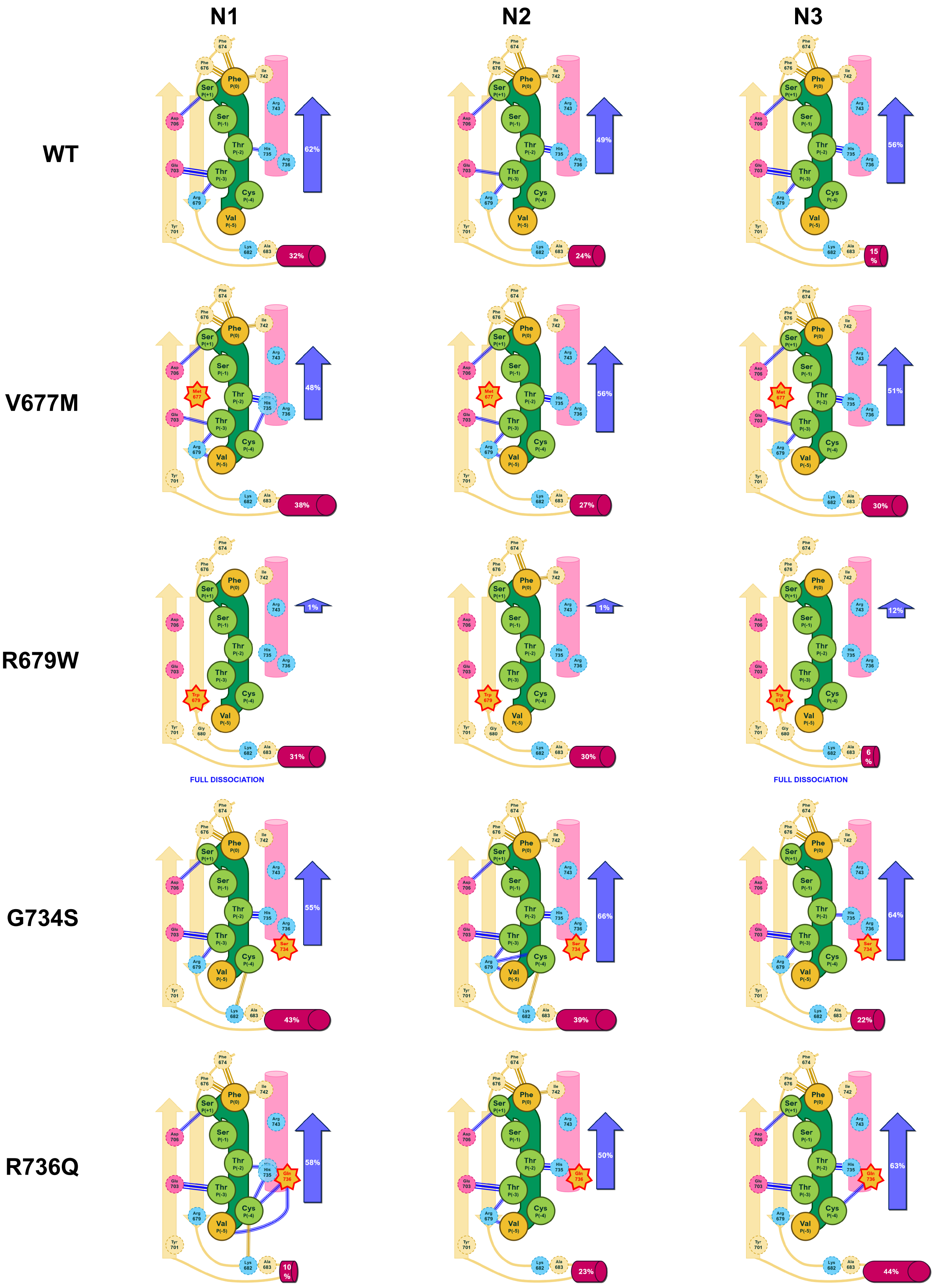

N1
N2
N3
WT
62%
32%
49%
24%
56%
15 %
V677M
48%
38%
56%
27%
51%
30%
R679W
1%
31%
FULL DISSOCIATION
1%
30%
12%
6 %
FULL DISSOCIATION
G734S
55%
43%
66%
39%
64%
22%
R736Q
58%
10 %
50%
23%
63%
44%

R743H

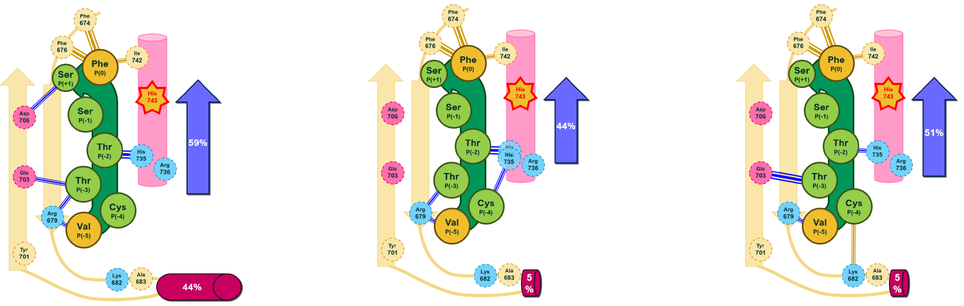

**F.**

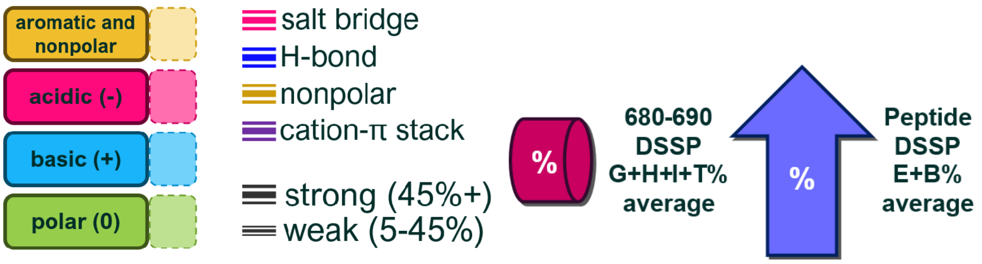

#### Figure S3.

**(A-E)** Schematic representations of residue:residue interactions observed in the MD runs (N1-3) of the different complexes. Types of the interactions are color-coded as indicated. Prevalence of helical and extended secondary structures for the β2- β3 loop and the bound peptide are also indicated, respectively. **(F)**

### S4. Analysis of the PDZ core

#### A. PCA figures, mode 1-2, separated by peptide, colored by variant

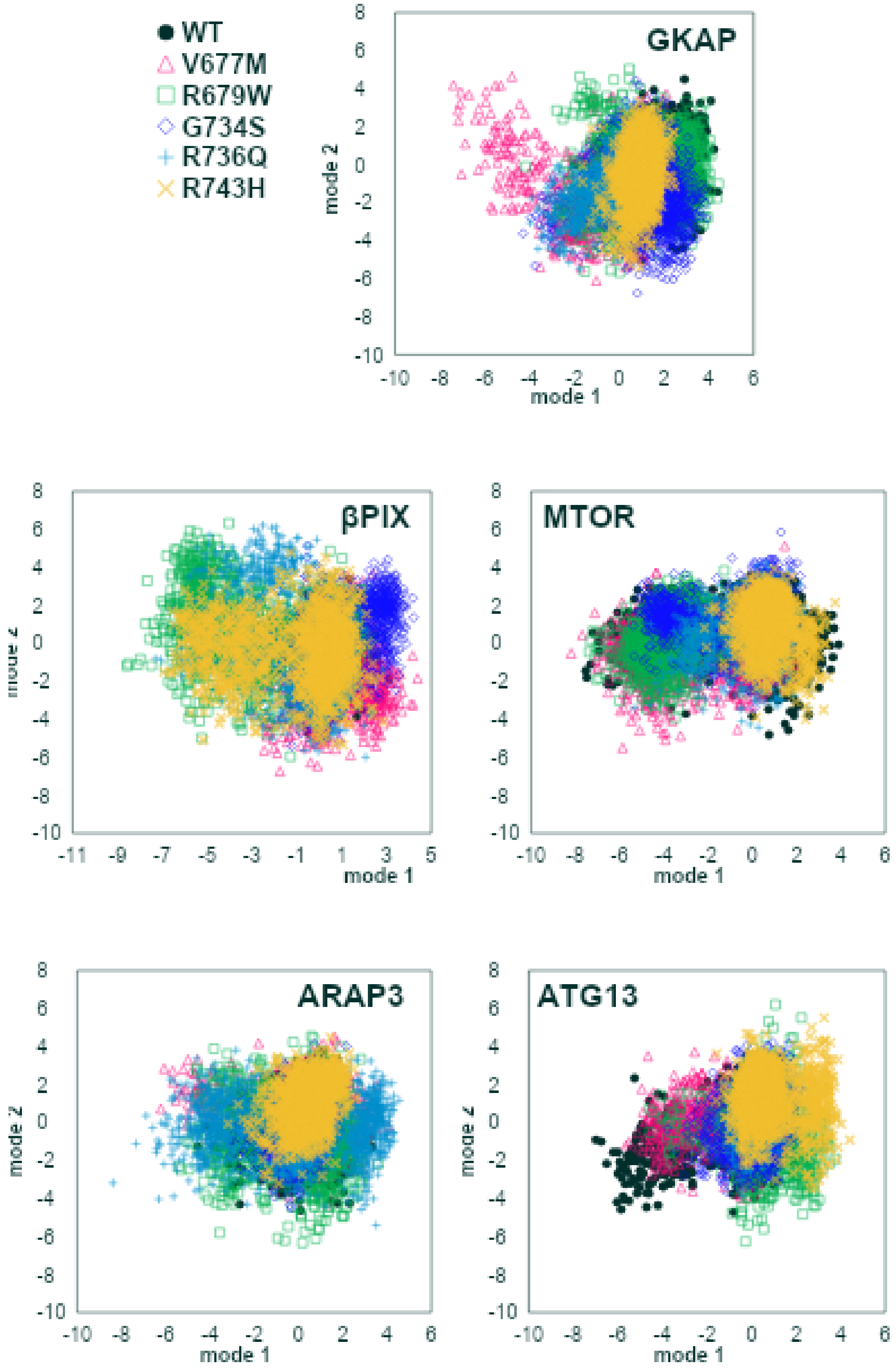

B. Histograms of PCA mode 1 and mode 2, separated by peptide, colored by variant

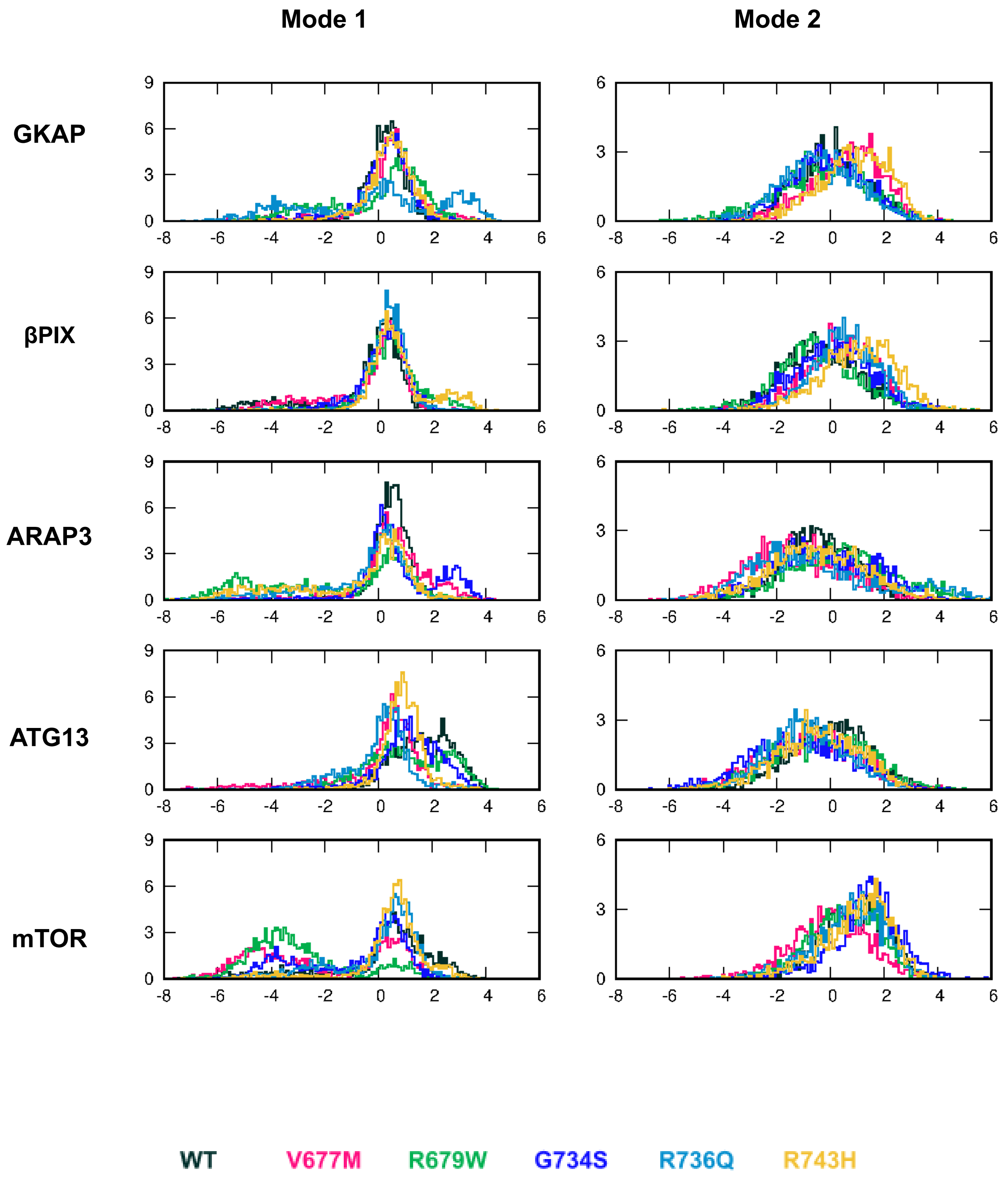

#### C. PCA figures, mode 1-2, separated by variant, colored by peptide

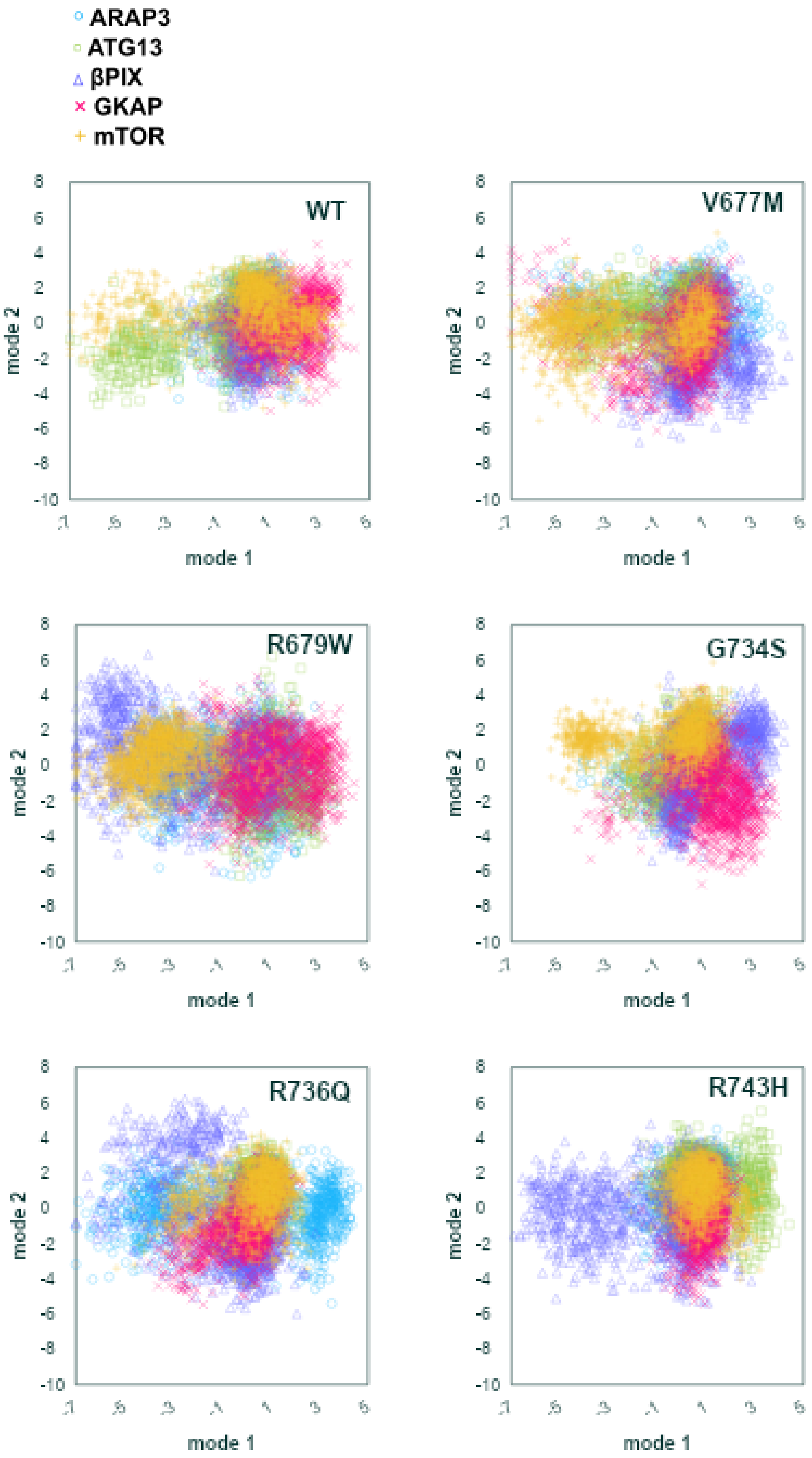

#### D. Histograms of PCA mode 1 and mode 2, separated by variant, colored by peptide

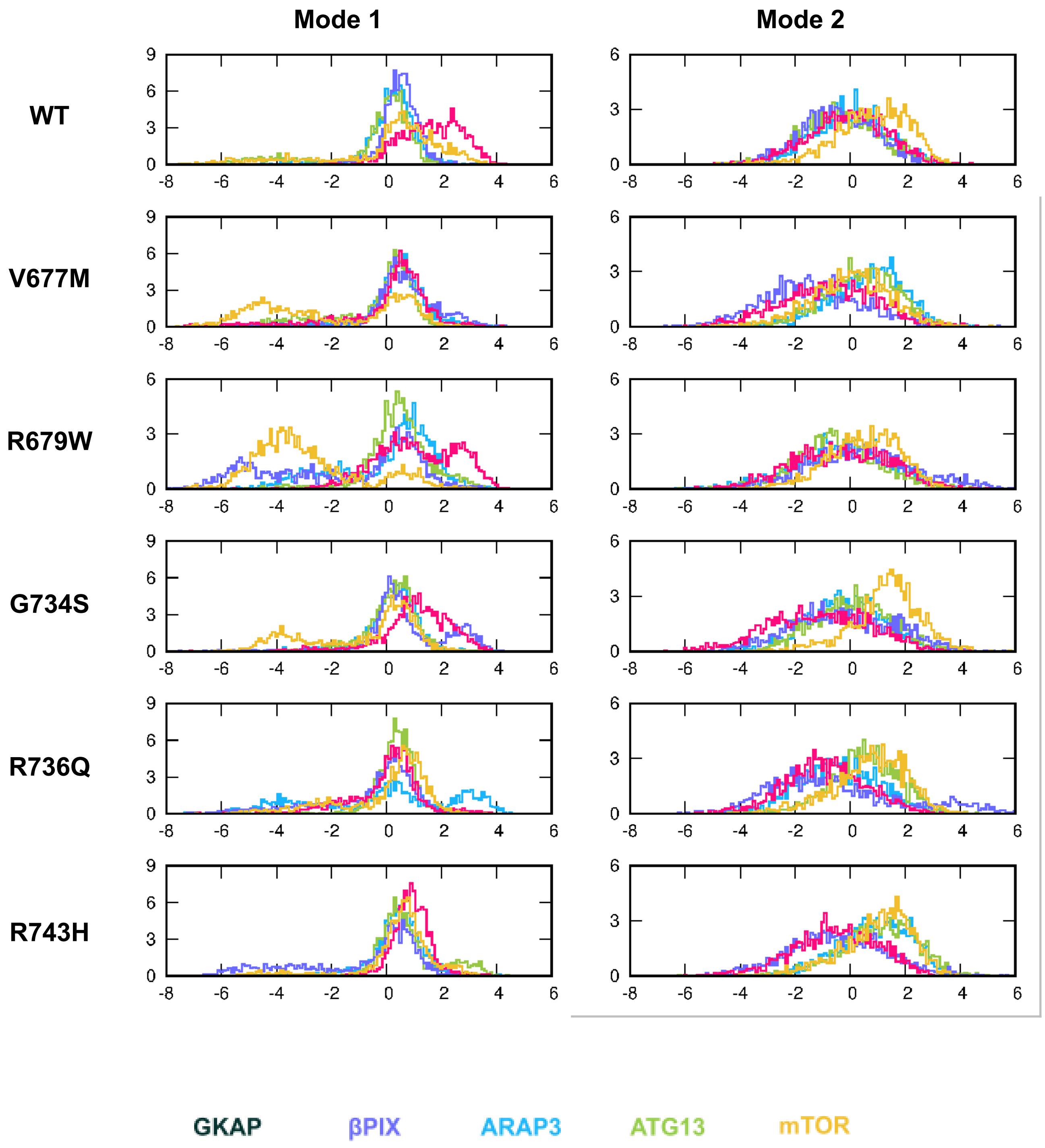

#### E. PCA figures, mode 3-2, separated by peptide, colored by variant

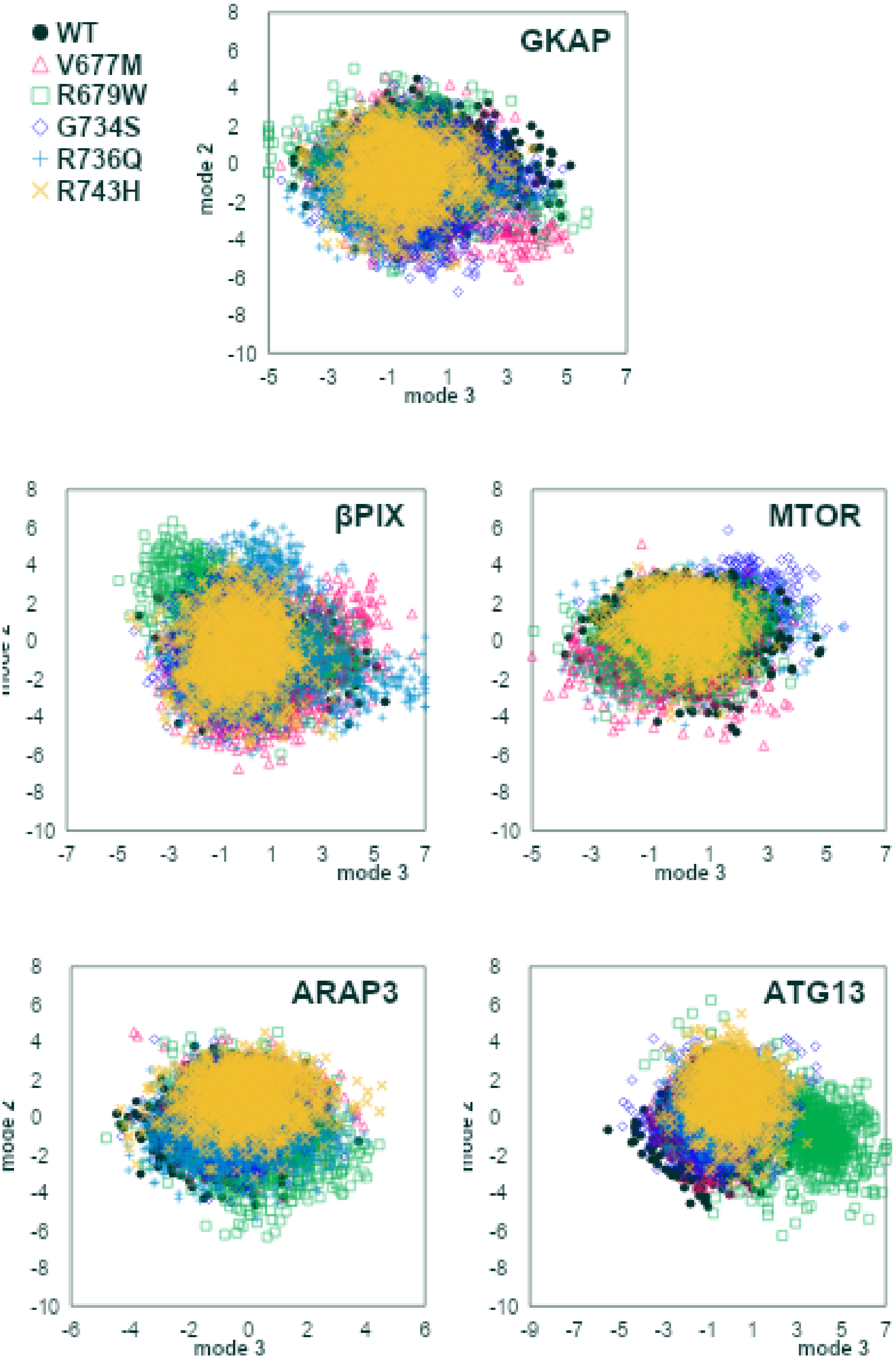

#### F. Histograms of PCA mode 2 and mode 3, separated by peptide, colored by variant

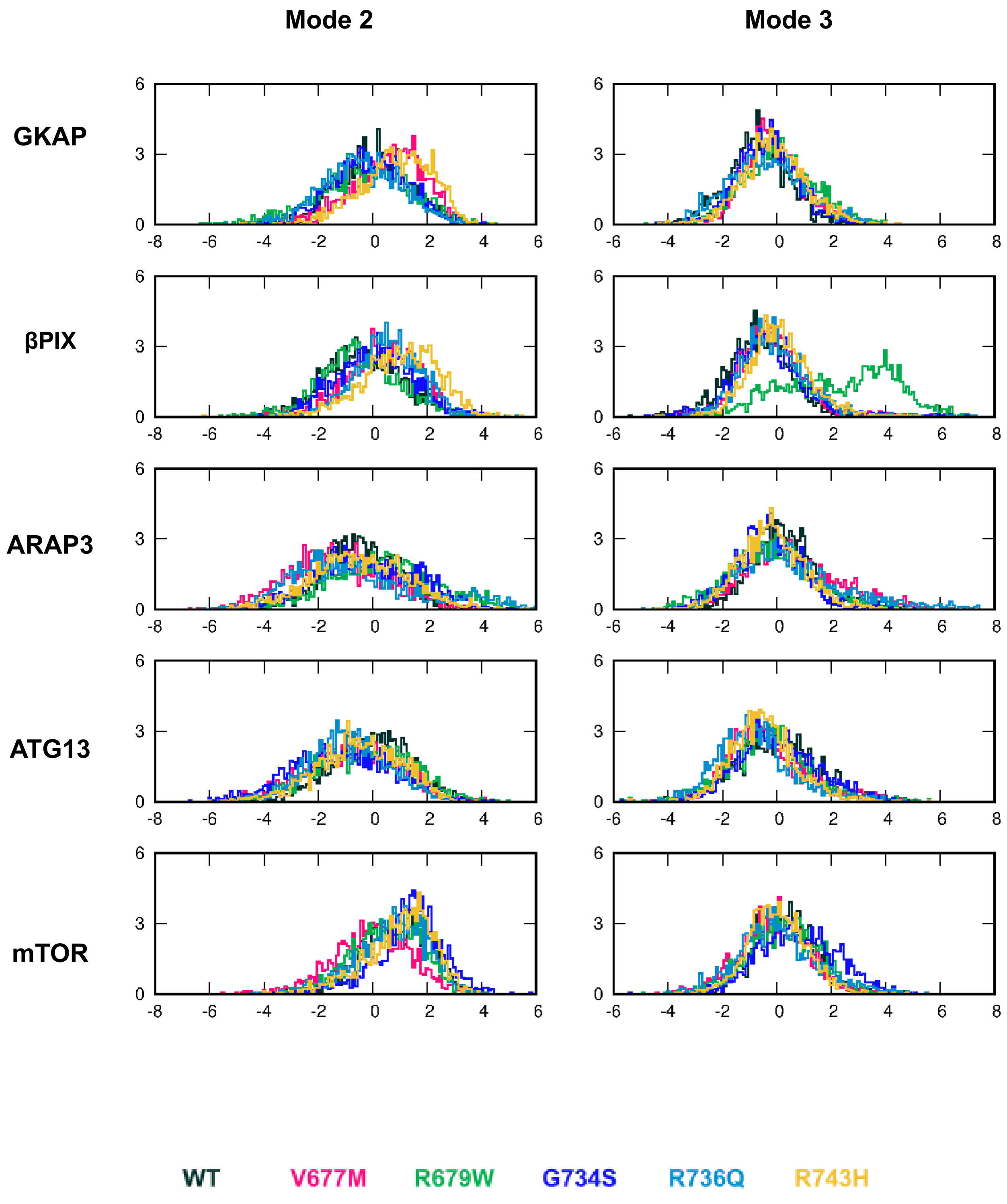

G. PCA figures, mode 3-2, separated by variant, colored by peptide

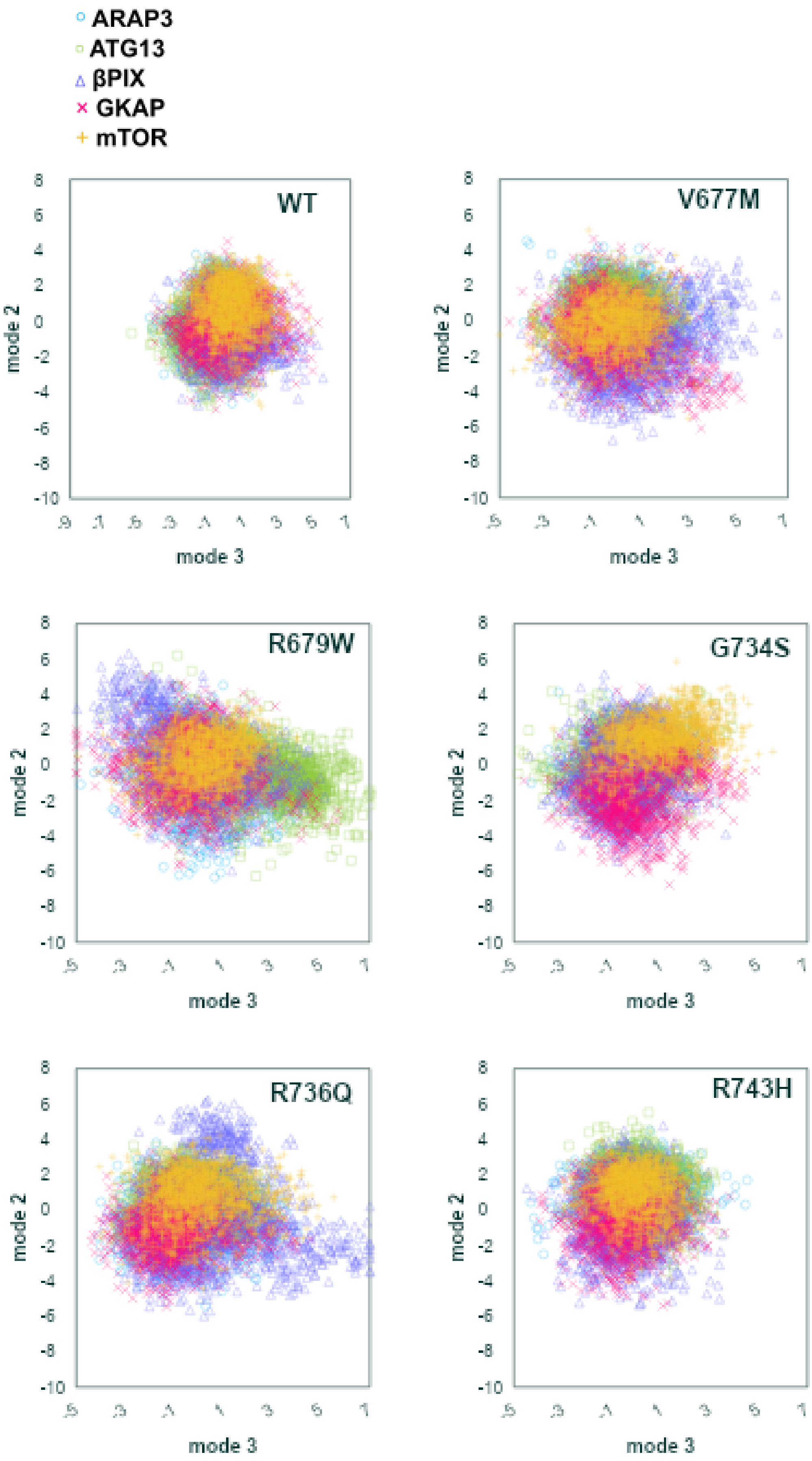

#### H. Histograms of PCA mode 2 and mode 3, separated by variant, colored by peptide

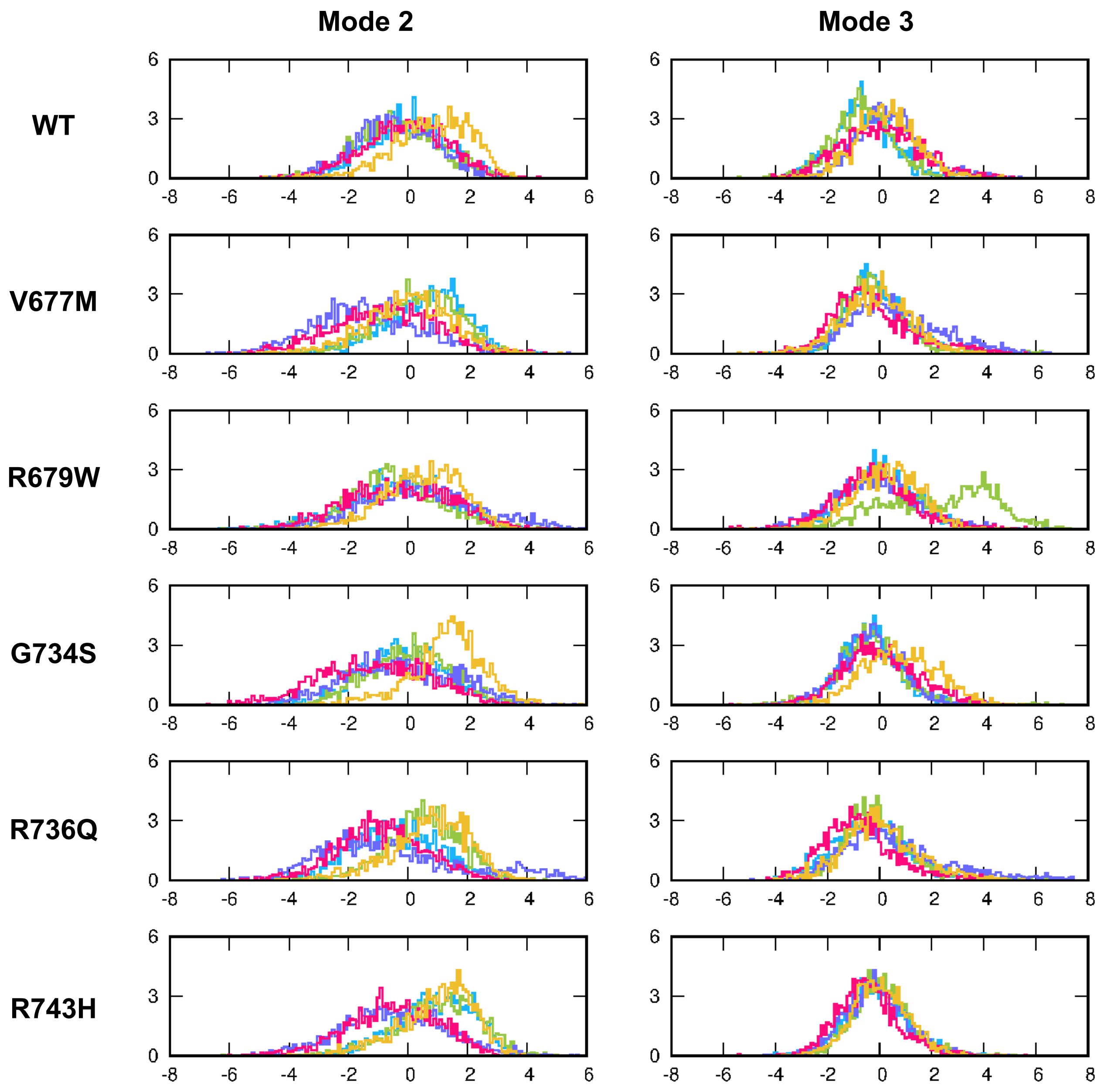

**Figure S4.** GKAP βPIX ARAP3 ATG13 mTOR

Mode 1, 2 and 3 of the PCA on the PDZ core are plotted against each other in **(A)**, **(C)**, **(E)** and **(G)** separated into individual plots by variant and peptide. Histograms showing distributions in mode occupancies are included in **(B)**, **(D)**, **(F)** and **(H)**.

### S5. Analysis of the β2-β3 loop

#### A. RMSF along the 680-700 segment

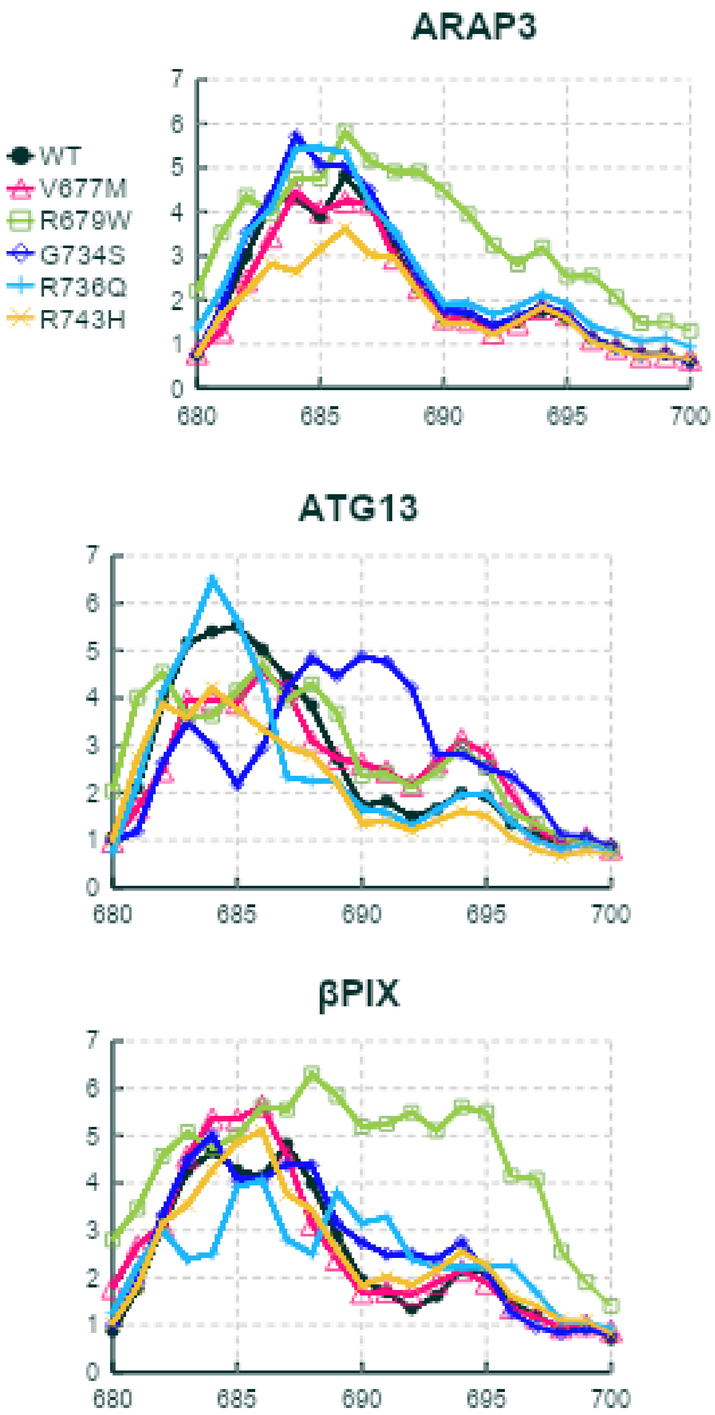

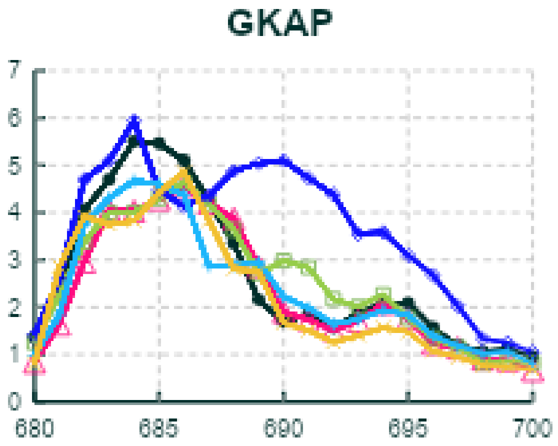
GKAP
7
6
5
4
3
2
1
0
680
685
690
695
700

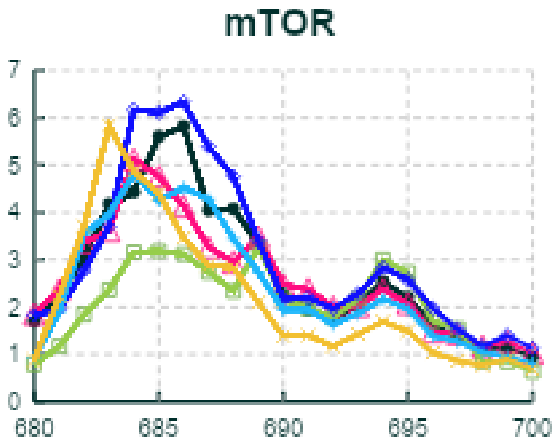
mTOR
7
6
5
4
3
2
1
0
680
685
690
695
700

#### B. Helical propensity of the 680-690 segment

| | GKAP | | | βPIX | | | mTOR | | | ARAP3 | | | ATG13 | | |
|---|---|---|---|---|---|---|---|---|---|---|---|---|---|---|---|
| | N1 | N2 | N3 | N1 | N2 | N3 | N1 | N2 | N3 | N1 | N2 | N3 | N1 | N2 | N3 |
| **WT** | 15% | 12% | 21% | 40% | 38% | 17% | 32% | 41% | 22% | 15% | 29% | 18% | 32% | 24% | 15% |
| **V677M** | 17% | 8% | 23% | 14% | 18% | 12% | 38% | 14% | 16% | 31% | 27% | 35% | 38% | 27% | 30% |
| **R679W** | 39% | 16% | 35% | 16% | 39% | 36% | 13% | 33% | 35% | 29% | 12% | 5% | 31% | 30% | 6% |
| **G734S** | 44% | 23% | 28% | 20% | 40% | 12% | 1% | 3% | 33% | 9% | 11% | 16% | 43% | 39% | 22% |
| **R736Q** | 39% | 43% | 15% | 44% | 42% | 19% | 31% | 19% | 3% | 11% | 15% | 21% | 10% | 23% | 44% |
| **R743H** | 22% | 14% | 48% | 36% | 10% | 6% | 44% | 27% | 40% | 26% | 36% | 37% | 44% | 5% | 5% |

#### C. Cumulative explained variance of PCA modes

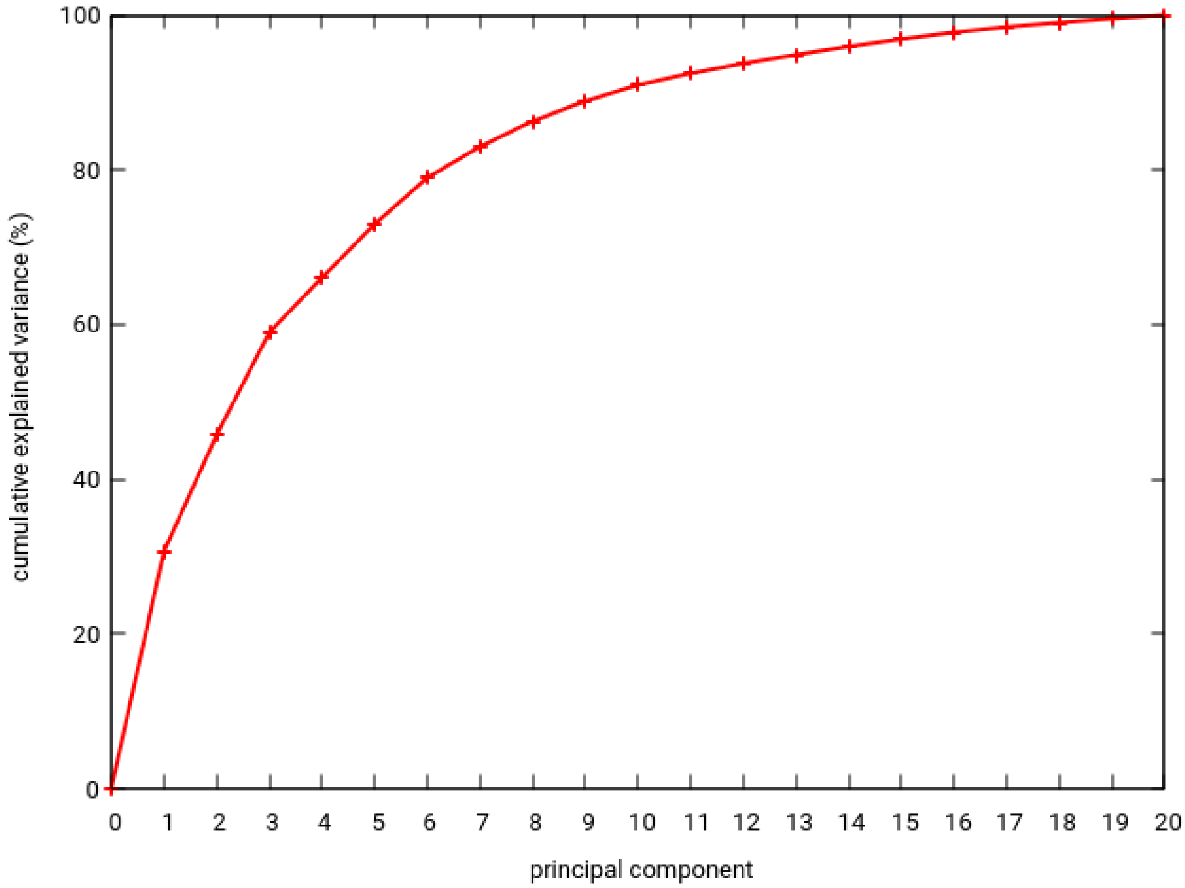

#### D. PCA figures, mode 1-2, separated by peptide, colored by variant

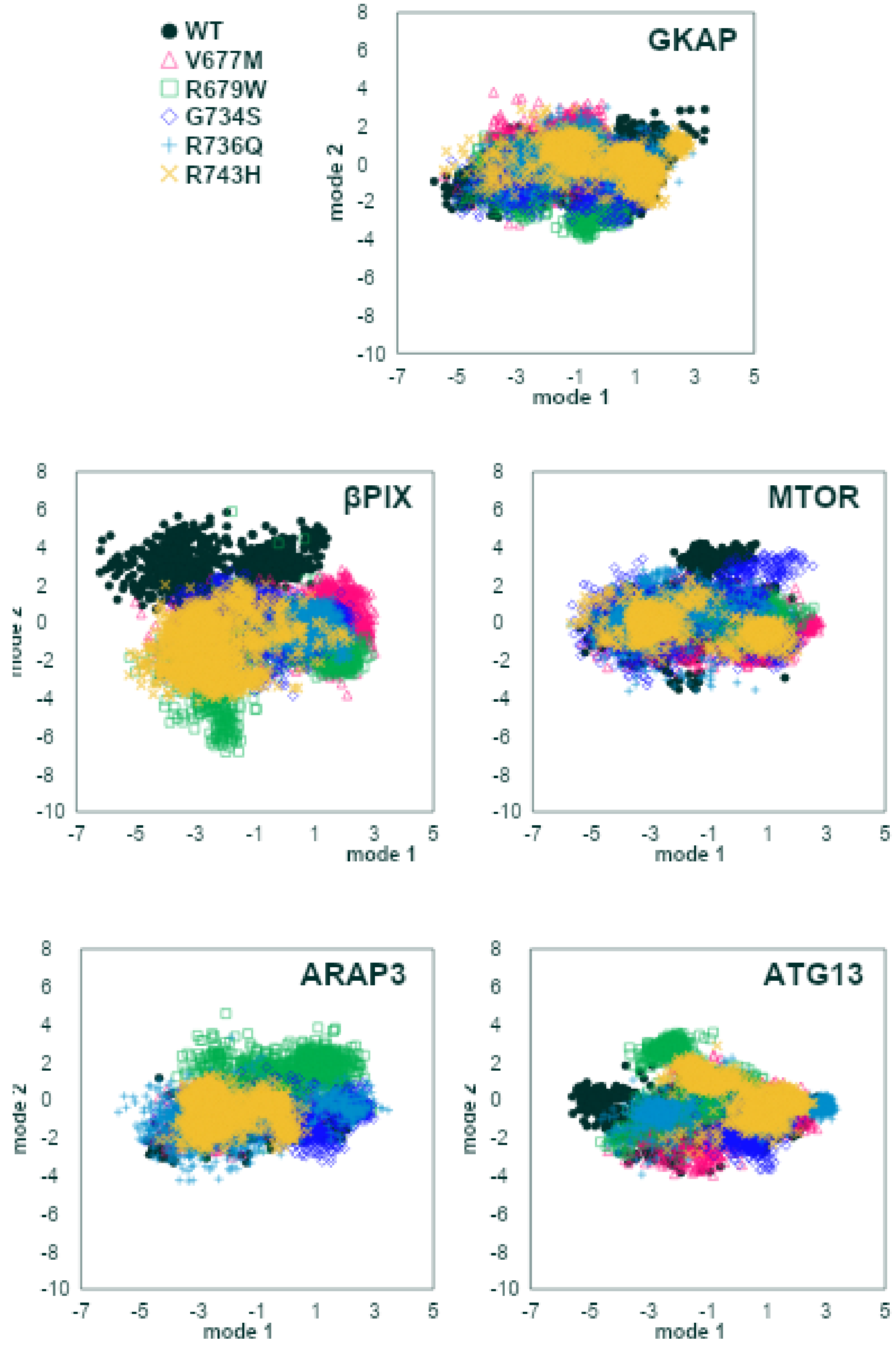

#### E. Histograms of PCA mode 1 and mode 2, separated by peptide, colored by variant

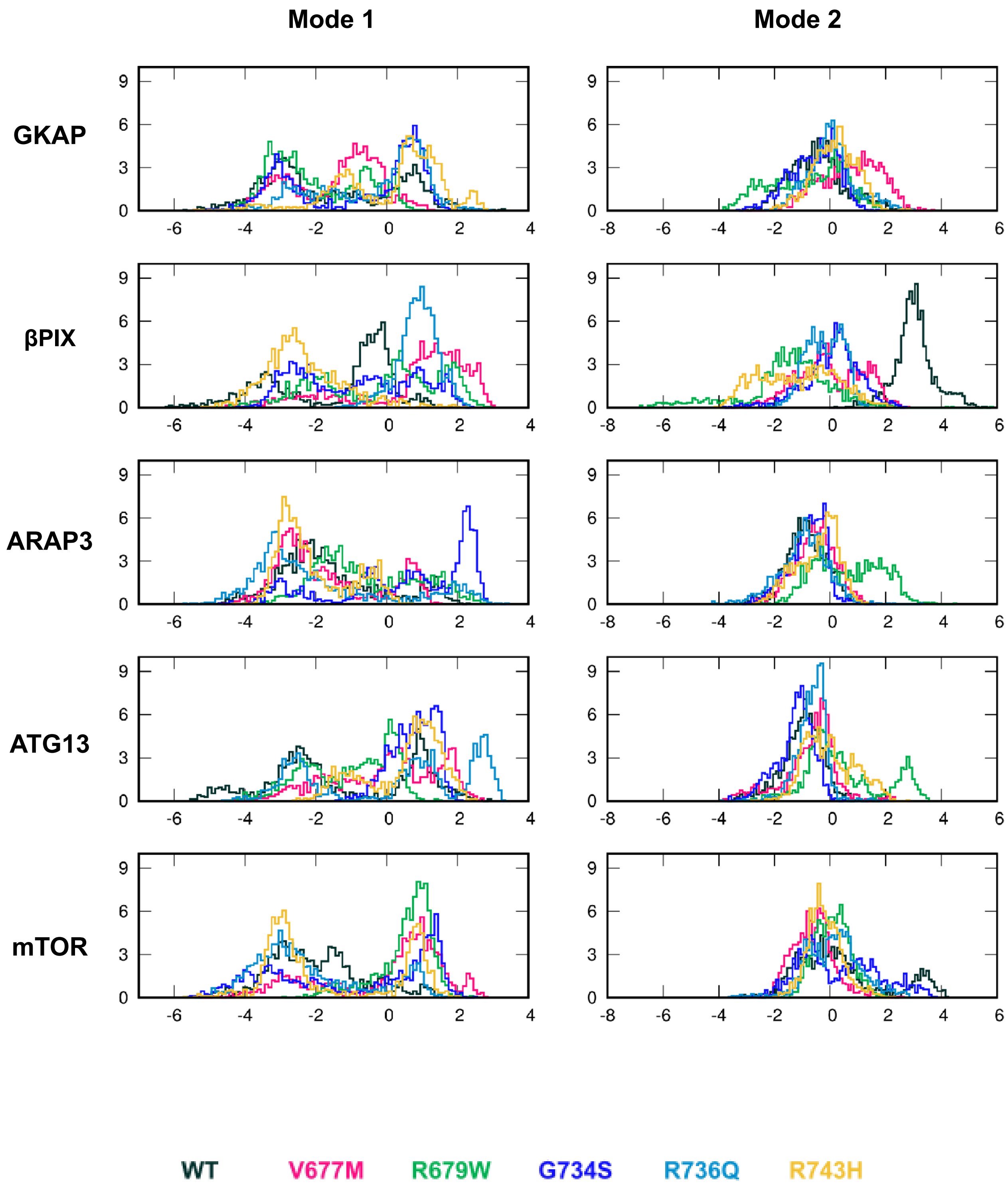

F. PCA figures, mode 1-2, separated by variant, colored by peptide

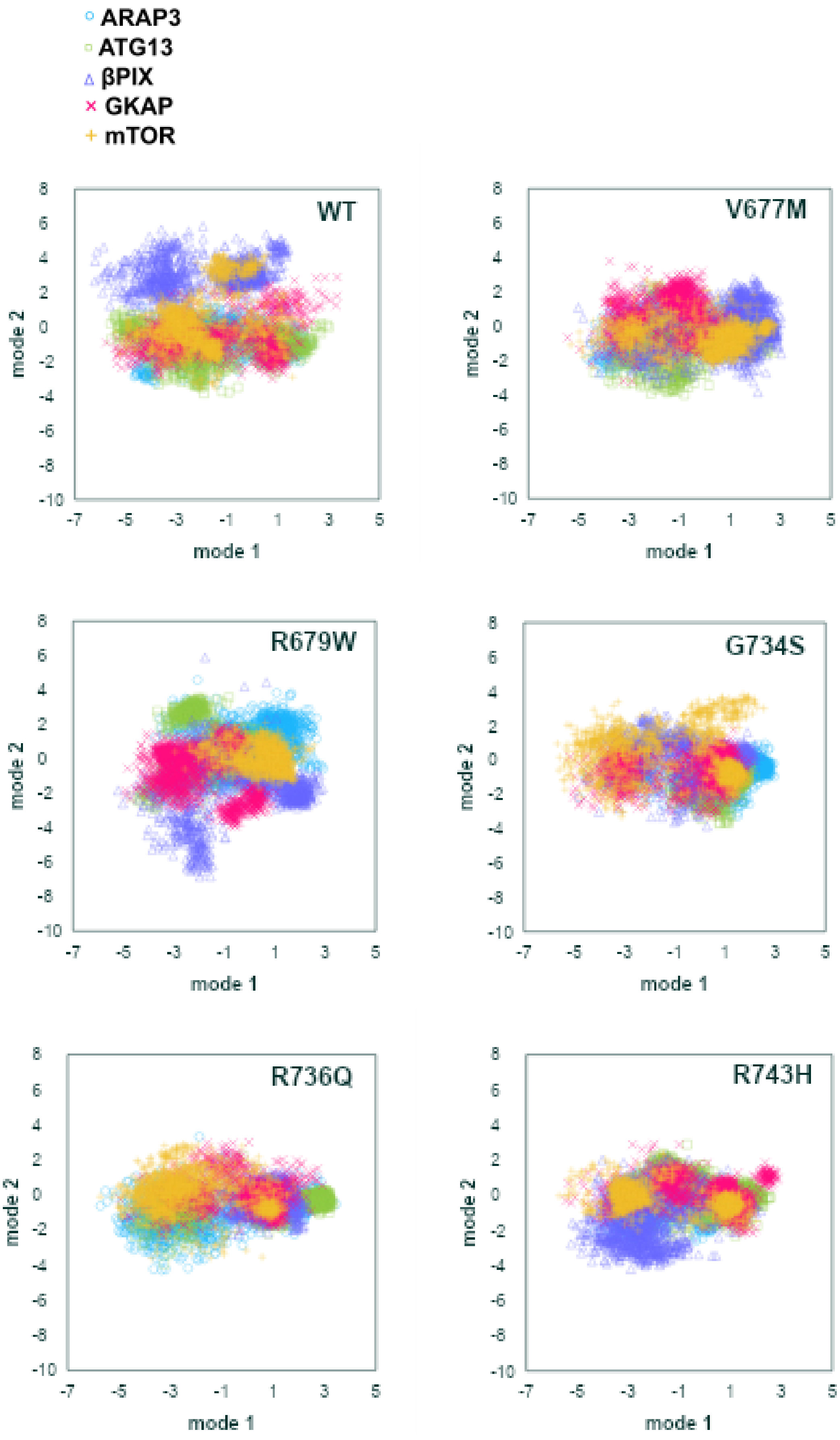

#### G. Histograms of PCA mode 1 and mode 2, separated by variant, colored by peptide

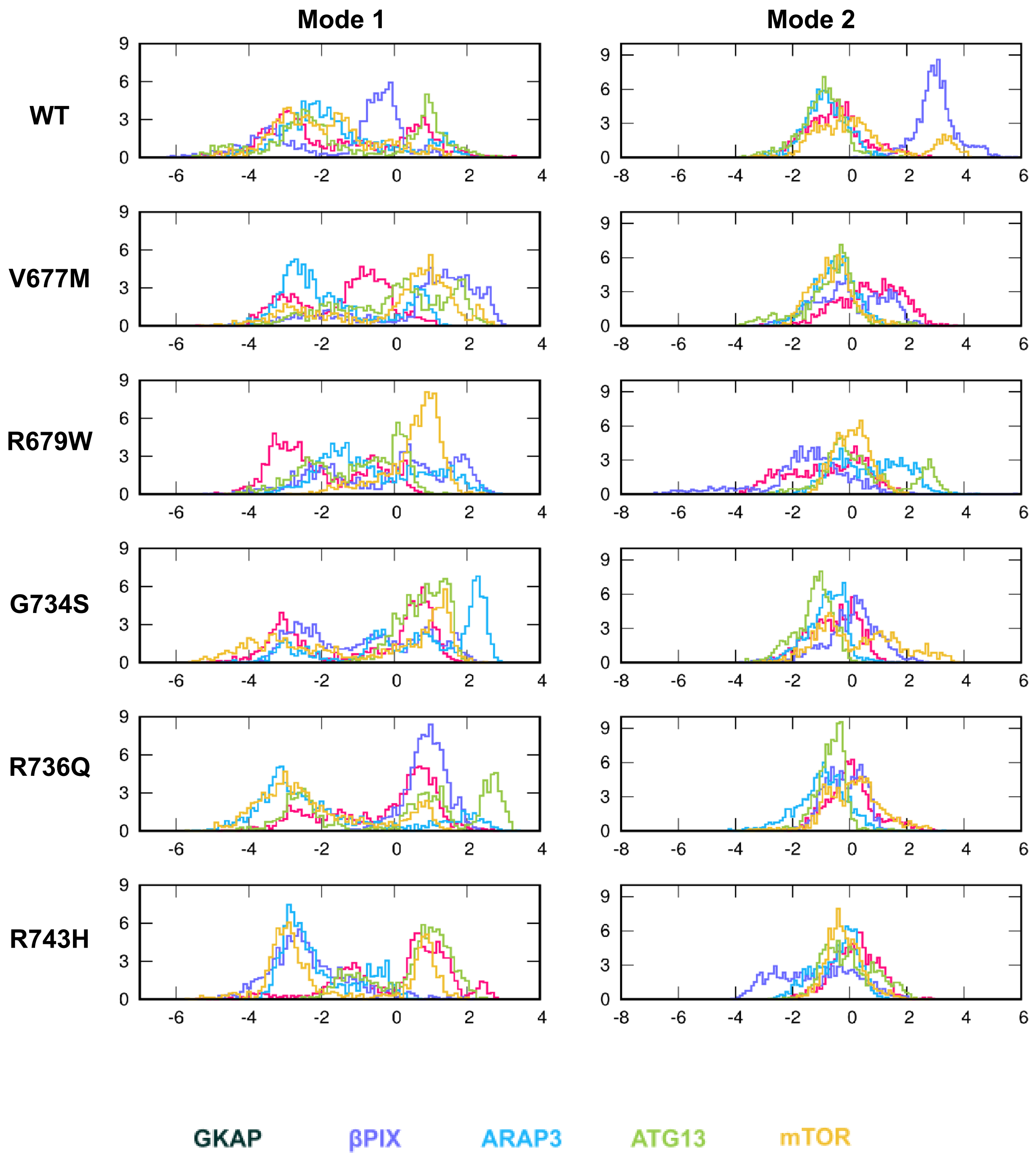

#### H. PCA figures, mode 3-2, separated by peptide, colored by variant

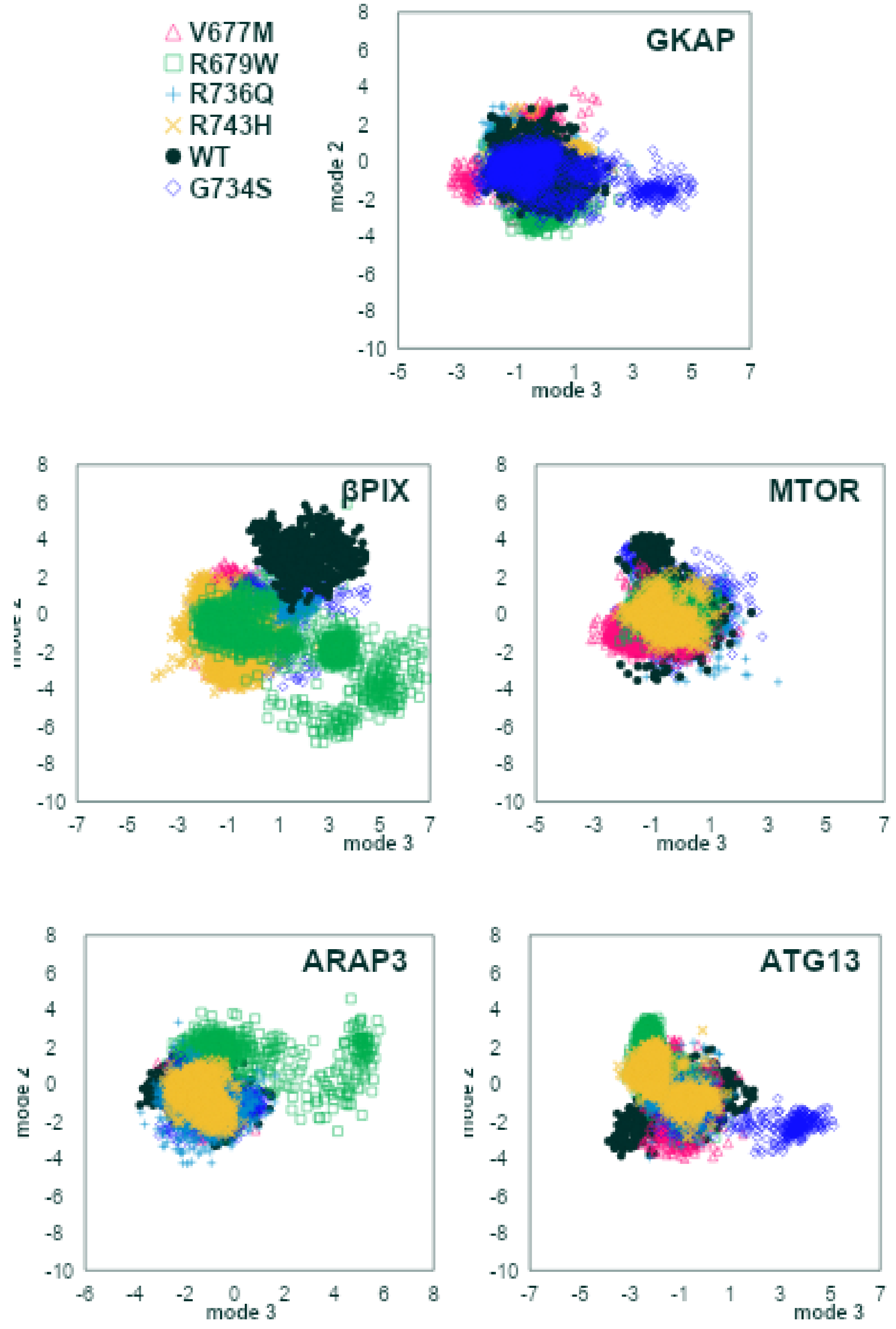

#### I. Histograms of PCA mode 3 and mode 2, separated by peptide, colored by variant

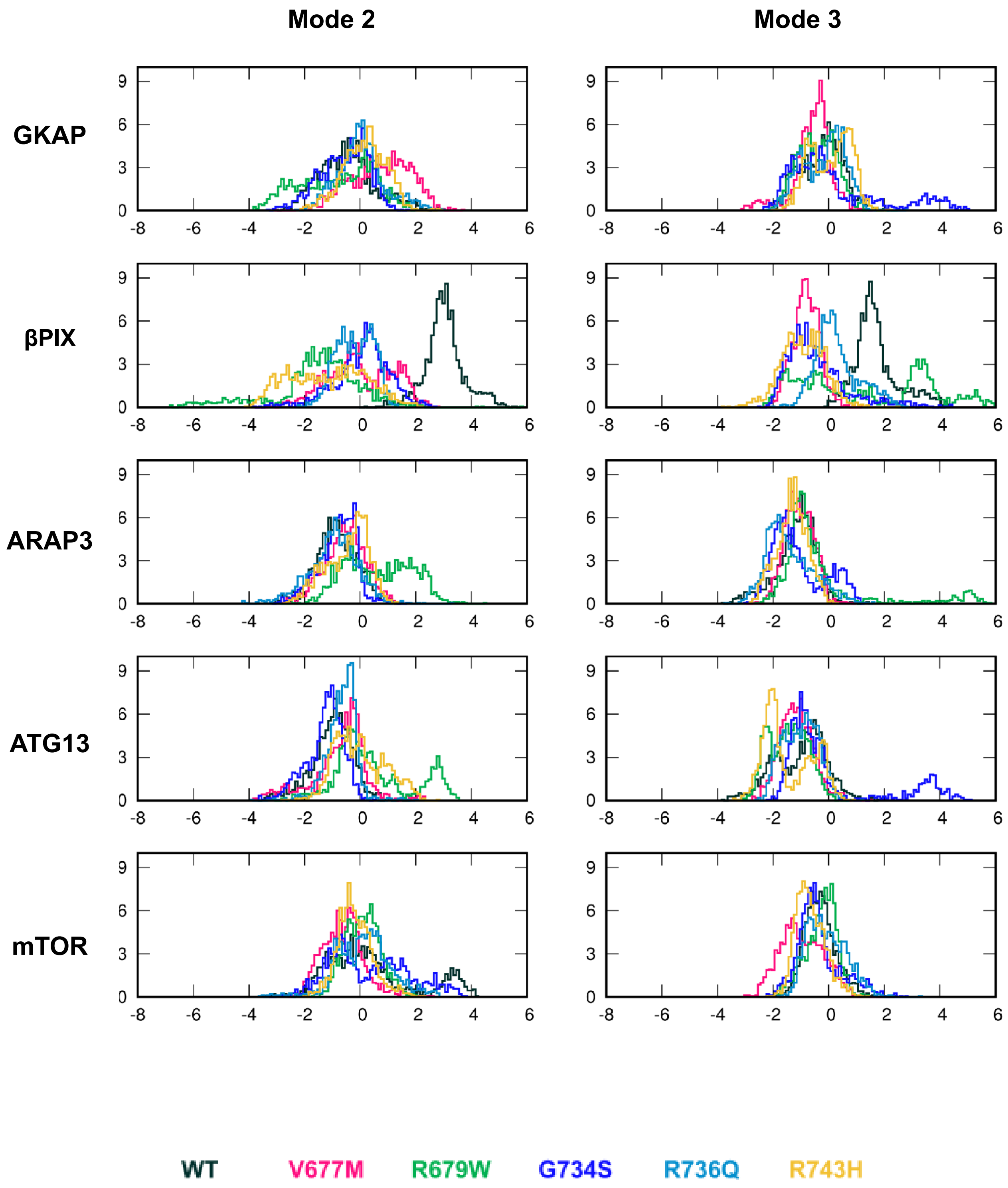

#### J. PCA figures, mode 3-2, separated by variant, colored by peptide

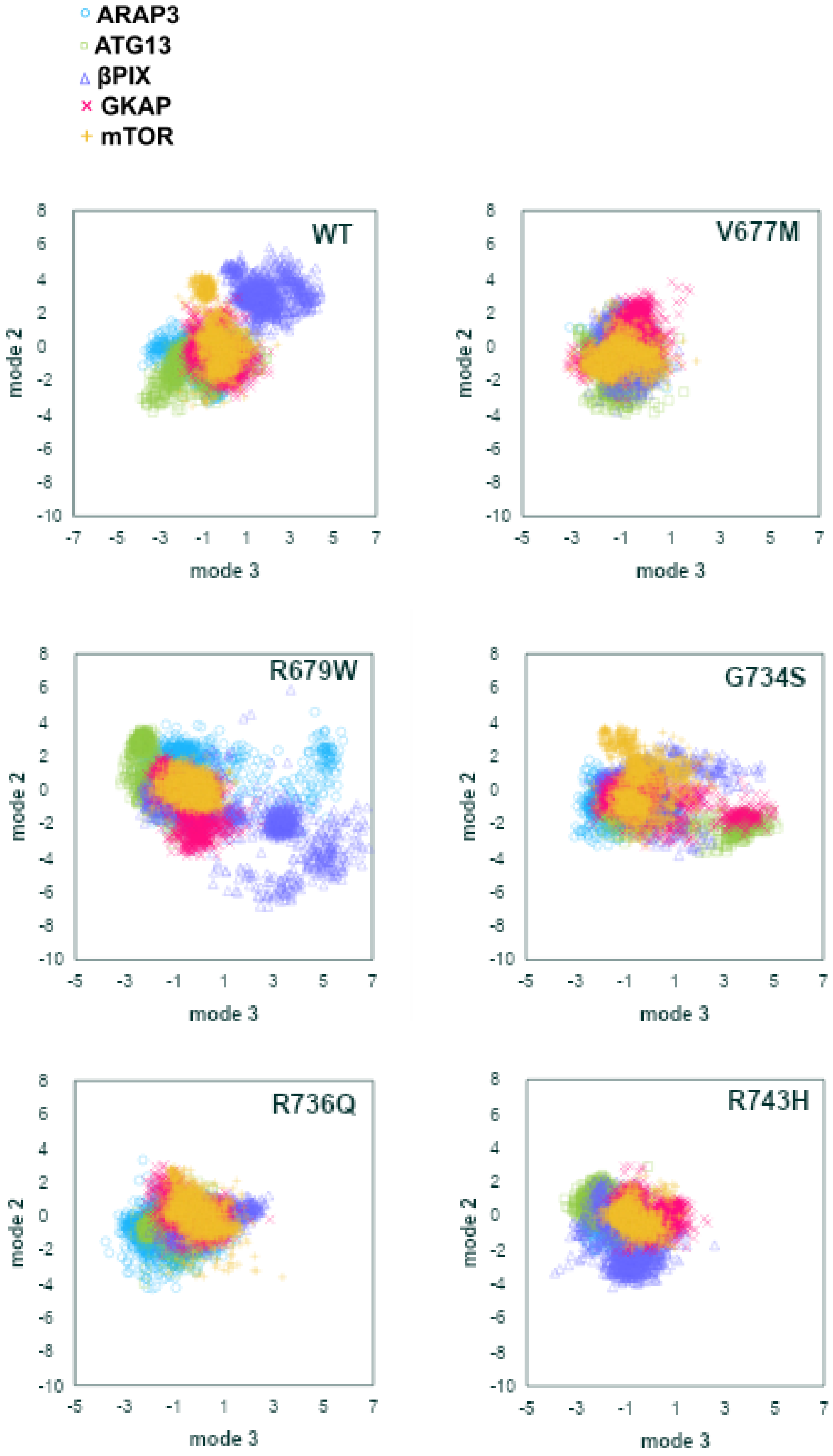

#### K. Histograms of PCA mode 3 and mode 2, separated by variant, colored by peptide

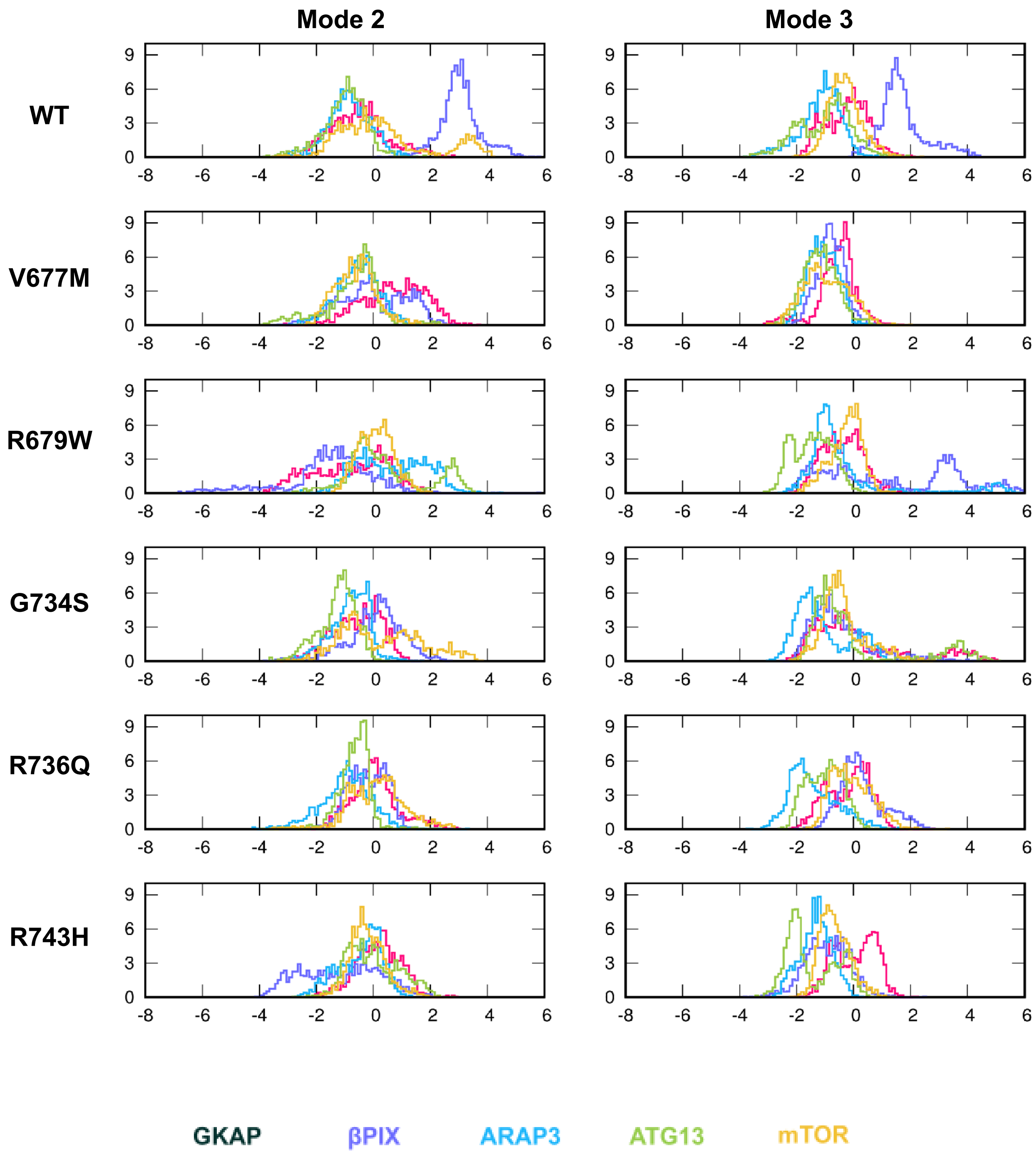

**Figure S5.**

Dynamics of the β2-β3 loop in the MD simulations. **(A)** RMSF values calculated for the loop (residues 680-700). **(B)** Summary of helical propensities assigned to the 680-690 by DSSPCont (sum of G, H, I and T annotations). Mode 1, 2 and 3 are plotted against each other in **(D)**, **(F)**, **(H)** and **(J)** separated into individual plots by variant and peptide. Histograms showing distributions in mode occupancies are included in **(E)**, **(G)**, **(I)** and **(K)**.

#### S6. H735 *X* angle distributions

##### A. GKAP

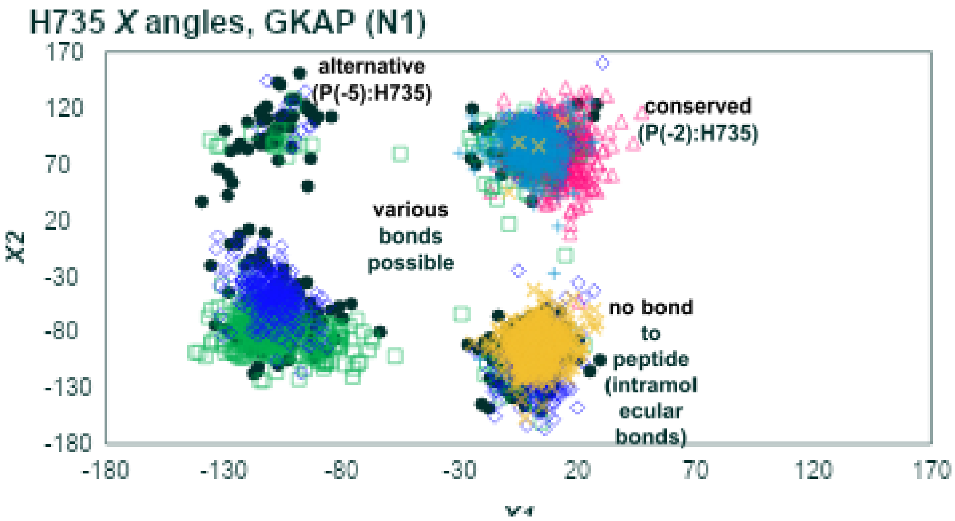

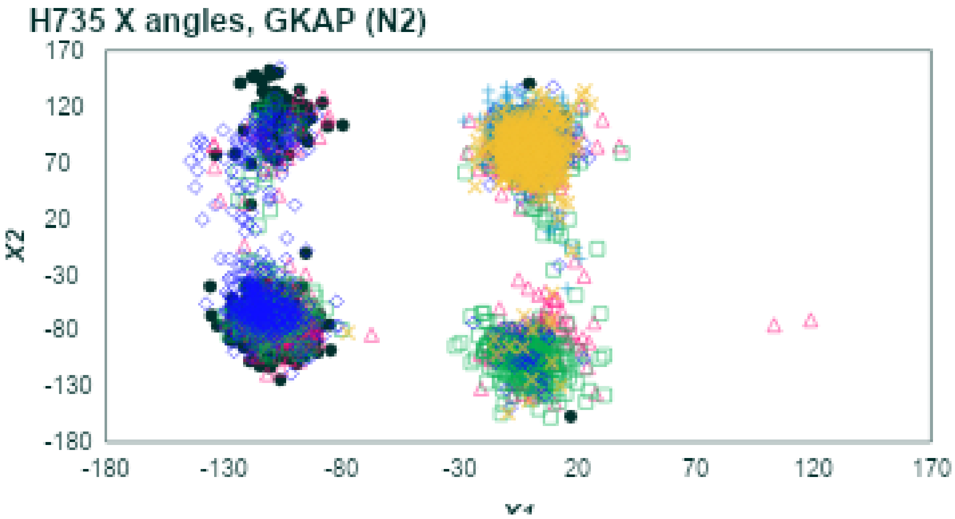

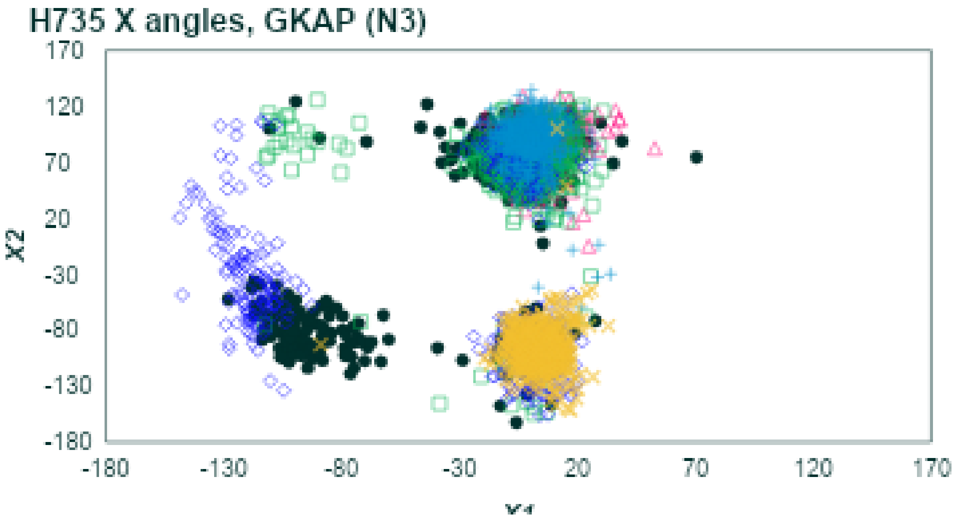

**B. mTOR**

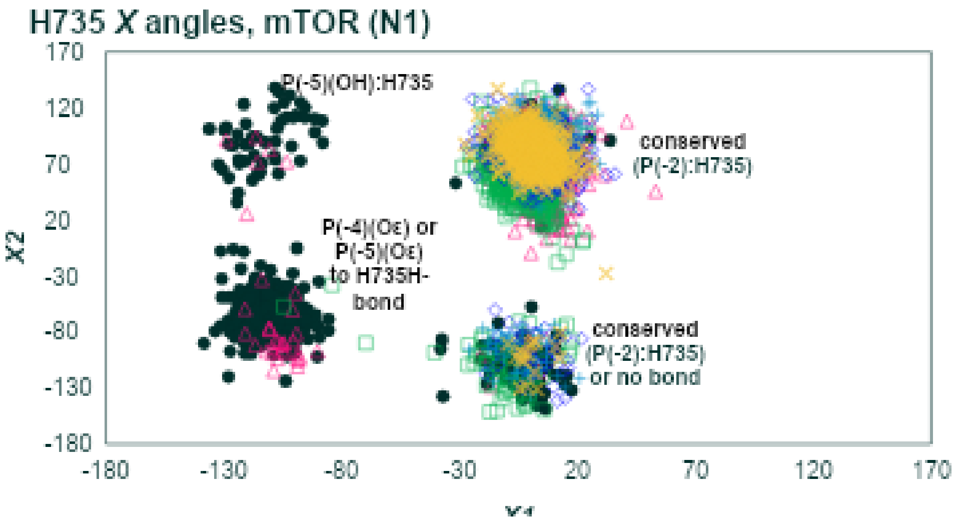
H735 X angles, mTOR (N1)
170
120
70
20
-30
-80
-130
-180
X2
-180
-130
-80
-30
20
70
120
170
P(-5)(OH):H735
conserved (P(-2):H735)
P(-4)(Oε) or P(-5)(Oε) to H735H-bond
conserved (P(-2):H735) or no bond

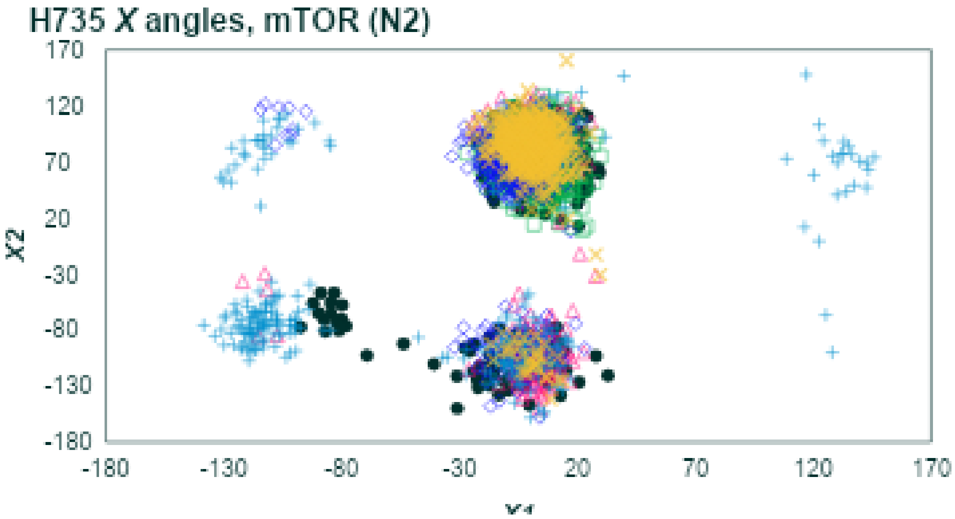
H735 X angles, mTOR (N2)
170
120
70
20
-30
-80
-130
-180
X2
-180
-130
-80
-30
20
70
120
170

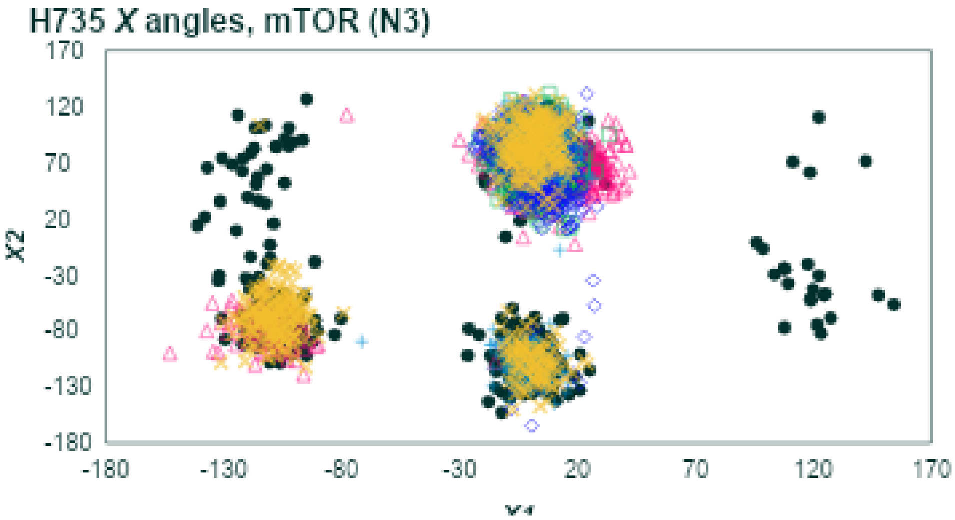
H735 X angles, mTOR (N3)
170
120
70
20
-30
-80
-130
-180
X2
-180
-130
-80
-30
20
70
120
170

#### C. ARAP3

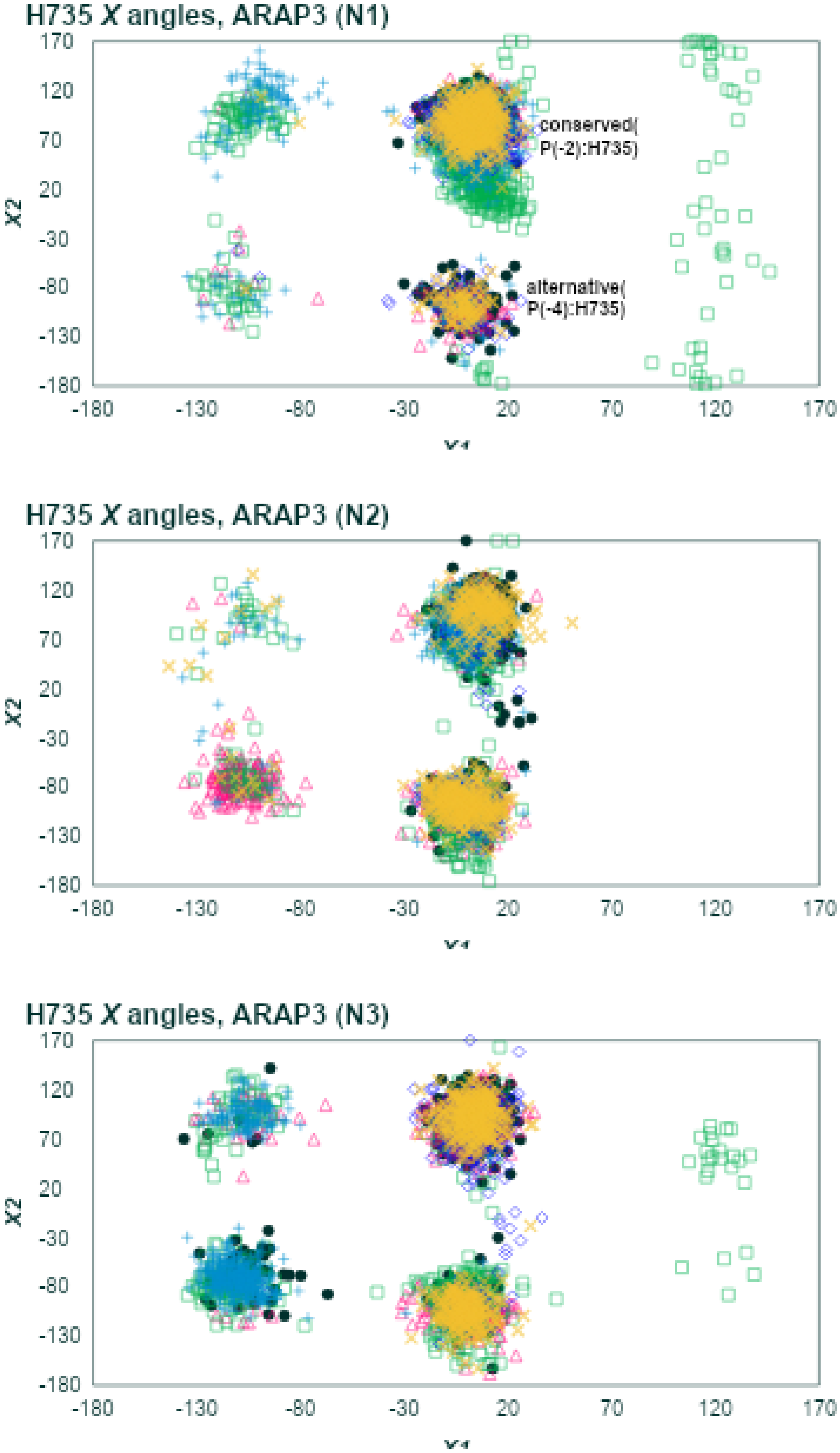

**Figure S6.**

**(A-C)** H735 *X* dihedral distributions for the GKAP, mTOR and ARAP3 peptides respectively. Plots for the first run include descriptions of the hydrogen bonds characteristically occuring in the conformation correspondiing to each cluster.
